# Signals beneath the ectoderm: Mesodermal FGF signaling keeps the dorsally closing epithelium in shape

**DOI:** 10.64898/2026.11.05.764722

**Authors:** Mina Amini, Hellen Y. Chiou, Richard D. Mortensen, Stephanie M. Fogerson, Melissa Sican, Katrina Focht, Khang Huynh, Janice M. Crawford, Daniel P. Kiehart

## Abstract

Morphogenesis requires coordinated biochemical signaling, actomyosin contractility and dynamic adhesion within and between tissues. During *Drosophila* development, ectodermal fibroblast growth factor (FGF) ligands regulate mesoderm morphogenesis through the FGF receptor (FGFR) Heartless. We find that the mesoderm reciprocally influences ectodermal morphogenesis. We show that loss of the FGF ligands Pyramus or Thisbe, Heartless, or its adaptor Stumps transforms normally straight dorsal seams and the formed dorsal epidermis into highly distorted, dynamically changing zigzags during dorsal closure. In these mutants, mesoderm is specified but defective. Mesodermal expression of Heartless restores epithelial stability in *heartless* mutants, whereas ectodermal expression does not. Genetic ablation of mesoderm produces more extensive seam distortions that persist beyond the end of closure, thereby displacing the dorsal midline. Despite these striking deformations, closure continues to completion. Together, our findings reveal a non-cell-autonomous requirement for mesodermal FGFR signaling in stabilizing the closing ectoderm and its newly formed seams. They identify a reciprocal influence of mesoderm on the ectoderm that patterns it and show that stable tissue geometry can be uncoupled from successful completion of a morphogenetic movement.

## Introduction

Morphogenesis coordinates signaling and mechanical interactions among tissues (Tan and Chan, 2026). Actomyosin contractility changes cell shape; dynamic cell–cell and cell–matrix adhesions transmit forces and permit remodeling (Clarke and Martin, 2021; Fernandez-Gonzalez and Harris, 2023; Campàs et al., 2024). Signals exchanged across tissue boundaries along with cell fate and migration regulate these processes. Their coordination within and between tissues underlies stable tissue and embryo behavior.

*Drosophila* dorsal closure exemplifies this coordination (reviewed in Harden, 2002; Kiehart et al., 2017; Hayes and Solon, 2017). Two lateral epithelial sheets advance over the amnioserosa toward the dorsal midline, zipping together at anterior and posterior canthi. Mechanically coupled contributions include amnioserosa contraction and ingression, contractile, actomyosin-rich supracellular purse strings at the leading edges of the lateral epithelium, zipping at the canthi and opposing forces generated within the lateral epithelium (Kiehart et al., 2000; Hutson et al., 2003; Peralta et al., 2007; Toyama et al., 2008). Genetic perturbation, laser microsurgery and modeling show dynamic balance and partial compensation among these contributions, permitting closure despite substantial perturbation of individual force-producing processes (Hutson et al., 2003; Franke et al., 2005; Peralta et al., 2007; Gorfinkiel et al., 2009; Solon et al., 2009; Kiehart et al., 2017).

Mesoderm develops with the ectoderm and moves beneath it during dorsal closure. After mesoderm specification and invagination at stage 6, the fibroblast growth factor (FGF) ligands Pyramus (Pyr) and Thisbe (Ths) from the ectoderm activate the FGF receptor (FGFR) Heartless (Htl) in the mesoderm, where signaling requires the FGF-specific adaptor Stumps (a.k.a. Downstream of FGF/Dof; Gisselbrecht et al., 1996; Michelson et al., 1998; Vincent et al., 1998; Stathopoulos et al., 2004; Wilson et al., 2005; Kadam et al., 2009; McMahon et al., 2010; reviewed in Sun et al., 2020). Htl signaling promotes collapse of the invaginated mesodermal tube onto the ectoderm, establishing tissue contact and a smooth mesodermal monolayer by stage 10 (Wilson et al., 2005; McMahon et al., 2010). Dorsolaterally spread and correctly positioned mesodermal cells receive inductive signals including those from Decapentaplegic (Dpp), that contribute to specification of cardiac and somatic mesoderm (Staehling-Hampton et al., 1994; Frasch, 1995; Stathopoulos et al., 2004; Sun et al., 2020). Htl disruption therefore affects mesoderm organization and multiple differentiated lineages (Beiman et al., 1996; Shishido et al., 1997). Somatic muscles (this study) and cardiac tubes (Haack et al., 2014; Balaghi et al., 2023) move toward the embryo’s dorsal midline during dorsal closure in the overlying ectoderm at stages 13-15. How mesoderm influences ectodermal closure remains unclear.

Here, we show that loss of two FGF ligands produces pronounced, dynamically changing, zigzag seams, instability in the surrounding epidermis and perturbs advancing epithelial edges. Mutations in the FGF tyrosine kinase receptor *htl* and its downstream effector *stumps* phenocopy the loss of the FGF ligands. Tissue-specific experiments confirm that Htl functions in the mesoderm. Two-color, live imaging highlights disrupted dorsal mesodermal organization and captures abnormal mesoderm dynamics in *htl* mutants. *twist* (*twi*) and *snail* (*sna*) mutants, which fail to establish mesoderm, develop more extensive epithelial instability. Laser ablation demonstrates altered mechanical responses in FGF/FGFR-pathway mutants lacking Pyr or Htl. Our findings uncover a reciprocal consequence of ectoderm-to-mesoderm FGF signaling: the mesoderm contributes to the stability of the overlying ectoderm, and FGF signaling in the mesoderm is required for this non-cell-autonomous function during dorsal closure.

## Results

### *Df(2R)ED2247* reveals amnioserosa defects and dynamic epithelial instability during dorsal closure

A forward genetic screen for loci required for dorsal closure using chromosomal deficiencies (*Dfs*), which delete defined genomic intervals that contain numerous, contiguous genes, previously identified abnormalities in embryos homozygous for *Df(2R)ED2247* (hereafter *Df(2R)22*; Mortensen et al., 2018; genomic interval shown in FIG. S3). We performed high-resolution live imaging of *Df(2R)22* embryos using GFP-tagged E-cadherin (E-cad-GFP) to visualize adherens junctions and GFP-tagged Spaghetti squash, the non-muscle myosin II regulatory light chain (Sqh-GFP-RLC; Karess et al., 1991; Royou et al., 2002, 2004), to visualize actomyosin-rich epithelial structures.

E-cad-GFP imaging showed that control amnioserosa retained junctional continuity and contact with the advancing epithelium as the opening narrowed (FIG. S1A; Movie 1). In contrast, *Df(2R)22* embryos developed variable amnioserosa defects (Figs S1B,C, S2B,C). The more frequently observed phenotype involved local separation of peripheral amnioserosa (PAS) from dorsal-most epithelial (DME) cells, disrupted junctional continuity and disintegration of individual amnioserosa cells, particularly later in closure (FIG. S2C^i^–C^ii^, cyan arrowheads; C^ii^, yellow arrowheads; C^iii^–C^iv^, magenta arrowheads, respectively; Movie 2). More extensive PAS–DME separation and progressive tissue disintegration developed earlier in a subset of embryos, leaving cellular debris in the dorsal opening (FIG. S1B^iii^–B^viii^; Movie 3). These differences in timing and severity establish a range of defects in amnioserosa-integrity that accompany the epithelial-instability phenotype described below.

Sqh-GFP-RLC imaging revealed a previously unrecognized phenotype: dynamic epithelial distortions in the lateral and formed dorsal epithelium (Fig. 1). Controls formed a nearly straight dorsal seam as zipping advanced from the two canthi (Fig. 1A^i^–A^vi^; Movie 4). *Df(2R)22* embryos instead developed multiple, changing displacements that distorted the seam into zigzags (Fig. 1B^ii^–B^xi^; Movie 5). Colored arrowheads follow individual distortions on both the anterior and posterior sides of the opening. Distortions arose at different positions and times and evolved independently or concurrently within already fused dorsal epithelium. For example, the cyan-marked posterior distortion is first visible near the canthus in the displayed sequence and becomes strongly tapered over the next 90 min (Fig. 1B^ii^–B^iv^). It partly subsides within 4 min and increases in magnitude again over the next 26 min (Fig. 1B^v^–B^vi^). Over the following 2 h 10 min, it changes in magnitude and orientation, becoming nearly parallel to the midline before largely relaxing—about 4 h 10 min after its first appearance in the sequence (Fig. 1B^vii^–B^xi^). Other regions follow distinct trajectories rather than a uniform seam displacement. Matching asterisks mark residual positions as distortions subside near completion (Fig. 1B^xii^). White arrows in the closer views mark locally twisted epithelial geometry and closely clustered myosin-labelled cell boundaries (Fig. 1C^vii^,C^ix^). Red arrows show progressive PAS–DME separation while the surrounding epithelium remains distorted (Fig. 1B^vi^–B^x^). Thus, epithelial instability is characterized by continuously changing local geometry in the fused dorsal epithelium and along the advancing epithelial edges that shape the dorsal opening.

**Figure 1.**
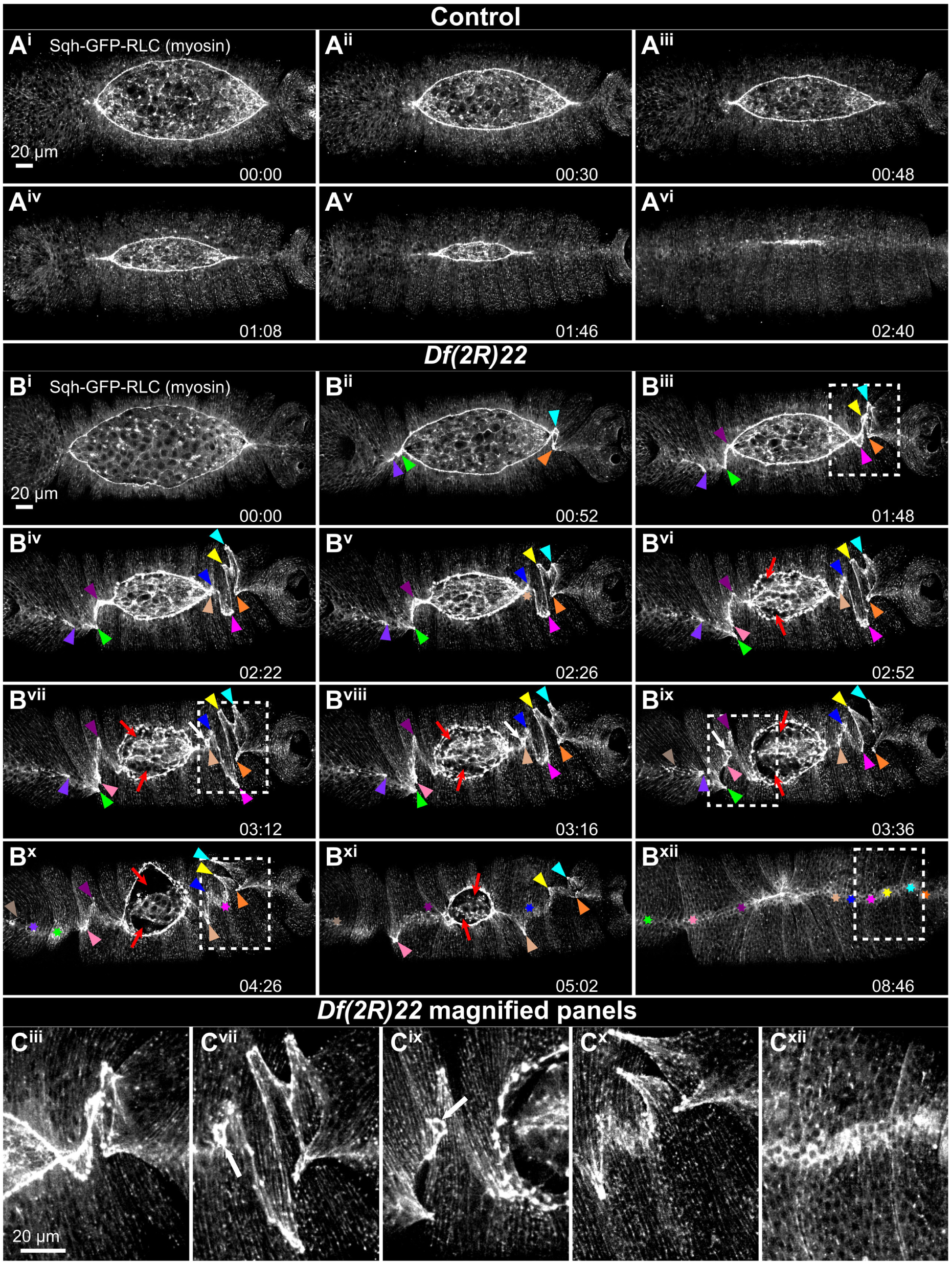
*Df(2R)22* causes dynamic dorsal epithelial distortions. Sqh-GFP-RLC labels non-muscle myosin II in control (A^i^–A^vi^) and *Df(2R)22* (B^i^–B^xii^) embryos. Colored arrowheads follow individual distortions; color matched asterisks mark positions after relaxation. Red arrows indicate PAS–DME separation. Dashed boxes identify regions enlarged in C^iii^, C^vii^, C^ix^, C^x^ and C^xii^ from the corresponding B panels; white arrows highlight complex local configurations. Times, elapsed h:min from each series’ first frame. PAS, peripheral amnioserosa; DME, dorsal-most epithelial cells. Scale bars, 20 *μ*m.

The distorted epithelial phenotype was detectable with both myosin- and cadherin-based reporters, though it was more readily resolved with cytoskeletal reporters, including sGMCA, a GFP–Moesin actin-binding-domain reporter driven by the *spaghetti squash* promoter. We scored three categories—indistinguishable from controls (absent), subtle but observable, or prominent distortions—to quantify penetrance (the fraction affected) and qualitative expressivity among affected embryos. Epithelial distortions affected 31/32 myosin-labelled *Df(2R)22* embryos (97%; 22 prominent, in one embryo such distortions were absent) and 22/22 cadherin-labelled embryos analyzed (100%; 14 prominent; FIG. S4A; Table S4, section A). Penetrance and expressivity were assessed within each reporter background (FIG. S4). Cadherin-based reporters combined with lower-resolution imaging help explain why earlier studies revealed amnioserosa defects but did not resolve the less conspicuous epithelial instability and zigzag seams.

### Genetic dissection of *Df(2R)ED2247* identifies the FGF ligands *pyr* and *ths* as regulators of epithelial stability

Overlapping deficiencies, in trans to the parent *Df(2R)ED2247*, separated epithelial instability from amnioserosa disintegration: *Df(2R)22B* and laboratory-generated *Df(2R)22F* retained epithelial distortions without amnioserosa disintegration, whereas *Df(2R)22C* retained both phenotypes. Their overlaps highlighted the *pyr / ths*-containing interval and a distinct region associated with amnioserosa disintegration (FIG. S3; Table S1; Materials and Methods).

Deleting either FGF ligand independently reproduced epithelial instability. *Df(2R)pyr^36^* and *Df(2R)ths^238^* remove the respective transcription units (Kadam et al., 2009) and each produced dorsal epithelial distortions in trans to *Df(2R)22* (Fig. 2A, Movie 6, for *pyr*; Fig. 2B, Movie 7, for *ths*). Tracking individual distortions in *pyr* mutants showed repeated excursions away from and back toward the dorsal midline, usually on the side where each distortion initially arose (Fig. 2F). In myosin-labelled embryos, distortions affected 22/27 *Df(2R)pyr^36^ / Df(2R)22* embryos (81%; 13 prominent), 9/9 *pyr^18^ / Df(2R)22* embryos (100%; six prominent) and 17/19 *pyr^18^ / Df(2R)pyr^36^* embryos (89%; ten prominent). Corresponding F-actin cohorts showed 6/ 7 affected *Df(2R)pyr^36^ / Df(2R)22* embryos (86%; three prominent) and 20/20 affected *pyr^18^ / Df(2R)22* embryos (100%; four prominent; Fig. 2E; FIG. S4B; Table S4, section B). Both ligands are therefore required for epithelial stability.

**Figure 2.**
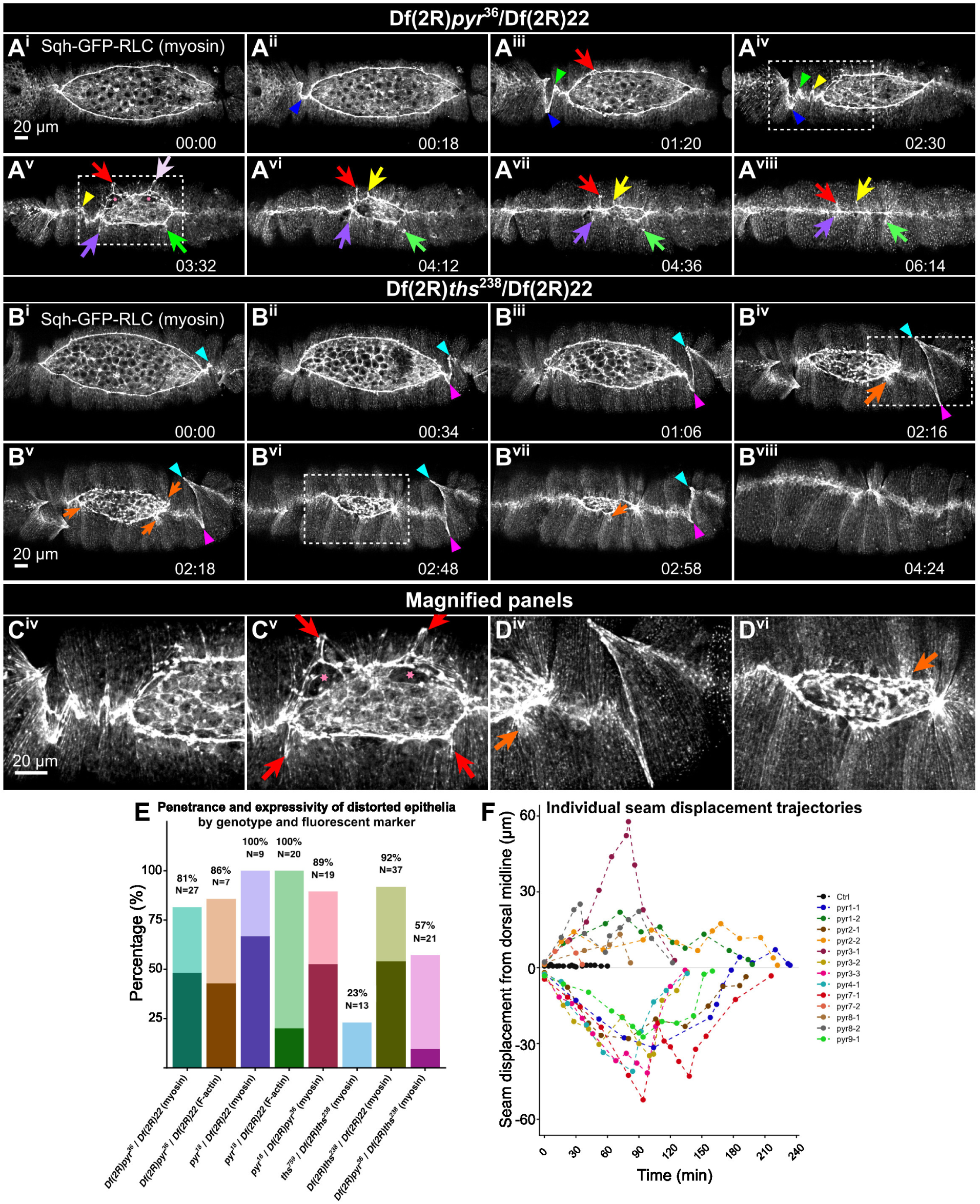
Both FGF ligands are required for epithelial stability during closure. Sqh-GFP-RLC time series show *Df(2R)pyr^36^* / *Df(2R)22* (A^i^–A^viii^) and *Df(2R)ths^238^* / *Df(2R)22* (B^i^–B^viii^). Colored arrowheads track seam distortions; arrows track deformations of purse-string and seam-associated actomyosin structures or mark abnormal local attachments, and pink asterisks indicate transient PAS–DME separation. Magnified panels correspond to A^iv^→C^iv^, A^v^→C^v^, B^iv^→D^iv^ and B^vi^→D^vi^. (E) Penetrance and qualitative expressivity by genotype and reporter; dark and light portions denote prominent and subtle but observable distortions, respectively. Labels show percentages and embryo counts. (F) Individual *pyr* seam-displacement tracks (13 tracks from seven embryos) and a single control embryo reference. Lines connect observations within the 13 displacement tracks; time in F is in minutes. Times, h:min. Scale bars, 20 *μ*m.

The molecularly defined *ths^759^* deletion produced subtle distortions in 3/13 *ths^759^ / Df(2R)ths^238^*embryos (23%), versus 34/37 *Df(2R)22 / Df(2R)ths^238^* embryos (92%; 20 prominent) (Fig. 2E). This suggests residual Ths activity despite deletion of most of the FGF core domain and its original null interpretation (Klingseisen et al., 2009).

Embryos trans-heterozygous for severe alleles *Df(2R)pyr^36^*and *Df(2R)ths^238^*, retaining one wild-type copy of each ligand, showed distortions in 12 of 21 cases (57%). Distortions were generally mild, with only two embryos showing higher expressivity (10%; Fig. 2E). One wild-type copy of each ligand therefore reduced but did not prevent epithelial instability. This phenotype indicates an interaction between *pyr* and *ths*, consistent with second-site non-complementation (see, for example, Halsell and Kiehart, 1998).

Instability affected both the formed dorsal epidermis, the seam it includes and the leading-edge of the lateral epidermis in regions of the embryo that were still dorsally open. Colored arrowheads track seam distortions, whereas colored arrows track changing deformations of purse-string and seam-associated actomyosin structures (Fig. 2A^iii^,A^v^–A^viii^,B^iv^–B^vii^). Transient PAS–DME separations accompanied some distortions (Fig. 2A^v^,C^v^, pink asterisks). Unlike the progressive detachment associated with amnioserosa deterioration in *Df(2R)22* (FIG. S1), these separations narrowed over approximately 24 min in the illustrated *pyr* embryo. Orange arrows mark abnormal local attachments near the canthi or purse strings in *ths* embryos (Fig. 2B^iv^–B^vii^,D^iv^,D^vi^).

Despite these changing local distortions, closure continued, and the distortions largely relaxed as the opening approached closure in both *pyr* and *ths* mutants, though seams in some embryos retained slight residual deviation from straight or gently arced. Mean narrowing rates were 7 ± 2 nm s^-1^ in *pyr* and 9 ± 1 nm s^-1^ in controls (mean ± s.d.; six *pyr* embryos and four controls; Holm-adjusted P = 0.289 across four mutant–control comparisons; Figs S5, S6; Table S3). These measurements did not establish a difference in mean rate between *pyr* and control.

Collectively, these results identify the FGF ligands *pyr* and *ths* as required for epithelial stability during dorsal closure and show that substantial epithelial instability need not prevent closure progression.

### Heartless and Stumps are required for epithelial stability during dorsal closure

We asked whether epithelial stability depends on the FGF receptor and its signaling machinery. Pyr and Ths activate Htl, which requires the adaptor/scaffolding protein Stumps to transduce the signal (Michelson et al., 1998; Vincent et al., 1998; Imam et al., 1999). We therefore examined epithelial behavior in *htl* and *stumps* mutants.

Controls maintained an essentially straight or slightly arced seam (Fig. 3A; Movie 8), whereas *htl^AB42^ / Df(3R)BSC650* embryos developed distortions that changed dramatically during closure and largely subsided by its completion (Fig. 3B; Movie 9). Homozygous *stumps^09904b^* mutants developed similar distortions (Fig. 3C,E^ii^; Movie 10). Epithelial instability was highly penetrant across myosin, F-actin and cadherin reporters in *htl^AB42^ / Df(3R)BSC650* embryos (92–100%; FIG. S7A; Table S4, section C). Homozygous *htl^AB42^* cohorts were affected in 18/22 myosin-labelled embryos (82%) and 28/28 F-actin-labelled embryos (100%); all seven F-actin-labelled *stumps^09904b^* embryos examined were affected (100%).

**Figure 3.**
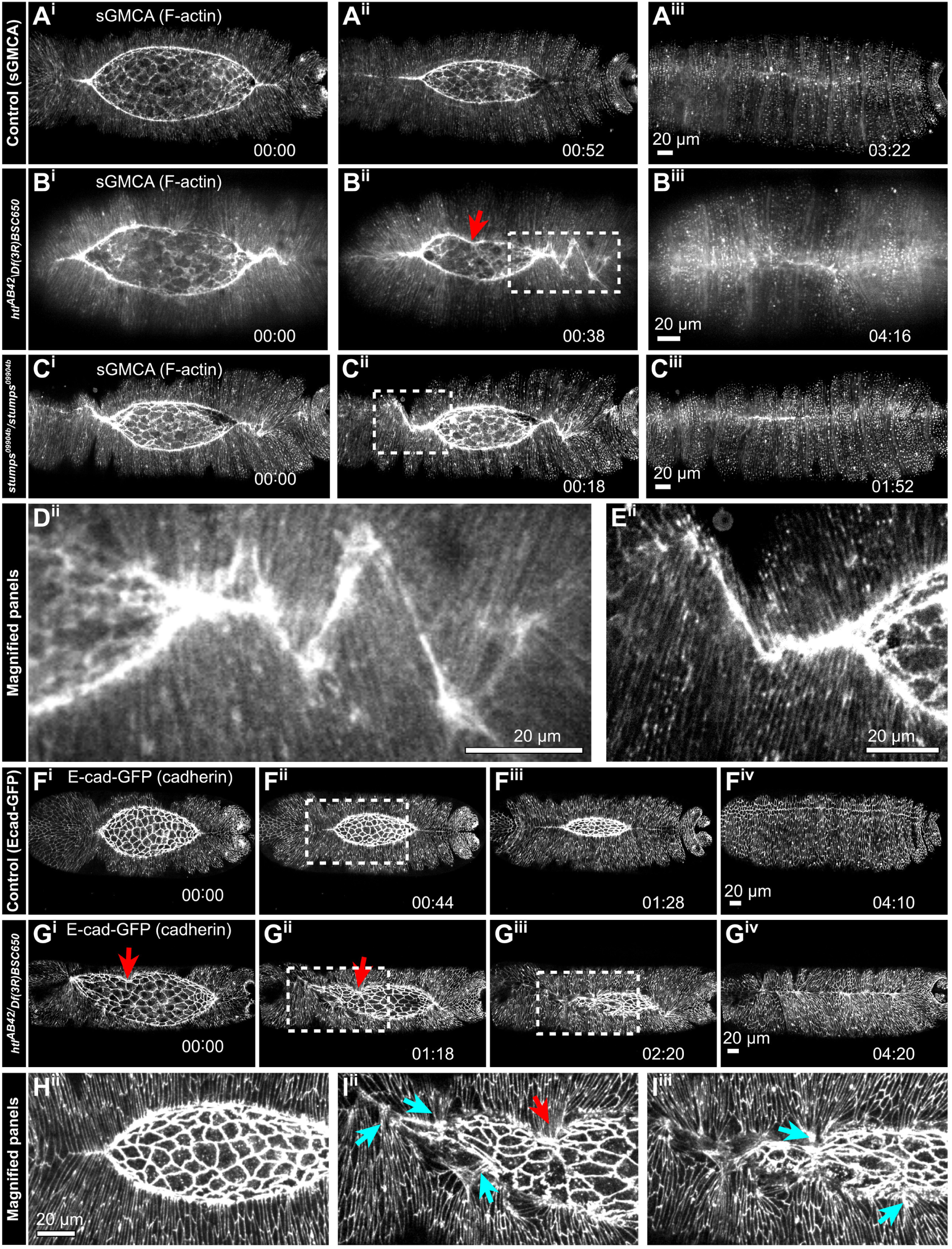
Htl and Stumps are required for stable epithelial geometry. sGMCA-labelled control (A), *htl^AB42^ / Df(3R)BSC650* (B) and homozygous *stumps^09904b^* (C) embryos are shown through closure. Enlargements of B^ii^ and C^ii^ are D^ii^ and E^ii^, respectively. Cadherin-labelled control and *htl* embryos are F and G; H^ii^ enlarges F^ii^, and I^ii^–I^iii^ enlarge G^ii^–G^iii^. Red arrows show local leading-edge kinks; cyan arrows mark abnormal leading-edge cellular associations. Roman indices identify successive frames or their matching magnifications. Times, h:min. Scale bars, 20 *μ*m.

In *htl* mutants, epithelial instability involved the advancing epithelial edge as well as the formed dorsal seam. Uneven advancement of neighboring DME cells kinked the purse strings (Fig. 3B^ii^,G^i^–G^ii^,I^ii^, red arrows). Cadherin imaging further showed disruption of the ordered cellular associations at this edge: in *htl* mutants (Fig. 3G,I^ii^; Movie 11) ectopic DME–DME or DME–amnioserosa associations were often seen (Fig. 3I^ii^–I^iii^, cyan arrows; Movie 11). These defects extend epithelial instability across the formed dorsal epithelium and advancing lateral epithelial sheets, as in ligand mutants (Fig. 2). Mean closure rate in *htl* was 7 ± 1 nm s^-1^ (mean ± s.d.; six embryos), without a significant difference from controls (Holm-adjusted P = 0.281; FIG. S7B; Table S3). Htl was imaged with an F-actin reporter, whereas the rate-control cohort used Sqh.

Together, ligand, receptor and adaptor mutants establish that epithelial stability during dorsal closure requires Pyr/Ths-Htl-Stumps signaling.

### Mesodermal Heartless is sufficient to restore epithelial stability during dorsal closure

Htl localized to mesodermal derivatives beneath the closing epithelium, suggesting that its contribution to epithelial stability is non-cell-autonomous. Live imaging of Htl-mCherry, expressed under its own promoter/enhancer cassette (Irizarry and Stathopoulos, 2015) together with E-cadherin-GFP, detected Htl in underlying mesoderm rather than in the closing epithelium (Fig. 4A–C; Movie 12), consistent with its established mesodermal expression (Beiman et al., 1996; Shishido et al., 1997). Htl-mCherry-positive populations redistributed dorsally as the epithelial sheets advanced (Fig. 4A^i^–C^iv^). Later, Htl was prominent in cardiogenic mesoderm, including developing cardiac tubes (Fig. 4B^iii^–B^iv^,E^iii^, yellow arrowheads), and somatic muscles (Fig. 4B^iii^,E^iii^, cyan arrowheads). The orange bracket marks gut-associated signal, consistent with Htl expression in visceral mesoderm and developing gut musculature (Beiman et al., 1996; Sun et al., 2020).

**Figure 4.**
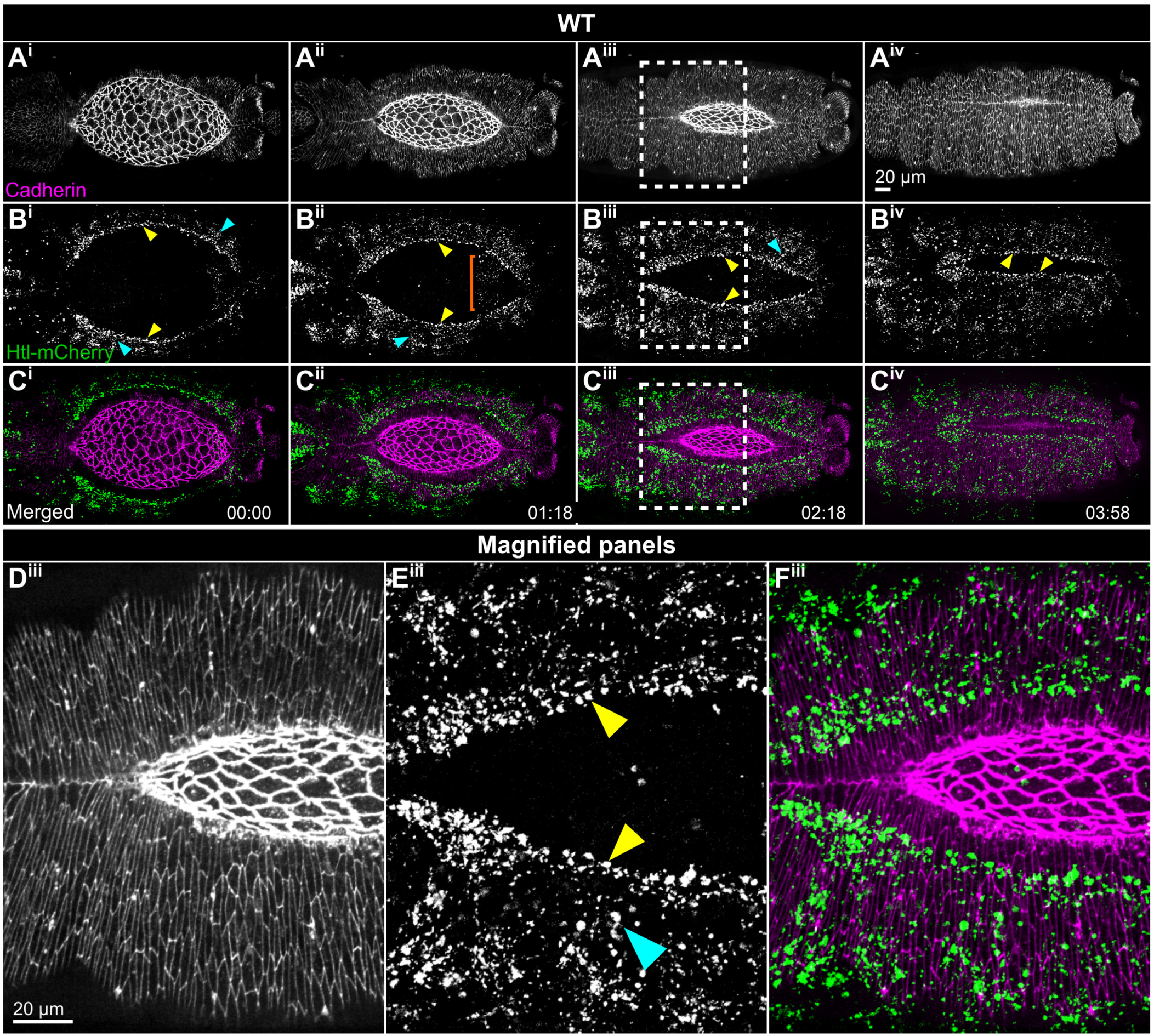
Htl localizes to mesodermal derivatives beneath the closing epithelium. (A–C) Maximum projections of E-cadherin-GFP (gray or magenta), Htl-mCherry (gray or green) and merged channels from one embryo. Yellow arrowheads identify cardiogenic populations; cyan arrowheads identify somatic-mesodermal populations, and the orange bracket in B^ii^ is to the left of gut-associated Htl-mCherry signal, consistent with visceral mesoderm/musculature (only the gut’s uppermost region is projected). Dashed boxes in A^iii^–C^iii^ correspond to D^iii^–F^iii^. Times, h:min from the first displayed frame. Scale bars, 20 *μ*m; separate bars apply to parent and magnified images.

To test whether undetectably low ectodermal Htl might stabilize the lateral and dorsal epithelium, we performed reciprocal tissue-specific expression of wild-type Htl in an *htl* mutant background. We drove *UAS-htl^WT^*in either the mesoderm with *twi-GAL4* or the ectoderm with *69B-GAL4* (Brand and Perrimon, 1993; Fig. 5). Because the *twi-GAL4* rescue-control background differed phenotypically from the *htl* mutant combinations described above, we first characterized these controls directly. Matched *twi-GAL4 > no UAS responder; htl^AB42^ / htl^AB42^*controls retained distortions in 9/9 F-actin embryos (100%; six subtle, three prominent) and 15/16 cadherin embryos (94%; Fig. 5A,B; Movie 13; Table S4, section D). Distortions were generally less pronounced than in the *htl* combinations analyzed previously, and their timing varied: in some embryos, they appeared only late in closure. Local pull-like deformations along the purse strings and the seams of the formed dorsal ectoderm were also evident, including in embryos with relatively modest distortions (Fig. 5B).

**Figure 5.**
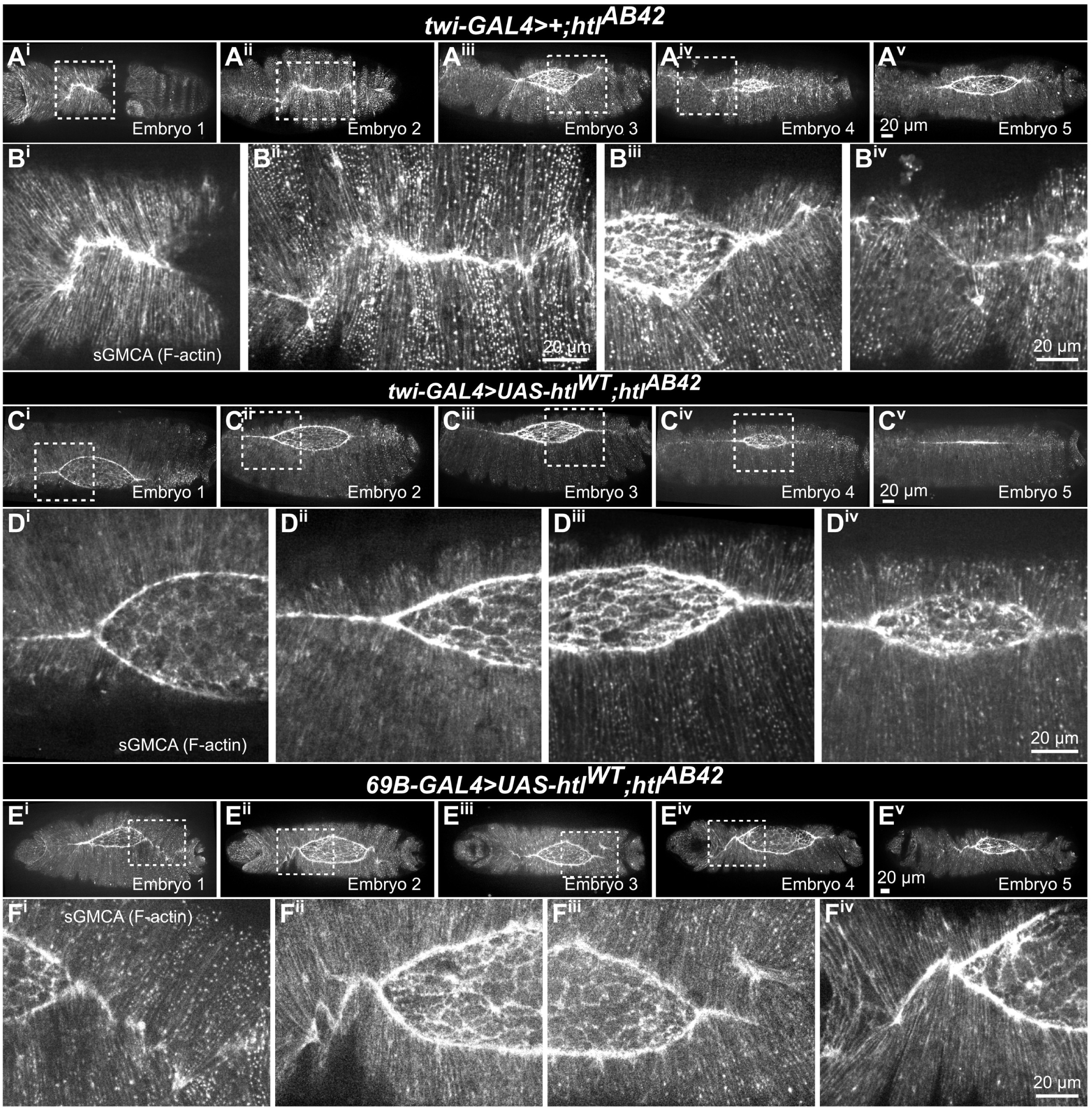
Mesodermal Htl expression restores epithelial stability. sGMCA images show matched *twi-GAL4 > no UAS responder; htl^AB42^ / htl^AB42^* “controls” (A); Magnifications of A^i^-A^iv^ (B^i^–B^iv^); mesodermal rescue, *twi-GAL4 > UAS-htl^WT^; htl^AB42^ / htl^AB42^* (C); Magnifications of C^i^-C^iv^ (D^i^-D^iv^); ectodermal expression, *69B-GAL4 > UAS-htl^WT^; htl^AB42^ / htl^AB42^* (E); and Magnifications of E^i^-E^iv^ (F^i^-F^iv^). Five representative embryos are shown per condition. Dashed boxes identify the enlarged regions. Images compare phenotypes rather than identical developmental time points. Scale bars, 20 *μ*m.

Restoring *htl* in the mesoderm via *twi-GAL4 > UAS-htl^WT^; htl^AB42^ / htl^AB42^* suppressed the distorted seam phenotype. Dorsal epithelial distortions were absent from all nine F-actin embryos (0/9) and all 16 cadherin embryos (0/16) scored (Fig. 5C; Movie 14; Table S4, section D). Five representative embryos are shown. The dorsal seam remained comparatively straight as closure progressed, although with subtle differences in morphology, including a somewhat elongated seam in embryo 4 (Fig. 5D^iv^). Mesodermal Htl therefore eliminated characteristic distortions in the matched *htl* mutant control background.

By contrast, ectodermal expression of *htl* did not eliminate epithelial instability. Dorsal epithelial distortions remained evident in 19/22 *69B-GAL4 > UAS-htl^WT^; htl^AB42^ / htl^AB42^* F-actin embryos (86%; seven subtle, 12 prominent), ranging from subtle to more pronounced configurations (Fig. 5E,F; Movie 15; Table S4, section D).

We evaluated *htl* expression in these backgrounds using hybridization chain reaction fluorescence in situ hybridization (HCR-FISH) for *htl* together with odd-skipped (ectodermal) and Mef2 (mesodermal) as spatial markers (FIG. S9C^ii^,E^ii^,F^ii^). In the ectodermal-expression genotype, *htl* transcript was detected in the ectoderm, confirming expression of the *UAS-htl^WT^* transgene in the tissue targeted by *69B-GAL4*. Thus, epithelial distortions in *69B-GAL4* driven *htl^WT^* occur despite ectodermal *htl* expression.

Together, these results indicate that mesodermal Htl activity contributes to overlying epithelial stability during dorsal closure (FIG. S10).

### How does loss of FGF signaling alter the relationship between mesoderm and ectoderm during dorsal closure?

Given Htl’s mesodermal site of action, we examined mesodermal organization and movement relative to the closing epithelium. We followed both tissues simultaneously using E-cadherin to mark ectodermal junctions and *twi-GAL4*-driven *UAS-RFP-Moesin* to visualize mesodermal F-actin-associated structures (Fig. 6A–C; Movie 16).

**Figure 6.**
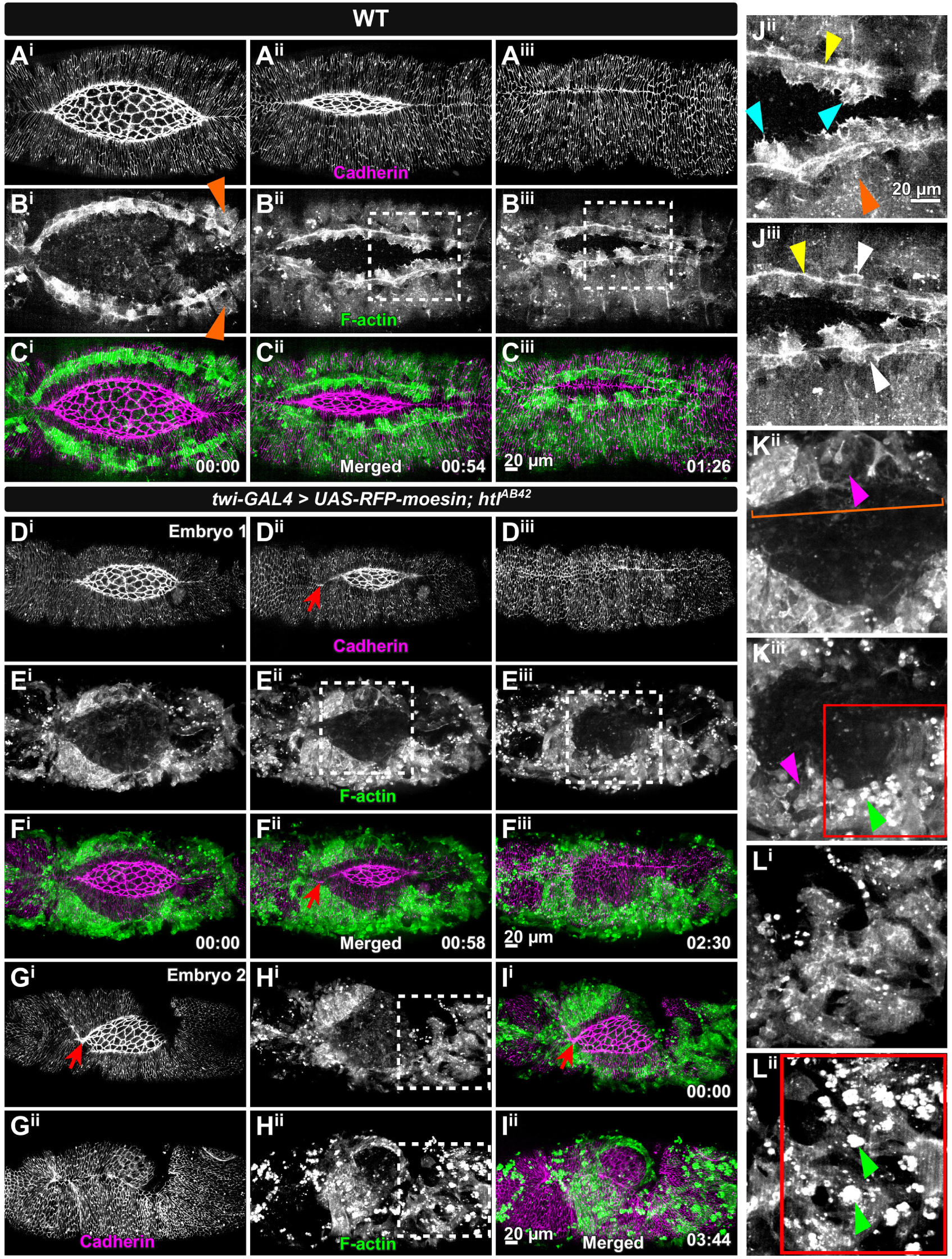
Htl loss disrupts mesodermal organization during dorsal movement. Control junctional (E-cadherin, gray, A), mesodermal (*twi-GAL4*-driven RFP-Moesin-actin-binding domain, B) and merged channels (E-cadherin, magenta, RFP-moesin-actin-binding domain in green, C). Two *htl* mutants: D–F and G–I. RFP-labelled F-actin is shown in green. Control magnifications J^ii^–J^iii^ correspond to B^ii^–B^iii^; cyan arrowheads identify protrusive cardiac cells; yellow arrowheads mark the mesodermal F-actin-rich cable at their trailing edge, distinct from the dorsal-opening purse string; white arrowheads identify pericardial cells farther ventral to this cable. Orange arrowheads in B^i^ mark somatic-muscle organization. Mutant magnifications K^ii^–K^iii^ correspond to E^ii^–E^iii^, and L^i^–L^ii^ to H^i^–H^ii^. The orange bracket in K^ii^ marks disrupted dorsal mesodermal architecture, magenta arrowheads indicate fragmentation, and green arrowheads/red boxes show reporter-positive aggregates. Red arrows indicate epithelial distortions. Times, h:min. Scale bars, 20 *μ*m. WT elapsed times are rounded to the nearest minute.

In controls, bilateral cardiogenic rows lay behind the advancing epithelial edge. Cardioblasts extended dorsal protrusions (Fig. 6B^ii^–B^iii^,J^ii^, cyan arrowheads; Haack et al., 2014; Balaghi et al., 2023). An F-actin-rich mesodermal cable lay at the trailing, ventral edge of the protrusive cardiac cells (Fig. 6J^ii^–J^iii^, yellow arrowheads), consistent with the cardioblast–pericardial interface cable described by Balaghi et al. (2023). Pericardial cells lay ventrally behind the cable (Fig. 6J^iii^, white arrowheads). More ventral, segmentally organized somatic muscles (Fig. 6B^i^, orange arrowheads) moved dorsally with the epithelium and cardiogenic mesoderm (this study). Their coordinated progression was tracked relative to the cadherin-labelled epithelium.

Cardiac protrusions approached the dorsal-opening edge as closure progressed. In the first and second frames (Fig. 6C^i^–C^ii^), their image-plane separations from that edge were approximately 14–25 *μ*m and 8–10 *μ*m, respectively; corresponding separations of the trailing mesodermal cable were 46–50 *μ*m and 21–28 *μ*m. In the third displayed frame, after ectodermal closure, protrusive margins lay approximately 5–20 *μ*m from the formed seam and approximately 15–35 *μ*m from the cable (Fig. 6C^iii^).

This architecture was profoundly disrupted in *htl* mutants. Two-color imaging revealed mesodermal material beneath the epithelium but no ordered dorsal cardiogenic rows or regular somatic-muscle organization (Fig. 6E,H; Movie 17). The cadherin channel revealed accompanying epithelial distortions (Fig. 6D^ii^,F^ii^,G^i^,I^i^, red arrows). The characteristic cardiac/pericardial configuration and intervening cable were absent in the illustrated mutant region (Fig. 6K^ii^, orange bracket).

Despite its disorganization, *htl* mutant mesoderm continued moving dorsally during closure (Fig. 6D–I). As closure progressed, regions of the mesoderm lost integrity and fragmented (Fig. 6K^ii^–K^iii^, magenta arrowheads), while RFP-Moesin-positive aggregates and cellular debris became prominent (Fig. 6K^iii^,L^ii^, green arrowheads and red boxes). Mesodermal disruption and overlying epithelial abnormalities did not correspond precisely in space.

### Complete loss of mesoderm exacerbates ectodermal epithelial instability

Genetic ablation of mesoderm exacerbated epithelial instability, distinguishing complete mesodermal tissue loss from the disorganization caused by FGF-pathway disruption. We examined *sna^1^* and *twi^1^*mutants, which fail to establish a distinct, invaginated mesoderm (Leptin, 1991; Reeves and Stathopoulos, 2009; Martin, 2020; Sun et al., 2020).

Both mutants developed conspicuous zigzag distortions of the forming dorsal epithelium (Fig. 7A, Movie 18, for *sna*; Fig. 7B,C, Movie 19, for *twi*). In myosin-labelled embryos, *sna^1^* embryos were affected in 12/12 cases (100%) and *twi^1^* in 10/11 (91%). F-actin-labelled embryos showed 10/11 affected *sna^1^* embryos (91%) and 11/11 *twi^1^*embryos (100%; FIG. S11A; Table S4, section E).

**Figure 7.**
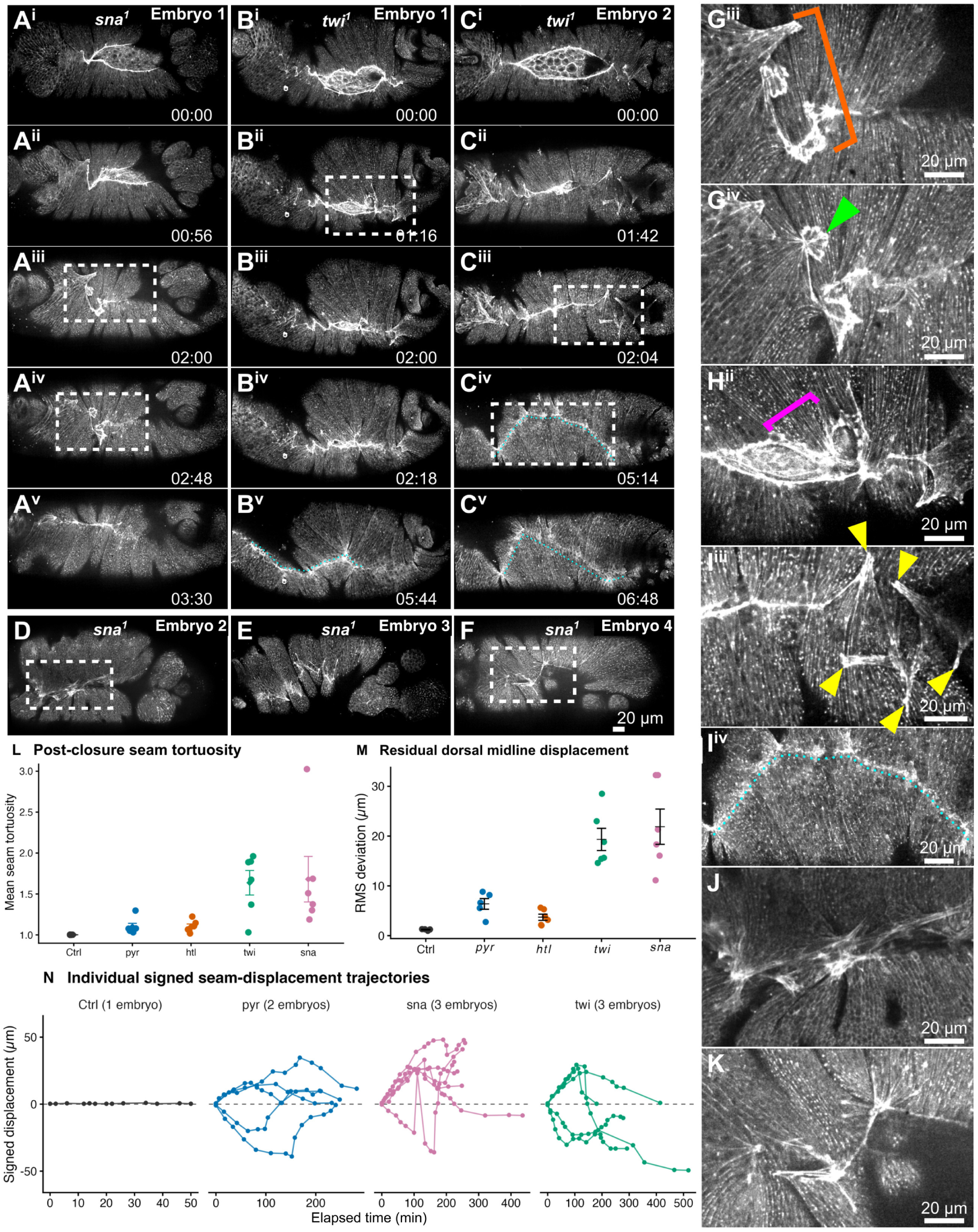
Local distortions and residual midline deviations persist past completion of closure in mesoderm-minus mutants. Sqh-GFP-RLC images show *sna^1^* (A,D–F) and *twi^1^* (B,C) embryos. Magnifications: G^iii^–G^iv^ corresponds to A^iii^–A^iv^, H^ii^ to B^ii^, I^iii^–I^iv^ to C^iii^–C^iv^, J to E and K to F. Orange bracket, persistent distortions (G^iii^). Green arrowhead, cell boundaries meeting in a small distorted region (G^iv^). Magenta bracket, purse-string distortion and uneven DME advancement (H^ii^). Yellow arrowheads (I^iii^), mark oblique or nearly horizontal distortions. Cyan dotted trace shows midline deviation (I^iv^). (L) Mean seam tortuosity across two frames per embryo (10 and 20 min post closure); n=28 (control 4; each mutant 6). Points represent embryos; bars show mean ± s.e.m. *twi* / *sna* versus *pyr* / *htl*: 12 embryos per class (Welch P=0.00355). (M) Raw RMS deviation after local zigzag relaxation; n=27 (control 4, *pyr* 5, others 6 each), mean ± s.e.m. (N) Signed displacement trajectories before closure, relative to a fixed dorsal midline axis. Lines connect observations within a track (FIG. S13). Times in images, h:min; scale bars, 20 *μ*m.

Zigzag distortions remained conspicuous near or after closure in *twi* and *sna* (Fig. 7A^iii^–A^iv^,G^iii^, orange bracket; B^v^,C^iv^–C^v^,I^iv^), whereas they largely relaxed by the time closure was complete in *pyr* and *htl* (Fig. 2A^v^–A^viii^,B^iv^–B^vii^; Fig. 3A,B). They could run obliquely or nearly parallel to the dorsal midline (Fig. 7I^iii^, yellow arrowheads) and persisted alongside curvature of the overall dorsal midline (Fig. 7B^v^,C^iv^–C^v^,I^iv^, cyan dotted traces). Some displaced seam regions crossed the reference midline (Fig. 7N; FIG. S13).

In *twi* and *sna* mutants, epithelial instability was also exacerbated in the lateral epithelium. Asynchronous DME advancement more severely kinked the purse strings (Fig. 7H^ii^, magenta bracket). Additionally, several cell boundaries met in small, distorted regions (Fig. 7G^iv^, green arrowhead), such that deformations clustered or folded, producing complex local configurations (Fig. 7D–F,J,K).

Quantitatively, all four mutants had higher peak seam tortuosity than control (adjusted P ≤ 0.0090), but peak severity did not distinguish *twi* / *sna* from *pyr* / *htl* (difference = 0.1 ± 0.3 s.e., P = 0.636; Table S3). Maximum tortuosity and post-closure measurements are shown in FIG. S12.

Seam distortions persisted after closure. At 10 and 20 min after dorsal closure, mean tortuosity was 1.676 and 1.598 in *twi*, 1.679 and 1.683 in *sna*, 1.106 and 1.097 in *pyr*, and 1.107 and 1.099 in *htl*, respectively (six embryos per genotype). The equally weighted *twi* / *sna* genotype mean exceeded *pyr* / *htl* at both times (differences 0.571 and 0.542 at 10 and 20 min, respectively; Holm-adjusted P = 0.00174 each; Table S3), supporting greater persistence in mesoderm-specification mutants. Averaging the two time points within each embryo gave the same conclusion (Welch P = 0.00355; Fig. 7L).

The dorsal midline remained more displaced in *twi* / *sna* mutants even after prominent local zigzags had largely relaxed (Fig. 7M). RMS deviation quantified this lasting, largely post-closure displacement. Mean RMS deviation was 1.21 ± 0.08 *μ*m in controls, 6 ± 1 *μ*m in *pyr*, 3.7 ± 0.6 *μ*m in *htl*, 19 ± 2 *μ*m in *twi* and 22 ± 4 *μ*m in *sna* (mean ± s.e.m.). Both mesoderm-minus mutants exceeded control (*twi*: adjusted P = 0.00181; *sna*: adjusted P = 0.00629), and their equally weighted genotype mean exceeded *pyr* / *htl* by 15.57 *μ*m (95% confidence interval 10.70–20.45 *μ*m; P = 0.0000343; Table S3). Signed tracks follow individual distortions relative to a fixed reference axis (Fig. 7N; FIG. S13); tracks within an embryo are repeated observations.

Opening-height measurements also showed reduced mean closure rates in *twi* and *sna* relative to controls (5.1 ± 1.2 and 5 ± 1 versus 9 ± 1 nm s^-1^; mean ± s.d.; six embryos per mutant and four controls; Holm-adjusted P = 0.0146 and 0.0127, respectively; FIG. S11B; Table S3). Their mean rates were lower than those of *pyr* and *htl*, but direct comparisons reached significance only against *htl* (*twi*: adjusted P = 0.0439; *sna*: adjusted P = 0.0198); neither comparison with *pyr* was significant (adjusted P = 0.0885 each).

### Individual epithelial distortions develop and retract unevenly

Magnified images of a *sna* embryo resolve growth and retraction (Fig. 8). The cyan-marked distortion grows for 144 min (A^ii^–A^ix^) and retracts over 76 min (A^ix^–A^xv^). The orange-marked distortion develops over 184 min (A^ii^–A^xi^), retracts markedly within 4 min (A^xi^–A^xii^), and is largely retracted within 22 min (A^xiv^). The red-marked configuration changes from A^vi^ through A^xiv^ and largely subsides by A^xv^. Neighboring distortions thus grow gradually but retract over different, shorter intervals. These conspicuous seam distortions are accompanied by stretching of the lateral epidermis and formed dorsal epidermis – indicating that the mesoderm plays an essential role in balancing adhesive and contractile forces that are integral components of a morphogenically active epithelium.

**Figure 8.**
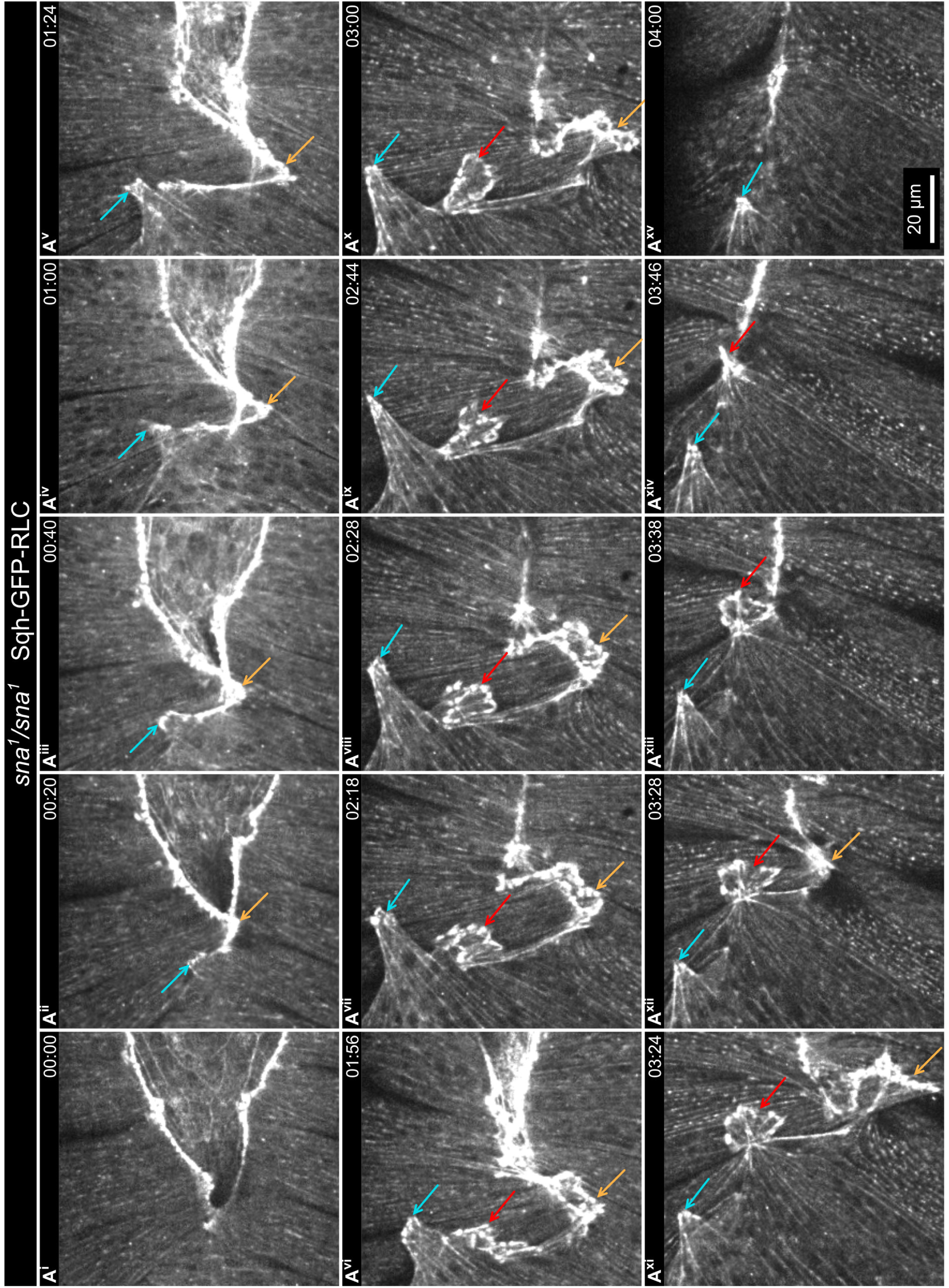
Individual seam distortions grow and retract at different rates in a *sna* embryo. High magnification views of a small region of a *sna* embryo labeled with Sqh-GFP-RLC. Cyan arrows: growth, A^ii^–A^ix^ (144 min); and retraction, A^ix^–A^xv^ (76 min). Orange arrows: growth, A^ii^–A^xi^ (184 min); marked retraction, A^xi^–A^xii^ (4 min), retraction largely complete by A^xiv^ (22 min). Red arrows: local changes in cell arrangements, A^vi^–A^xiv^, largely subsided by A^xv^. Times, h:min from A^i^; acquisition, 2 min/frame. Scale bar, 20 *μ*m.

### Loss of FGF signaling alters the mechanical response of dorsal closure tissues

Loss of Pyr or Htl reduced tissue recoil after laser microsurgery, revealing an altered mechanical response in the closing tissues (Hutson et al., 2003; Peralta et al., 2007; Kiehart et al., 2017). Canthus-to-canthus, A1 cuts bisect the amnioserosa along the dorsal midline and report on forces in the amnioserosa and the arced purse strings that contribute to closure and forces in the bulk of the lateral epidermis that oppose closure (Hutson et al., 2003; Peralta et al., 2007; see Fig. 9A, Movie 20, control; Fig. 9B, Movie 20, *pyr*; Fig. 9C, Movie 20, *htl*). Perpendicular cuts cross the amnioserosa, flanking dorsal most epidermal cells, their included purse strings, and one or two additional rows of lateral epithelial cells (ASPS cuts, Fig. 9D,F,H, Movie 21, control; Fig. 9E,G, Movie 21, *pyr*; Fig. 9I, Movie 21, *htl*). They report on forces along the anterior posterior axis of the embryo. To quantify recoil, bright fluorescent fiducials on either side of the surgical cuts were tracked with time. At 90 s, green pre-cut and magenta post-cut outlines were less separated in mutants than in controls.

**Figure 9.**
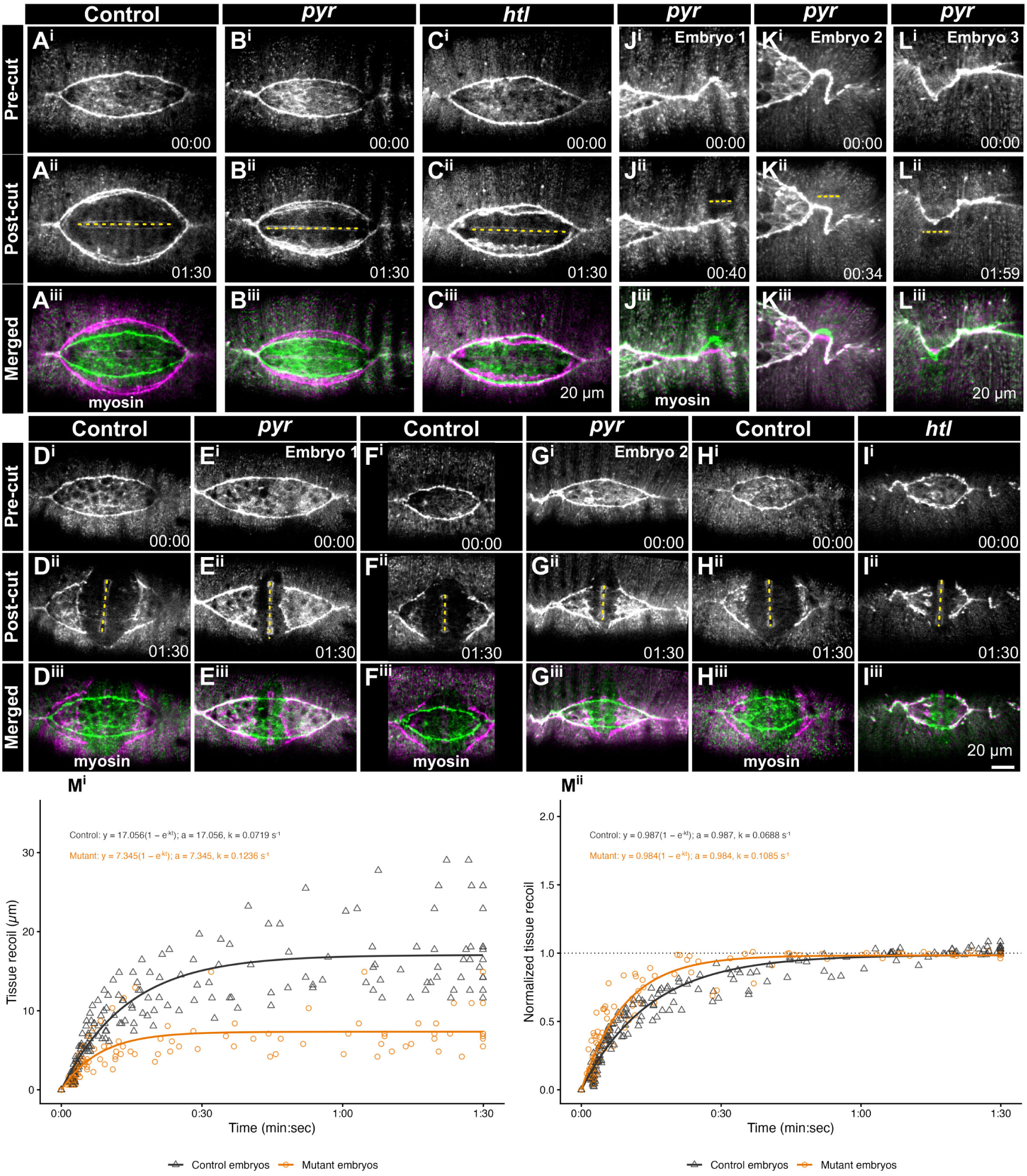
FGF-pathway mutants reduce tissue recoil after laser ablation. A1 amnioserosa cuts. Compare control, *pyr* and *htl* embryos (A–C). ASPS cuts cross the amnioserosa and opposing leading edges. Compare controls and mutants (D–I). Yellow dash-dotted lines mark incision positions in post-cut images. Rows i/ii show pre/post-cut images; row iii overlays pre-cut green and post-cut magenta. Cuts distal to the midline interrogate local zigzag peaks in three *pyr* embryos (J-L). Myosin labels tissue geometry; scale bars, 20 *μ*m; image times, min:s. (M^i^,M^ii^) Raw and individually plateau-normalized A1 recoil, fitted with D(t)=a(1−e^-kt^). Control triangles and *htl* circles track recoil of fiducials away from the surgical cut: seven embryos per genotype, 14 control and 12 mutant fiducials tracked. Inference compares embryo means of raw-track fitted plateau and rate constant using two-sided Welch t-tests. A1 plateau P = 0.003 and k P = 0.018. ASPS cuts were on three controls and six *htl* embryos with four and eight fiducials tracked and are statistically analyzed in Table S3.

After A1 cuts, recoil was smaller in *htl* mutants and approached its plateau faster. Tracks were fitted separately, and plateau and rate-constant k estimates were averaged within each embryo before comparison (14 tracks from seven controls; 12 tracks from seven *htl* embryos). The plateau fell from 17 ± 2 *μ*m to 8 ± 1 *μ*m, a 54% reduction (mean ± s.e.m.; seven embryos per genotype; Welch t-test, P = 0.003), and k increased from 0.068 ± 0.005 to 0.12 ± 0.02 s^-1^ (P = 0.018). Higher k denotes a faster approach to the fitted plateau (Fig. 9M^i^,M^ii^; Table S3).

Recoil was also reduced after ASPS cuts. Within-embryo averaging gave three control values (four tracks) and six *htl* values (eight tracks). The plateau fell from 28.4 ± 0.8 *μ*m to 10.2 ± 0.7 *μ*m, a 64% reduction (P = 1.2 × 10^-5^). The rate constant k was higher in *htl* (0.15 ± 0.04 versus 0.085 ± 0.005 s^-1^), but this difference was not significant (P = 0.16). (FIG. S14B,C; Table S3).

Recoil was reduced after both cut geometries, but the faster approach to the plateau was significant only after A1 cuts. A1 fits appear in Fig. 9M^i^,M^ii^, ASPS fits in FIG. S14B,C and *pyr* maximum-recoil distributions in FIG. S15B–D. Fitted parameters do not isolate tension, elasticity or viscosity.

Short microsurgical lesions near individual zigzag peaks produced variable extents of local recoil or relaxation (Fig. 9J–L; Movie 22), indicating spatial variation in the response to surgically induced, mechanical release (Fig. 9J–L).

### Genetic mapping links a chromosome 3R interval to epithelial instability

An independently identified chromosome 3R deficiency phenotype provided genetic clues to which processes downstream of FGF/FGFR signaling might influence epithelial stability. *Df(3R)BSC141* homozygotes, *Df(3R)BSC141 / Df(3R)BSC517* trans-heterozygotes and embryos homozygous for the overlapping sub-deficiency *Df(3R)BSC488* developed epithelial and seam distortions resembling those of FGF-pathway mutants (a *Df(3R)BSC141* homozygous embryo is shown as an example in Fig. 10A^i^–A^ix^; C^iii^,C^vi^, enlarged views; Movie 23).

**Figure 10.**
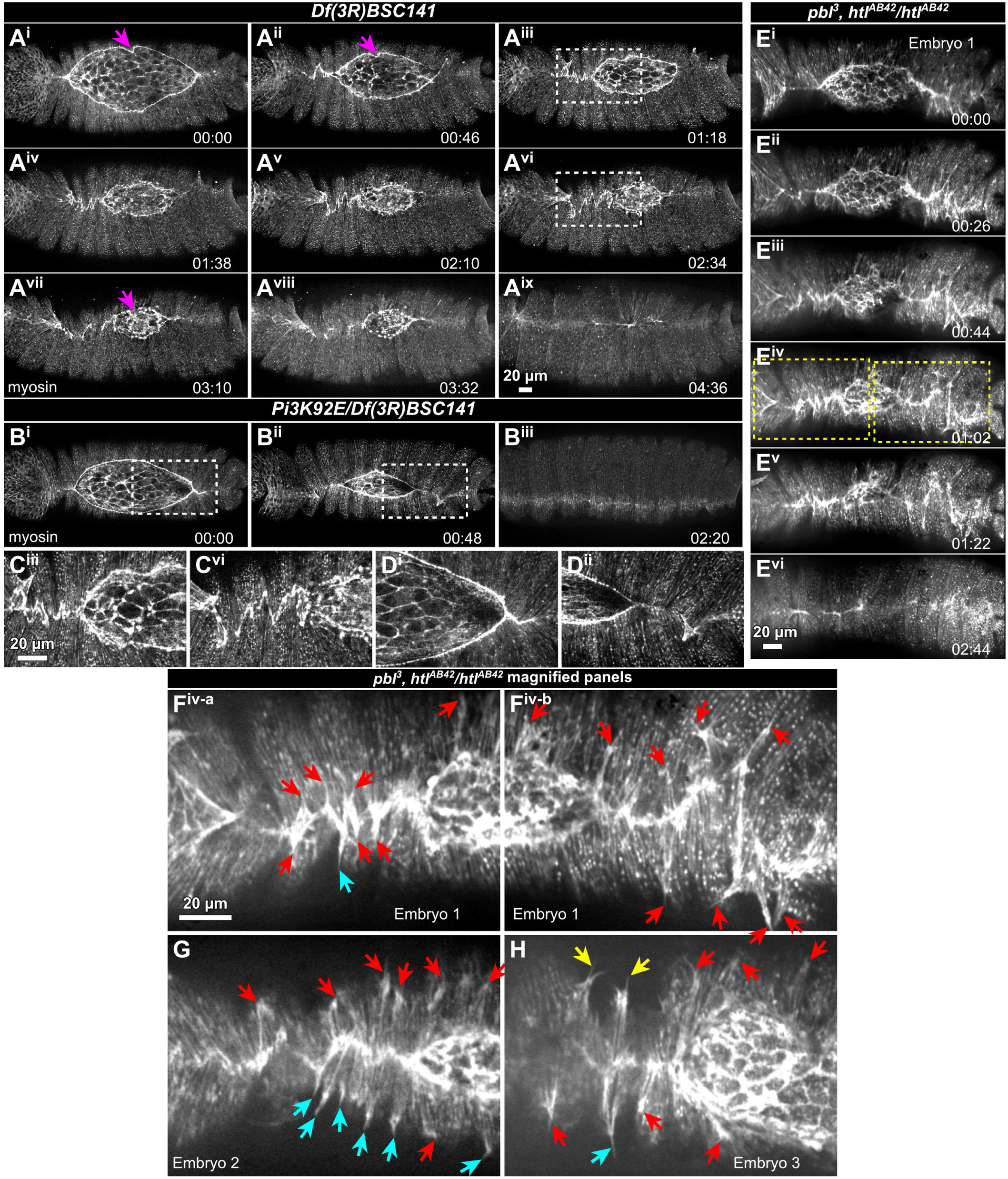
Candidate cortical regulators are associated with epithelial instability. Myosin-labelled *Df(3R)BSC141* homozygotes show changing distortions (A^i^–A^ix^); magenta arrows mark purse-string kinks (A^i^,A^ii^,A^vii^). Conspicuous zigzag distortions in A^iii^ and A^vi^ are shown in C^iii^ and C^vi^. *Pi3K92E / Df(3R)BSC141* images (B^i^–B^iii^; magnified images, D^i^–D^ii^) illustrate more subtle epithelial distortions. An *htl^AB42^* homozygous background carrying *pbl^3^* is shown in E^i^–E^vi^; yellow boxes in E^iv^ are magnified in F^iv^-a and F^iv^-b. G and H show additional embryos. In F^iv^-a/F^iv^-b, G and H, red arrows mark pull-like configurations. Cyan arrows show sharply tapered, extensions that may be filopodia-like. Yellow arrows show bifurcation of an extension into two similarly tapered branches. Times, h:min. Scale bars, 20 *μ*m.

The overlapping deletions encompass the adjacent genes *Pi3K92E* and *Lrrk*, among numerous others (FIG. S16). *Pi3K92E* encodes the class I PI3K catalytic subunit Dp110 and is of interest because receptor tyrosine kinases are frequently involved in cortical and cytoskeletal regulation (Leevers et al., 1996; Weinkove et al., 1997; Sun et al., 2020). *Lrrk* encodes a leucine-rich repeat serine/threonine kinase with reported roles in actin regulation (Bardai et al., 2018). Possible synergy between *Pi3K92E* and *Lrrk* is considered in the Discussion. Most *Pi3K92E / Df(3R)BSC141* embryos lacked conspicuous distortions, although one embryo showed subtle but observable epithelial distortions (Fig. 10B,D^i^–D^ii^; Movie 24). Available individual *Pi3K92E* or *Lrrk* homozygotes did not robustly reproduce the deficiency phenotype (data not shown).

A second tantalizing observation linked Htl-dependent epithelial stability to cortical regulation. In an *htl* mutant background, reducing *pebble* (*pbl*) dosage occasionally accentuated F-actin-reporter-positive pull-like and protrusive configurations (Fig. 10E–H; Movie 25). In F^iv^-a/F^iv^-b, G and H, red arrows mark prominent pull-like configurations; cyan arrows highlight sharply tapered, filopodia-like extensions, and yellow arrows mark bifurcation into two similarly tapered branches. These observations suggest a connection between Htl signaling and cortical regulation without establishing a common Htl–PI3K–Pbl pathway or a tissue-specific interaction.

## Discussion

Mesodermal Htl/FGFR signaling contributes to ectodermal epithelial stability during dorsal closure (Fig. 11B^i^). Loss of Pyr, Ths, Htl or Stumps produces pronounced, kinematically active distortions, yet closure continues. Mesodermal Htl expression restores stability, whereas ectodermal expression does not. Genetic ablation of mesoderm produces more extensive instability and post-closure midline deviation. Preliminary *Pi3K92E*, *Lrrk* and *pbl* observations suggest a key role for the cortical arm of FGF signaling in stabilizing the ectoderm. We favor a working hypothesis in which mesodermal distribution and cortical activity influence the overlying ectoderm. We cannot rule out additional input from the MAPK arm of the pathway.

**Figure 11.**
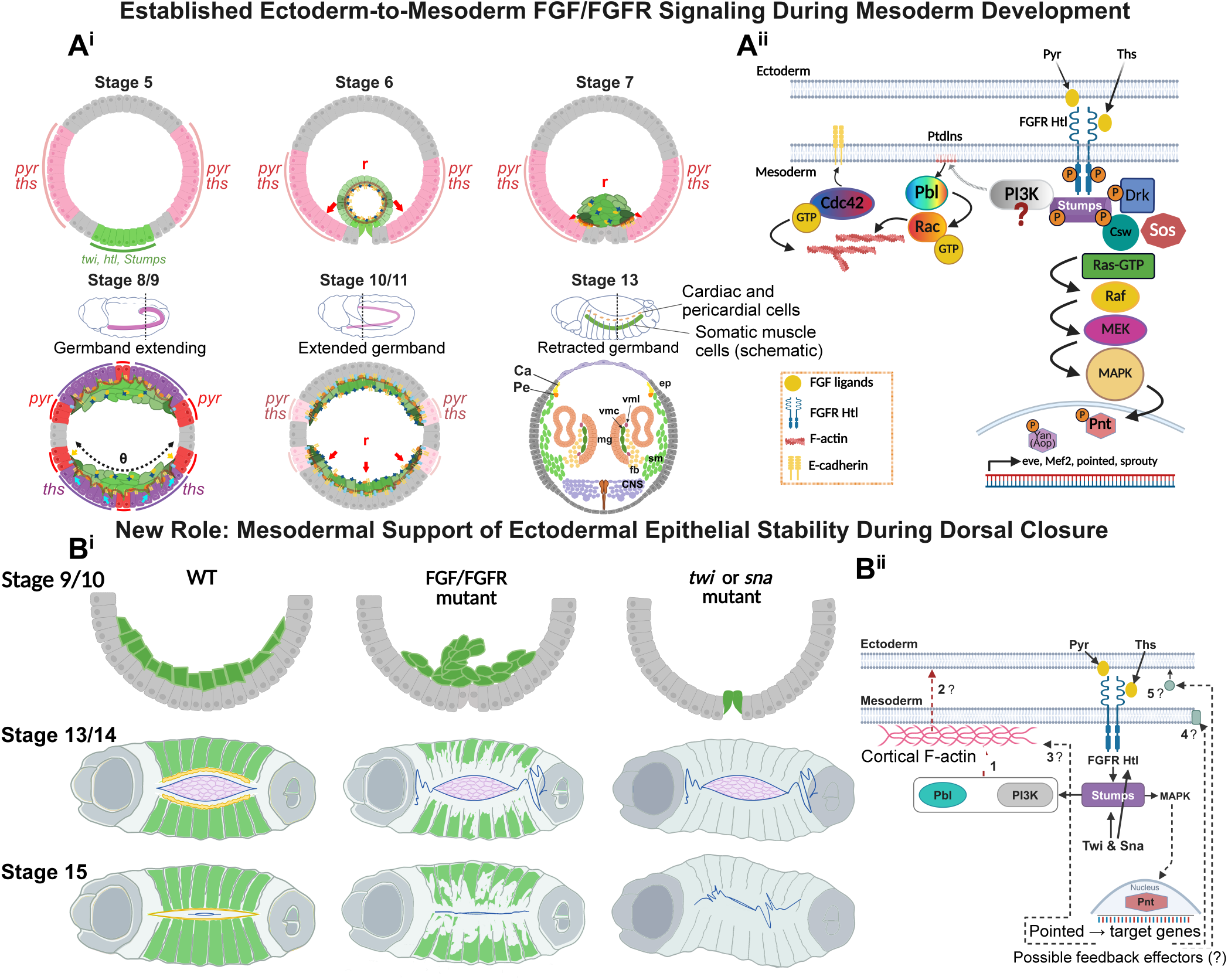
FGF signaling within the mesoderm supports ectodermal stability. (A^i^) Schematics of mesoderm development and germband movement with stages indicated. θ denotes angular dorsal spreading along the ectoderm; r denotes radial movement during tube collapse and intercalation into a monolayer (McMahon et al., 2008, 2010). Ca, cardiac cells; Pe, pericardial cells; ep, epithelium; sm, somatic muscles; fb, fat body; mg, midgut; vmc/vml, circular/longitudinal visceral muscle; CNS, central nervous system. (A^ii^) Established Htl signaling, including cortical regulators and Ras–MAPK outputs to Pointed (Pnt) and phosphorylated Yan (Aop). Redrawn from Muha and Müller, 2013 and Sun et al., 2020. (B^i^) Comparative schematic of WT, FGF/FGFR mutants and *twi* / *sna* mutants: green, mesodermal/muscle groups; violet, amnioserosa; blue, purse strings/seams; and yellow, cardiac structures. Cell numbers, proportions and outlines are schematic. (B^ii^) Possible routes by which FGF signaling may impact ectodermal epidermis stability and dynamics are indicated by dashed arrows and or numbers with question marks. (1) PI3K/Pbl-dependent cortical regulation; (2) cortical influence on ectodermal stability through mechanical interactions; MAPK-dependent effectors acting through (3) the mesodermal cortex, (4) mesodermal membrane proteins or (5) secreted signals. Cortical F-actin is shown over only part of the inner membrane for schematic clarity. The Src input into MAPK is omitted.

### Epithelial stability is separable from the progression of dorsal closure

Dorsal closure can reach completion even when the closing epithelium fails to maintain stable geometry. Forces generated by the amnioserosa, leading-edge actomyosin cables, zippering and the lateral epithelium allow compensation following surgical or genetic perturbation (Kiehart et al., 2000; Harden, 2002; Hutson et al., 2003; Peralta et al., 2007; Wells et al., 2014; Hayes and Solon, 2017; Kiehart et al., 2017). Similarly, *wingless* mutants can complete closure despite abnormal epithelial cell shapes and leading-edge organization (McEwen et al., 2000; Kaltschmidt et al., 2002). The extensive, continuously changing distortions in FGF/ FGFR mutants reveal further robustness of closure: the movement succeeds despite epithelial instability caused by defective mesodermal function.

The unstable epithelium undergoes pronounced pulls and rapid relaxation, consistent with altered coupling between contractility and adhesion. Rapid recoil of distorted cell boundaries could reflect release of local attachments, but the molecular targets of FGF signaling and the identities and behavior of those attachments remain unresolved.

The zigzag seam, readily visualized with fluorescent F-actin and myosin reporters, is a conspicuous readout of epithelial instability affecting both the lateral and newly formed dorsal epithelium when mesodermal function is perturbed. Distorted seams suggest compromised coordination of cell shape, cell–cell and cell–matrix adhesion, actomyosin contractility and tissue mechanics. Resolving junctional remodeling, cortical contractility and mechanical loading requires distinguishing cortical and MAPK-dependent contributions of Htl.

### Mesoderm organization and signaling to the ectoderm

The contribution of mesodermal Htl to ectodermal stability reveals a reciprocal consequence of the established ectoderm-to-mesoderm FGF system (Muha and Müller, 2013; Sun et al., 2020; Fig. 11A^i^). Ectodermal Pyr and Ths activate Htl in the invaginated mesoderm, promoting spreading and mesoderm–ectoderm contact. This positions mesodermal cells to receive inductive signals that specify differentiated lineages. Our findings reveal a later consequence for the signaling tissue: mesodermal Htl function contributes to overlying ectodermal stability during closure.

Localization and reciprocal expression identify a mesodermal route to ectodermal stabilization. Htl localizes to mesodermal derivatives during closure (Fig. 4), and mesodermal expression restores epithelial stability in an *htl* mutant background, whereas ectodermal expression does not eliminate distortions (Fig. 5). Whether Htl acts during closure or earlier to establish mesoderm that subsequently contributes to stability remains an open question.

Loss of FGF/FGFR signaling disrupts mesodermal organization without eliminating the tissue. *htl* mesoderm continues to spread and move dorsally despite disrupted somatic-muscle architecture and loss of ordered cardiogenic rows (Fig. 6). In the imaged embryos, mesodermal disorganization and overlying epithelial distortions did not show a simple spatial correspondence, arguing against assigning each zigzag to an immediately underlying mesodermal gap. Htl-dependent development may instead shape a mechanical or signaling environment experienced across the closing epithelium.

Genetic ablation distinguishes loss of mesoderm from disruption of its Htl-dependent organization. *twi* and *sna* mutants fail to establish internal mesoderm, and their epithelial instability persists longer and extends more conspicuously into the lateral epithelium. Residual mesoderm in FGF-pathway mutants may retain stabilizing contributions that become evident when closure finishes. Why epithelial distortions largely relax after FGF-pathway disruption but endure after mesoderm loss remains unresolved. The reduced mean closure rates in *twi* and *sna* need not result solely from loss of internal mesoderm: both genes regulate broad developmental programs, so altered tissue specification and earlier morphogenesis could also contribute. These experiments do not separate these possibilities.

Differences among ligand-mutant phenotypes may reflect how Pyr and Ths organize mesoderm for its later contribution to epithelial stability. The ligands activate the same receptor but differ in their molecular properties and distributions: Pyr is membrane tethered through a transmembrane domain, whereas Ths is diffusible (Stepanik et al., 2020; Sun et al., 2020). Both stabilize epithelia; penetrance and expressivity vary particularly in lateral epithelial abnormalities. Distinct spatial contributions to mesodermal organization could explain this variation without ligand-specific epithelial functions. Comparing mesodermal organization between ligand transheterozygotes could test this possibility.

### FGF/FGFR signaling influences tissue mechanics in dorsal closure

Microsurgical cuts (A1 and ASPS) link epithelial instability to reduced recoil after FGF/FGFR-pathway disruption. In *htl* mutants, recoil approaches its fitted plateau faster after A1 cuts, whereas the ASPS rate difference is not significant. These responses implicate altered amnioserosa and epithelial mechanics. Recoil integrates pre-existing load with passive material properties and active responses after severing, so reduced displacement cannot be equated with reduced tension alone.

Reduced recoil can accompany modest visible distortion; local geometry and mechanical response therefore need not change together. Htl signaling may influence the environment in which closure’s force-producing and adhesive systems operate, with zigzags emerging when force generation, force transmission and local attachments become unbalanced and/or poorly coordinated. The larger recoil plateau and slower approach in controls are consistent with an Htl-dependent mesodermal contribution to load transmission or mechanical relaxation in the coupled tissues. These fitted responses do not distinguish altered prestress from changes in effective stiffness or dissipation.

### Candidate signaling mechanisms downstream of Htl

How might mesodermal Htl activity stabilize the neighboring ectoderm? Candidate inputs include MAPK-dependent gene expression and cortical regulation involving Rho-family GTPases and potentially Pi3K92E (Muha and Müller, 2013; Fig. 11B^ii^). Htl-dependent protrusive activity and mesoderm–ectoderm contact provide a developmental precedent for coupling cytoskeletal behavior to tissue interactions (Schumacher et al., 2004; Sun and Stathopoulos, 2018).

A series of tantalizing observations motivates investigation of cortical regulation: low-penetrance distortions in the tested *Pi3K92E* combination, zigzag seams in overlapping chromosome 3R deficiencies that remove *Pi3K92E* and *Lrrk*, and occasional extensive protrusive expressivity in *pbl/+; htl / htl* embryos (Fig. 10). These deletions remove additional genes, so a targeted deletion of the adjacent *Pi3K92E* and *Lrrk* genes would test their combined contribution more directly; their close linkage makes targeted engineering preferable to conventional recombination. Lrrk can regulate F-actin organization in *Drosophila* neurons (Bardai et al., 2018), providing a rationale for examining it alongside Pi3K92E without establishing functional synergy in dorsal closure. Pebble regulates protrusive activity and mesodermal migration downstream of Htl (Schumacher et al., 2004; Smallhorn et al., 2004). These observations suggest how cytoskeletal regulation could alter migration or tissue interactions. The relevant Pi3K92E, Lrrk or Pbl activity could reside in mesoderm, ectoderm or both.

PI3K could couple cortical regulation to adhesion by generating phosphatidylinositol (3,4,5)-trisphosphate (PIP₃) at the membrane. In *Drosophila*, the PIP₃-binding region of Myoblast city is required for myoblast fusion (Balagopalan et al., 2006); in mammalian cells, PIP₃-rich protrusions support integrin recycling through a CYTH2–ARF6–AGAP1 module (Nikolatou et al., 2023). These findings provide precedents for lipid signalling to coordinate actin organization and membrane interactions. Such mechanisms could converge with Lrrk-dependent cortical regulation in mesoderm, ectoderm or both.

Htl–MAPK-dependent gene expression offers a complementary route to stabilization. Its products could modify the mesodermal cortex, alter membrane interactions or provide secreted signals to the ectoderm (Fig. 11B^ii^). Comparing mesodermal and ectodermal transcriptomes between mutant and wild-type embryos by single-cell RNA sequencing could identify differentially expressed genes encoding cortical regulators, membrane proteins or signals to the ectoderm, as well as ectodermal responses. Cortical regulation and transcriptional outputs may therefore act together to influence the mechanical and adhesive environment of the closing epithelium.

We contend that this remarkable new dorsal closure phenotype is worth pursuing aggressively, to identify its molecular basis or bases. This provides a route to identifying new ways in which mesoderm can affect ectodermal function during development.

## Materials and Methods

### *Drosophila* stocks and genetics

*Drosophila melanogaster* stocks and crosses were maintained using standard protocols (Hales et al., 2015). Stocks, alleles, fluorescent reporters, balancers and experimental genotype combinations are listed in Table S1. Deficiency screens and embryo-selection strategies paralleled Mortensen et al. (2018) and Fogerson et al. (2020). Tissue-specific expression used the GAL4/UAS system (Brand and Perrimon, 1993).

Embryos homozygous for *Df(2R)ED2247* (*Df(2R)22*) were previously described as having abnormal amnioserosa ingression and loss of amnioserosa integrity (Mortensen et al., 2018). Overlapping deficiencies included *Df(2R)BSC259* (22B), *Df(2R)BSC153* (22C) and *Df(2R)BSC329* (22D). Mapping records distinguish epithelial instability without amnioserosa disintegration in 22B and 22F from the combined phenotype in 22C (FIG. S3C). *Df(2R)22F* was generated in the Kiehart laboratory by CRISPR-mediated homology-directed repair. We will donate the new Df flies to the Bloomington Stock Center upon publication. Independent *pyr* and *ths* alleles and deficiencies were examined to test each ligand’s contribution.

As is custom, we used transheterozygous combinations of chromosomes (*e.g.*, allele/Df) whenever possible. Homozygous mutant embryos were used when their phenotype matched that of an appropriate transheterozygous combination. In addition we use italicized font to designate genes and normal font and capitalized first letters for proteins (*e.g.*, *htl* and Htl).

Chromosome 3R deficiency mapping examined *Df(3R)BSC141* homozygotes, *Df(3R)BSC141 / Df(3R)BSC517* trans-heterozygotes and the overlapping sub-deficiency *Df(3R)BSC488* (BDSC #9501, #25021 and #24992, respectively; FIG. S16). Available *Pi3K92E* and *Lrrk* alleles were subsequently examined as listed in Table S1. Genomic annotations were obtained from FlyBase (Öztürk-Çolak et al., 2024).

### Live imaging of control and mutant embryos

Fluorescent reporters marked F-actin (sGMCA), non-muscle myosin II (Sqh-GFP-RLC; Karess et al., 1991; Royou et al., 2002, 2004) and adherens junctions (E-cadherin-GFP). sGMCA contains the GFP-tagged Moesin actin-binding domain under the spaghetti squash promoter (Kiehart et al., 2000); the GAL4/UAS-driven GFP–Moesin actin-binding domain reporter is described by Dutta et al. (2002). For simultaneous imaging of mesoderm and ectoderm, *twist-GAL4^PB^*drove *UAS-RFP-Moesin actin-binding domain (Millard and Martin, 2008)* in the mesoderm and E-cadherin fluorescence marked epithelial junctions. Htl-mCherry was described previously (Irizarry and Stathopoulos, 2015; Sun and Stathopoulos, 2018). Reporter identities and sources are listed in Table S1.

Fluorescently marked balancers were selected against to identify mutant embryos. Controls expressed the same reporter as the corresponding mutant or deficiency background; comparisons were interpreted within matched reporter and genetic backgrounds.

### Embryo collection and preparation

Embryos were collected and selected for live imaging as described previously (Mortensen et al., 2018; Fogerson et al., 2020). Timed collections comprised 2–4 h egg lays at 25°C followed by approximately 24 h of aging at 16°C. Alternatively, embryos at dorsal-closure stages were selected directly from overnight collections. Embryos lacking balancer-associated fluorescence were selected using a Zeiss Discovery V12 SteREO fluorescence dissecting microscope. Selected embryos and appropriately staged controls collected in parallel were aligned, attached to coverslips with embryo glue (Sullivan et al., 2000), covered with halocarbon oil and mounted in an imaging chamber (Kiehart et al., 2006). Reagent sources are listed in Table S2.

### Microscopy and image processing

Four spinning-disk confocal imaging systems were used. Earlier datasets were acquired with a Yokogawa CSU-10 spinning-disk head on either a Zeiss Axiovert 200M with a Hamamatsu ORCA-ER CCD camera and MetaMorph software (Molecular Devices; 2×2 binning), or a Zeiss Axio Imager.M2m with a Hamamatsu C9100 EM-CCD camera and Micro-Manager 1.4 or 2.0 (1×1 binning). Objectives included Zeiss 40× 0.9 NA multi-immersion, 40× 1.3 NA oil-immersion and 40× 1.2 NA water-immersion lenses, as appropriate to the experiment.

Newer systems were configured around Yokogawa CSU-W1 confocal heads. The upright system used a Hamamatsu Orca Quest 2 qCMOS camera. The inverted system used a Hamamatsu Orca Fusion BT CMOS camera, 1×1 or 2×2 binning and a 40× 1.2 NA glycerol-immersion objective. Time-lapse z-stacks generally contained 11–16 planes; the illustrated wild-type two-color recording contained 41 Z planes and 94 time points. The wild-type mesoderm–ectoderm and Htl-mCherry/E-cadherin two-color recordings had median acquisition intervals of approximately 3.1 and 4.3 min, respectively; elapsed times were determined from image timestamps.

Images were processed in Fiji/ImageJ2 (Schindelin et al., 2012). Selected Z-stacks were converted to maximum-intensity projections. Orientation, cropping and brightness/contrast adjustments were used for presentation; comparative adjustments were applied consistently within experiments. Other measurements were made in Fiji before figure assembly. For the descriptive distances in Fig. 6, native merged images were measured manually at 3.15 pixels *μ*m^-1^ along four selected central image-vertical transects per frame (two per side) in one embryo. The reference was the opening edge before closure and the formed seam afterward; resolved cardiac protrusive margins and the trailing cable were measured separately.

### Scoring epithelial instability

Dorsal epithelial distortions were classified as absent (indistinguishable from controls), subtle but observable, or prominent. Penetrance is the proportion of analyzed embryos displaying distortion; expressivity is its qualitative severity. Scores and denominators were evaluated within the corresponding reporter and genetic background.

### Dorsal seam tortuosity

Seam tortuosity was calculated for each measured frame as:

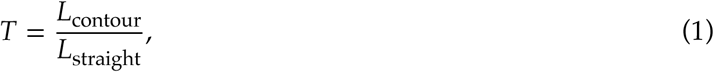

the traced seam length divided by the straight-line distance between the same endpoints. T = 1 describes a straight seam; larger values indicate greater distortion. Mutant analysis began at detectable tortuous behavior; control seams were followed from an arbitrarily selected dorsal-opening height over successive frames. Measurements were repeated within embryos. Each embryo was the biological replicate. Technically analyzable, strongly expressive embryos were selected across all four mutant genotypes; penetrance was evaluated separately.

Peak seam tortuosity was compared using a one-way model with genotype as a categorical factor. Dorsal-opening height was measured in micrometers at the same frames as tortuosity to describe closure progression. The dataset comprises 284 paired measurements from 28 embryos (control, n=4; each mutant, n=6). Images were acquired every 2 min. Height was treated as a geometric measure of closure progression, not developmental age.

Seam measurements after closure were timed based on each embryo’s first closed frame (*i.e.*, no dorsal opening). Tortuosity for a given seam segment was tracked through time. Tortuosity in frames 10 and 20 min after closure was determined, yielding two measurements per embryo across 28 embryos. Mean tortuosity across the two time points supplied one summary per embryo for Fig. 7L; mutant-class means were compared by a two-sided Welch test. A repeated-measures model with genotype, post-closure time and their interaction and an embryo random intercept estimated equal-genotype-weight contrasts of *twi* / *sna* versus *pyr* / *htl* at each time. The two time-specific tests were Holm adjusted together. The sampled mutant embryos were selected for clear distortions and a traceable seam, so these comparisons do not estimate population penetrance.

### Dorsal midline deviation

Deviations of seams from the dorsal midline were measured in one late image per embryo after prominent local zigzags had largely subsided. A straight reference axis followed the overall anterior posterior axis specified by the position of the canthi. Perpendicular distances from the observed seam to this axis were measured (in *μ*m) at 8–14 distributed positions along the analyzable seam, approximately evenly spaced where possible. Frame selection was based on the residual-deviation endpoint rather than a fixed dorsal-opening-height threshold.

One RMS value was calculated per embryo as:

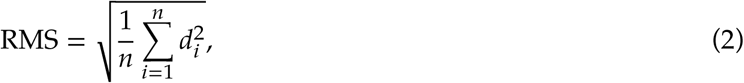

where d_i_ is a perpendicular displacement and n is the number of positions in that embryo. Raw values in *μ*m were plotted with mean ± s.e.m. Mutants were compared with control on the raw RMS scale using two-sided Welch tests, with Holm adjustment across the four mutant–control comparisons. A separate contrast compared mean(*twi*, *sna*) with mean(*pyr*, *htl*), weighting each genotype equally and using a Welch–Satterthwaite standard error and degrees of freedom. The embryo was the statistical unit. RMS measures displacement amplitude; it is distinct from the dimensionless contour-length ratio T.

### Individual zigzag seam displacement trajectories

Spatially distinct zigzag seam distortions were tracked through time on one side of an embryo (anterior or posterior). For each track, the maximum perpendicular displacement from a straight dorsal reference axis was retained with a sign denoting its side of that axis. Reference axes were held fixed through a sequence, with whole-embryo movement corrected when necessary to preserve the biological reference. Times recorded as h:min were converted to elapsed minutes; acquisition-frame indices in the multi-genotype dataset were converted using 2 min/frame. Individual tracks were plotted rather than averaged. Multiple tracks from one embryo are repeated local observations, not independent biological replicates. The *pyr*-only and multi-genotype datasets were analyzed separately.

### Dorsal closure rate

Dorsal-opening height was measured between the opposing leading edges at a consistent position near the center of the opening. For each embryo, an ordinary least-squares regression estimated the slope of opening height against elapsed time, through the first measured closed frame when present; subsequent zero-height measurements were excluded. Narrowing rate was the negative slope, expressed in nm/s. Rates were compared at the embryo level using two-sided Welch tests, with Holm adjustment across the four mutant–control comparisons.

Images in the closure-rate dataset were acquired every 2 min. The four direct comparisons of *twi* or *sna* with *pyr* or *htl* formed a separate Holm-adjusted family. Htl images used an F-actin reporter, whereas the other cohorts used Sqh; comparisons involving *htl* therefore also differ in marker background. A separate cohort of seven *htl* embryos expressing Sqh-GFP-RLC was analysed using the same per-embryo linear-regression approach (FIG. S8). This cohort comprised four homozygous and three heteroallelic *htl* embryos. Embryo-level rates were compared with Sqh-GFP-RLC-labelled controls using a two-sided Welch t-test.

### UV laser ablation

Ultraviolet (355 nm) microsurgery compared control, *pyr* and *htl* tissues. The beam from a Minilite II Nd:YAG laser (Continuum, San Jose, CA) entered the epi-port of a Zeiss Axio Imager.M2m/Yokogawa spinning-disk microscope and was focused through a Zeiss 40× 1.2 NA water-immersion objective onto the specimen. A beam-steering system drove defined A1 and ASPS (see Results) incisions at the specimen plane (Kiehart et al., 2006, section VIII and fig. 8). Images were acquired before, during and after ablation. A mark on the vitelline envelope was used to verify incision position and compare cut placement and length. Pre- and post-surgery overlays illustrated tissue displacement. Local cuts at zigzag peaks were analyzed qualitatively.

### Recoil measurement and modeling

One or two retained fiducial marks per cut per embryo were followed relative to the pre-surgical configuration. Embryo and fiducial counts are reported separately. Recoil was modeled as:

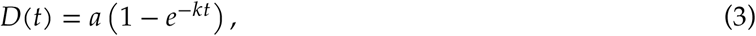

with displacement D in *μ*m, time t in s, plateau a in *μ*m and rate constant k in s^-1^. Each fiducial track was fitted separately by exponential least squares in R, with a and k constrained to be non-negative. Parameter estimates from markers in the same embryo were averaged, and independent embryo averages were compared using two-sided Welch t-tests, without multiplicity adjustment across the four cut-by-parameter comparisons. A1 cuts included seven embryos per genotype (14 control and 12 mutant tracks); ASPS included three control and six mutant embryos (four and eight tracks). For display, raw and individually plateau-normalized observations were pooled within genotype to plot descriptive curves. For both A1 and ASPS, each track was normalized by its own fitted plateau; normalized fits were not used for additional hypothesis tests. Fitted parameters describe recoil phenomenologically and do not isolate tension, pre-stress, elasticity or viscosity. Maximum-recoil distributions were compared with the corresponding reporter control using two-sided Welch t-tests; the violin plots display unadjusted P values.

### HCR-FISH

Whole-mount hybridization chain reaction fluorescence in situ hybridization (HCR-FISH) used Molecular Instruments HCR Gold reagents and HiFi probes for *htl*, *odd* and *Mef2*, with amplifiers X2/647 nm, X1/ 546 nm and X3/633 nm, respectively. *odd* and *Mef2* provided ectodermal and mesodermal spatial references. The sGMCA reporter was imaged with 488-nm excitation for morphological context and was not an HCR-FISH target. Five conditions and their exact genotype combinations are listed in Table S1; GAL4-only and UAS-only controls were included for HCR-FISH but not the live-imaging rescue experiment. Embryos were fixed, methanol-devitellinized and processed through hybridization, probe washes and hairpin amplification before mounting in VECTASHIELD.

Images were acquired on a Leica STELLARIS 8 with LAS X. Acquisition settings were held constant across the five backgrounds within the experiment while avoiding saturation and minimizing spectral overlap. Reagent identities, amplifier assignments and catalog details are listed in Table S2.

### Statistical reporting

Analyses used R 4.5.2 with lme4 2.0-6, lmerTest 3.2-1, emmeans 2.0.4 and ggplot2 4.0.3. Unless specified otherwise, tests were two-sided. Estimates and their associated uncertainties are reported to matching decimal places. Exact or bounded P values, adjustment methods and sample sizes are provided in Table S3. Repeated observations and multiple tracks within one embryo were distinguished from independent embryos. Figure legends identify biological replicates, repeated tracks and error bars. Model estimates are reported with standard errors (s.e.). Standard deviation (s.d.) describes between-embryo variation; s.e.m. denotes the standard error of the mean. F(df1, df2) gives the numerator and denominator degrees of freedom, respectively; d.f. denotes degrees of freedom for other tests.

## Use of AI

AI-assisted tools supported manuscript organization, language editing, proofreading, and precision of figure layout in Inkscape.

## Acknowledgements

We acknowledge important conceptual inputs or the gift of fly stocks from Prof. Amy Bejsovec, Danny Lew, Kaelyn Sumigray, Dave McClay, Angelike Stathopoulos, Zahra Zarei, Hans-Arno Müller, Eric Spana and Kevin Cook at the Bloomington Stock Center. Stocks obtained from the Bloomington *Drosophila* Stock Center (NIH P40OD018537) were used in this study. Bioinformatics available on FlyBase were invaluable. This work was supported by NIH grant R35GM127059 to DPK.

---

**Figure S1.**
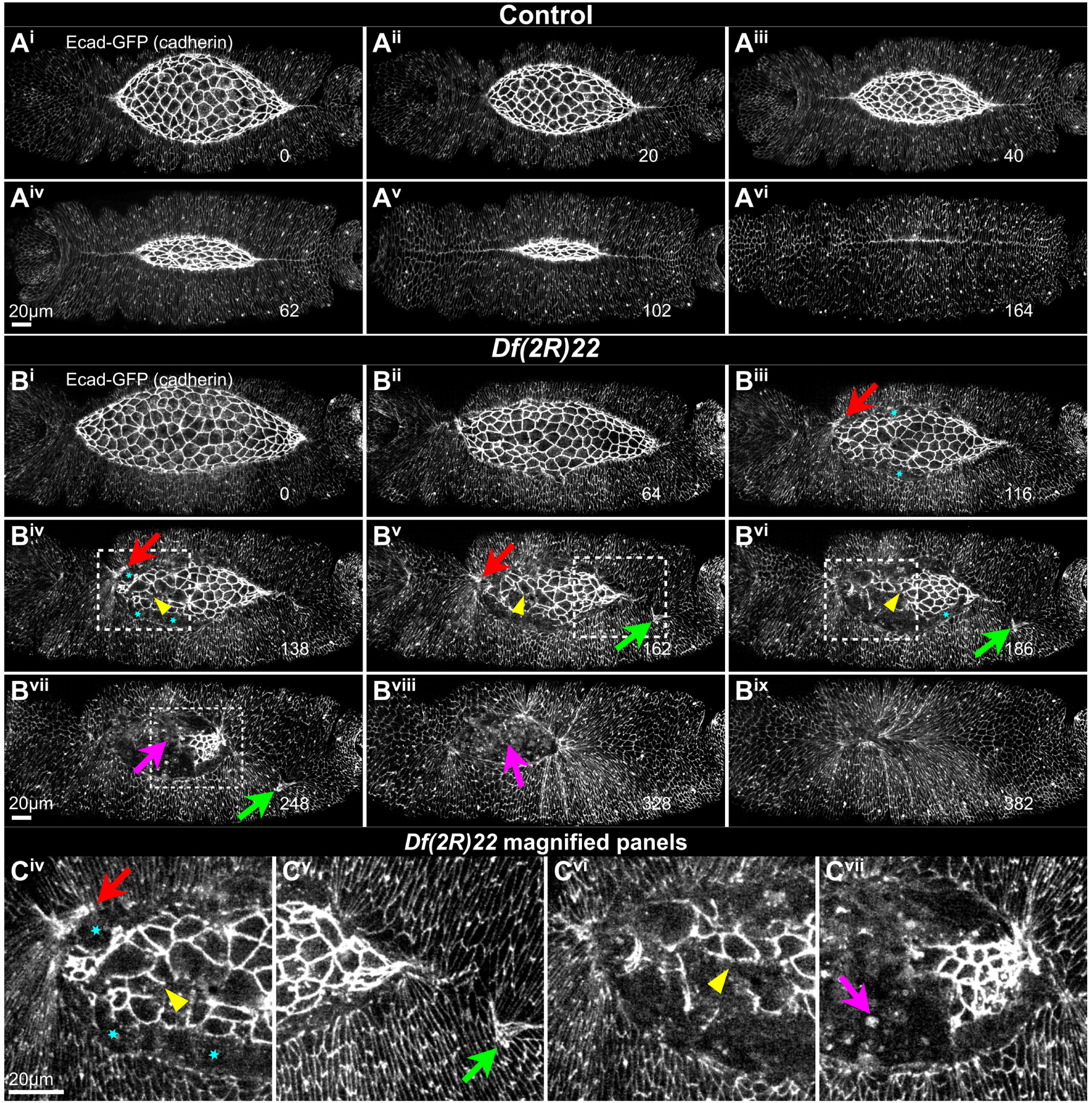
Cadherin imaging of amnioserosa integrity during dorsal closure in control and *Df(2R)22* homozygous embryos. E-cad-GFP labels junctions in control (A^i^–A^vi^) and *Df(2R)22* (B^i^–B^ix^) embryos. The mutant series shows progressive disruption of amnioserosa junctional organization and its interface with the epithelium. Dashed boxes in B^iv^–B^vii^ identify the regions enlarged in C^iv^–C^vii^. Cyan asterisks indicate PAS–DME separation. Red and green arrows indicate regions of epithelial distortion. Yellow arrowheads indicate local junctional discontinuities. Magenta arrows indicate cellular debris. Times are elapsed minutes. Scale bars, 20 *μ*m.

**Figure S2.**
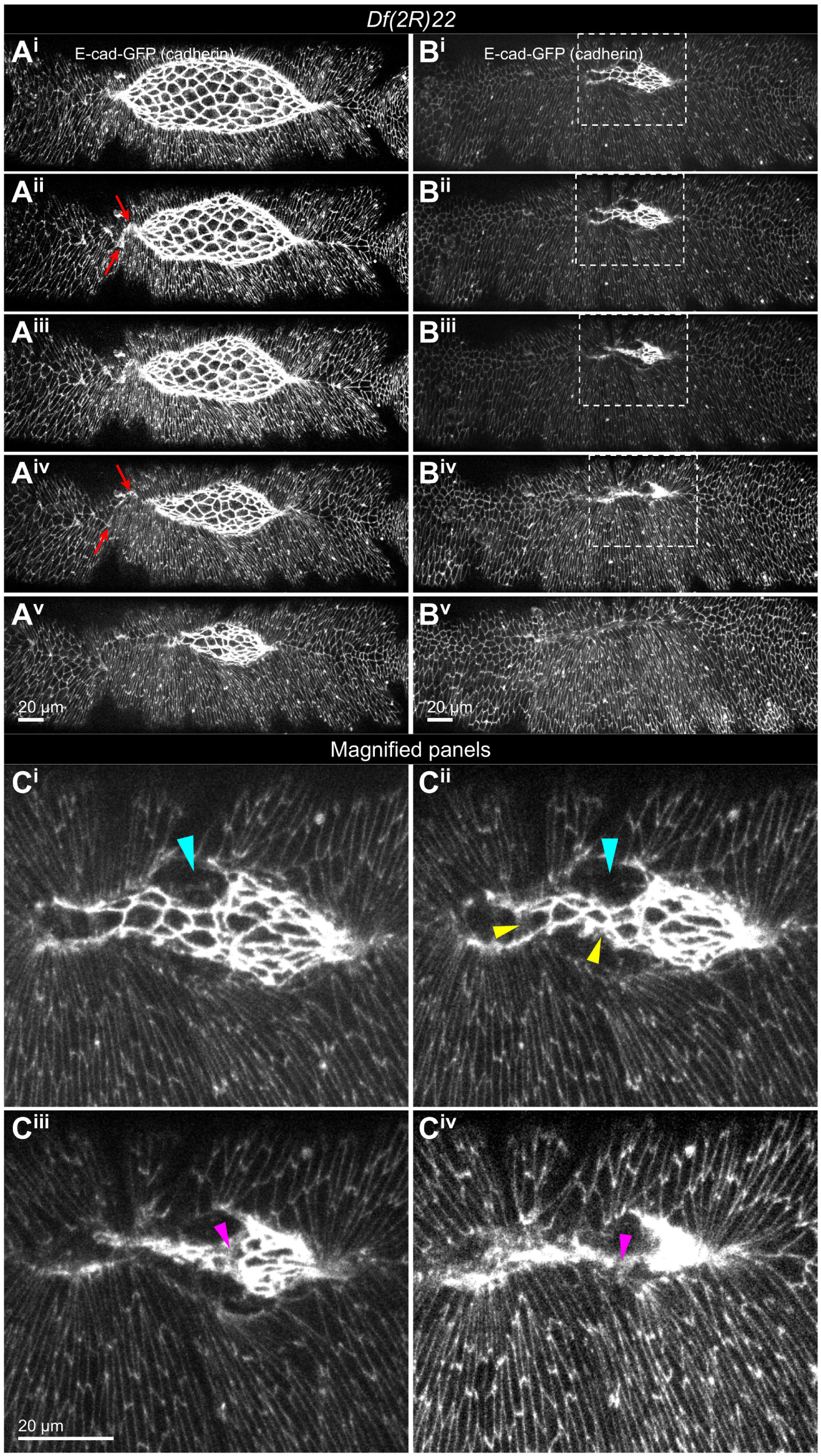
Cadherin labeled *Df(2R)22* embryos with amnioserosa defects in late closure. A *Df(2R)22* embryo is shown in successive frames (A^i^–B^v^). E-cad-GFP labels cadherin-containing junctions. Red arrows mark epithelial distortions (A^ii^,A^iv^). Dashed boxes in B^i^–B^iv^ identify the regions enlarged in C^i^–C^iv^, respectively. Cyan arrowheads indicate PAS–DME separations (C^i^,C^ii^); yellow arrowheads indicate local junctional discontinuities (C^ii^); magenta arrowheads indicate disintegration of amnioserosa cells (C^iii^,C^iv^). Scale bars, 20 *μ*m.

**Figure S3.**
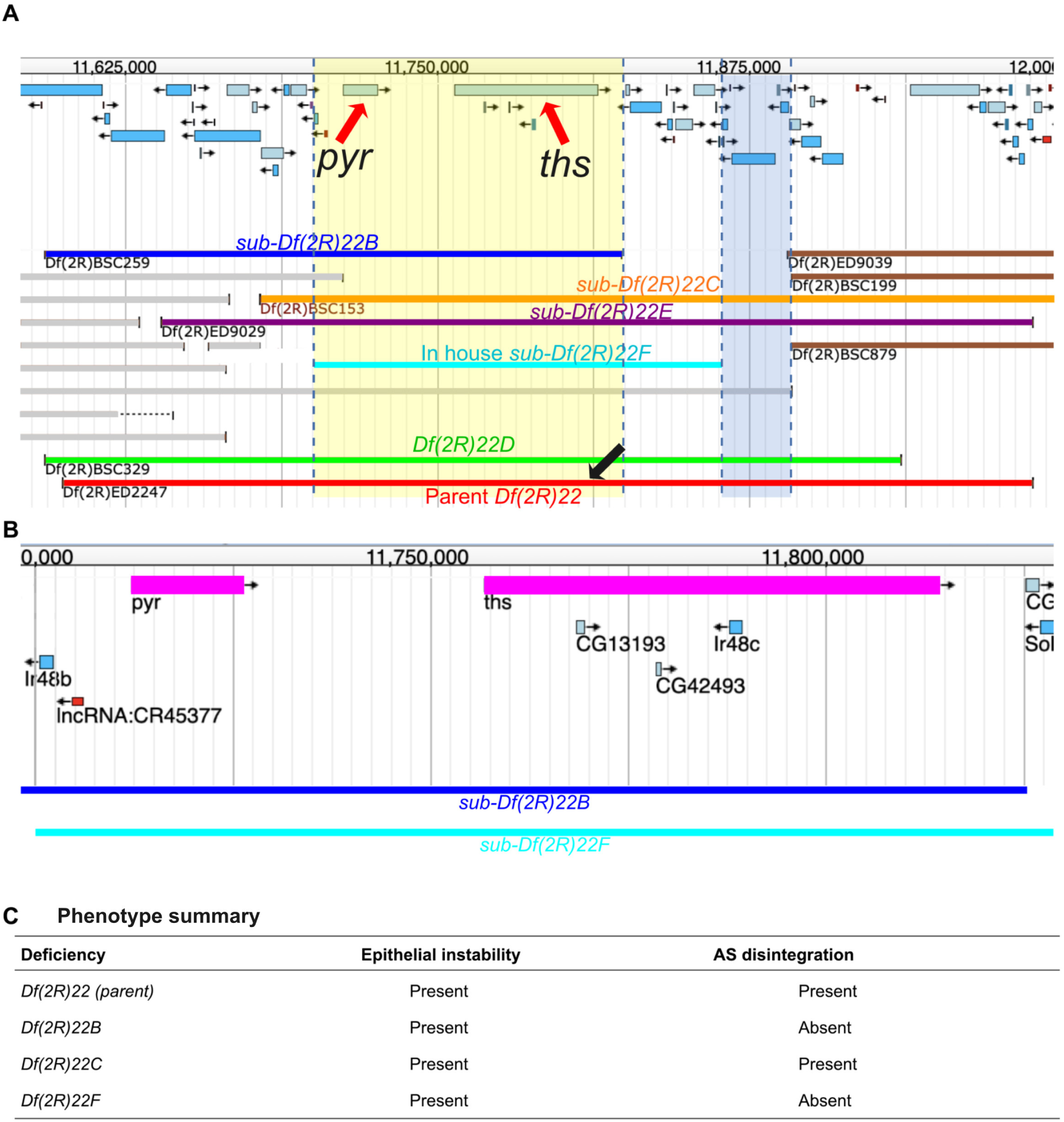
FlyBase view of the genes and deficiencies near *Df(2R)22* that mapped separate intervals responsible for epithelial instability and amnioserosa disintegration. (A) Parental *Df(2R)22* and overlapping sub-deficiencies. Yellow shading marks the interval responsible for epithelial-instability (contains *pyr* and *ths*). Blue marks the separate region that is apparently associated with amnioserosa disintegration. Dashed lines delimit these regions. The 22F endpoint is schematic rather than a molecularly defined breakpoint. (B) Enlarged view of the adjacent *pyr* and *ths* transcription units within the yellow region, with sub-*Df(2R)22B* and sub-*Df(2R)22F* shown below the genes. Coordinates specify location on 2R as per FlyBase. (C) Phenotype summary: 22B and 22F cause epithelial instability without amnioserosa disintegration, whereas 22 and 22C retain both.

**Figure S4.**
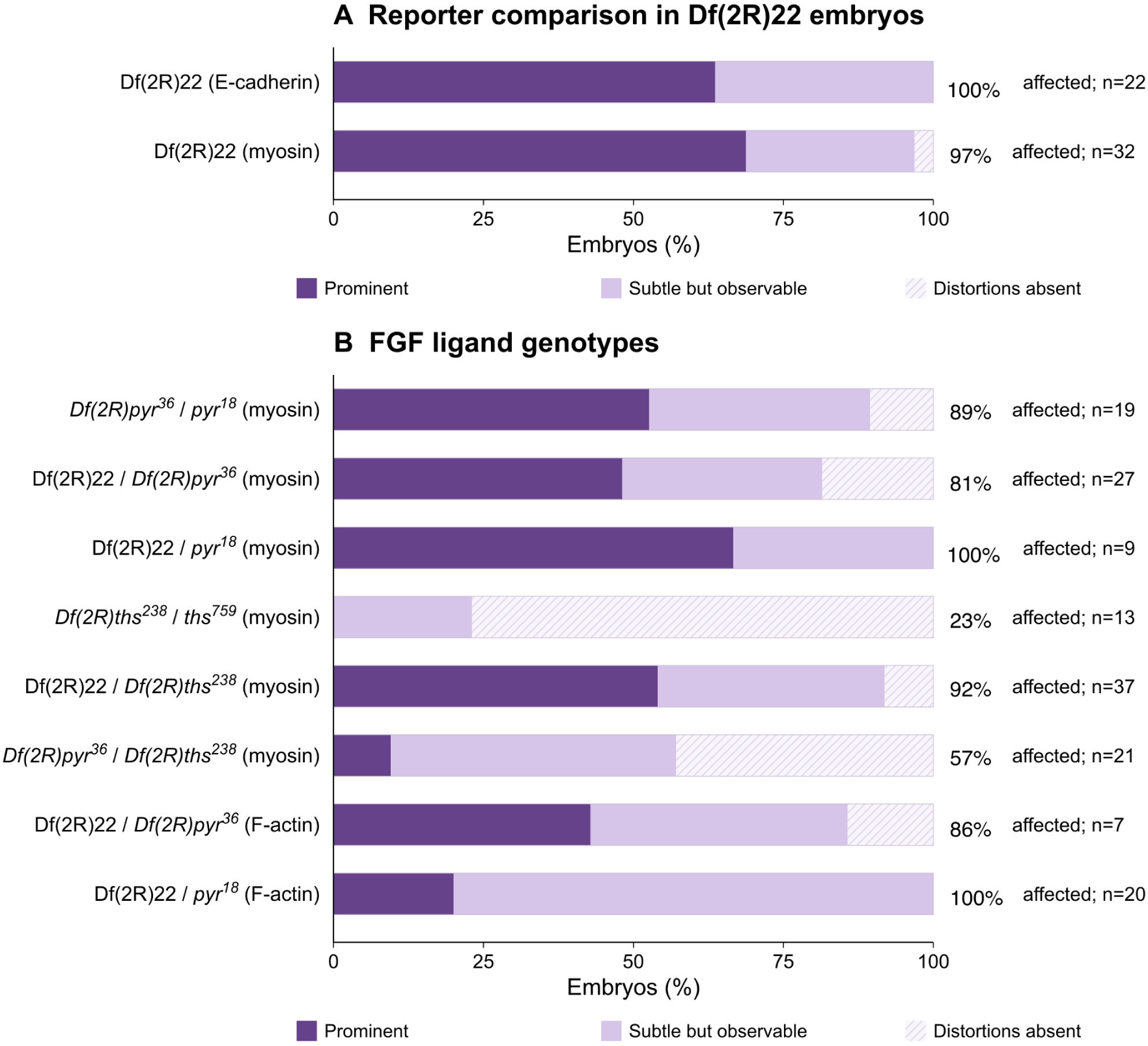
Penetrance and expressivity across deficiency and ligand genotypes. FGF ligand genotypes and fluorescent reporters are shown to the left of the penetrance/expressivity bars. (A) *Df(2R)22* embryos imaged with E-cadherin or myosin reporters. (B) Independent *pyr* and *ths* allele combinations with myosin or F-actin reporters for *in vivo* imaging. Dark and light segments show prominent versus subtle but observable distortions, respectively. Segment heights show the percentages of scored embryos that displayed a prominent or subtle phenotype. The total affected percentage and the number of embryos scored are printed to the right of each bar. Unaffected embryos account for the remainder to 100%. Counts are listed in Table S4; genotype and reporter cohorts are shown separately.

**Figure S5.**
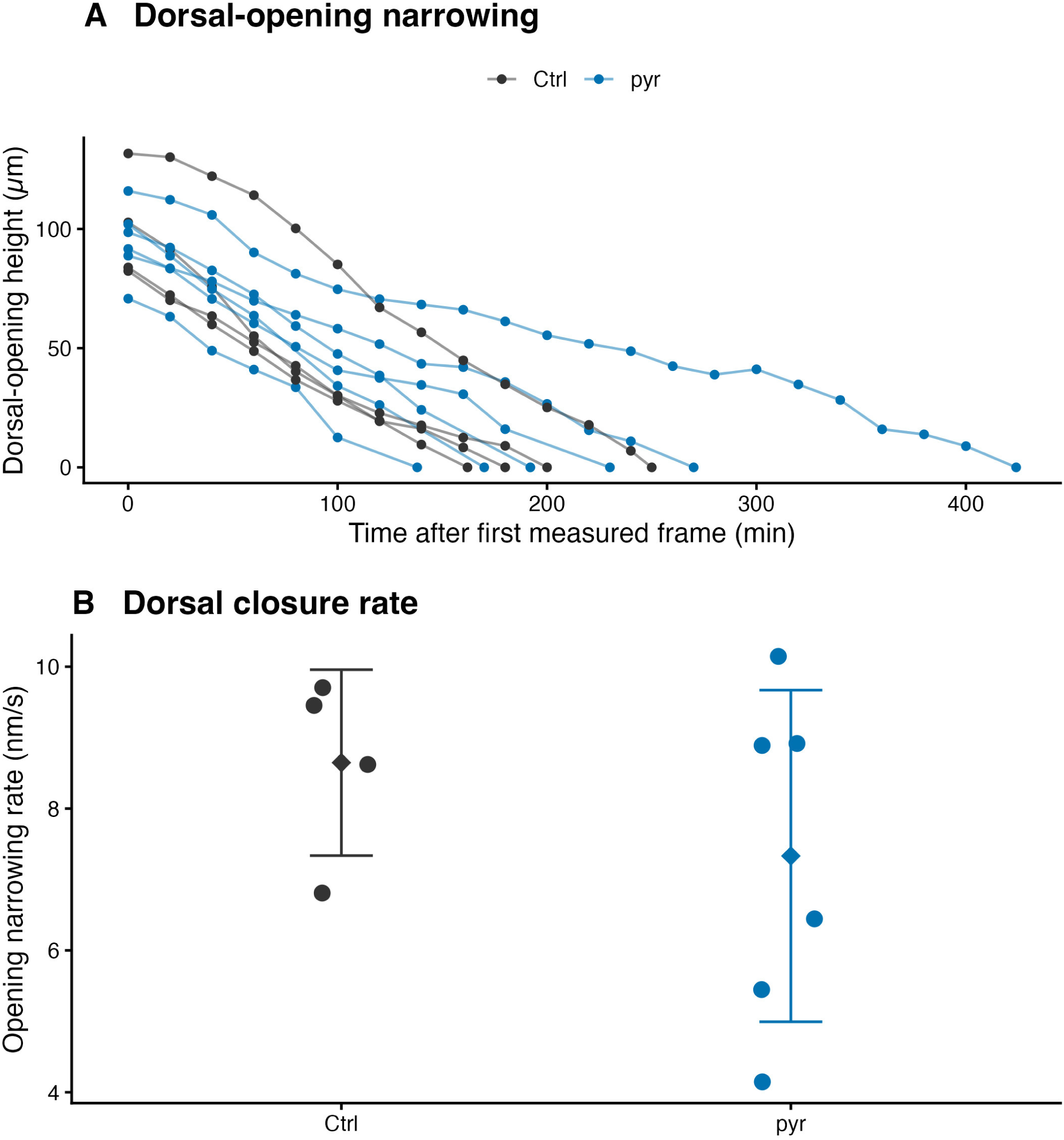
Dorsal-opening narrowing and closure rates in control and *pyr* embryos. (A) Dorsal-opening height versus time; lines connect measurements within embryos. Time is measured from each embryo’s first observation. (B) Narrowing rates estimated by ordinary least-squares regression of dorsal-opening height against time, with one slope per embryo (FIG. S6). Positive values indicate narrowing. Points represent embryos; diamonds and whiskers show means ± s.d. Control n=4; *pyr* n=6. Mean rates were 9 ± 1 and 7 ± 2 nm/s, respectively (Holm-adjusted P=0.289 across four mutant–control comparisons).

**Figure S6.**
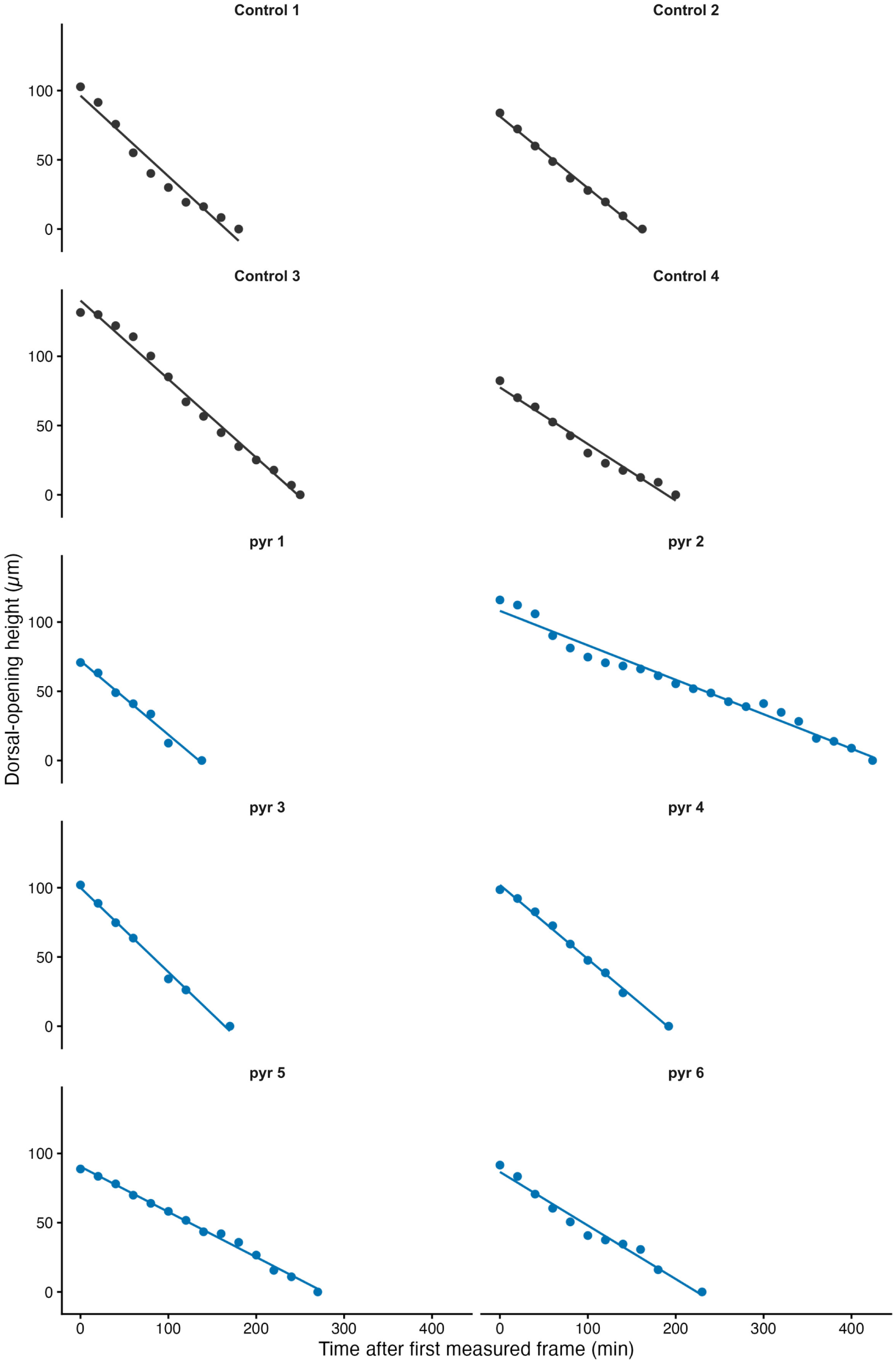
Individual embryo closure-rate fits for control and *pyr*. Points show dorsal-opening height measurements and lines show ordinary least-squares fits. Control n=4; *pyr* n=6. These are the observations summarized in FIG. S5. Narrowing rate is the negative slope of dorsal-opening height against time, expressed in nm/s.

**Figure S7.**
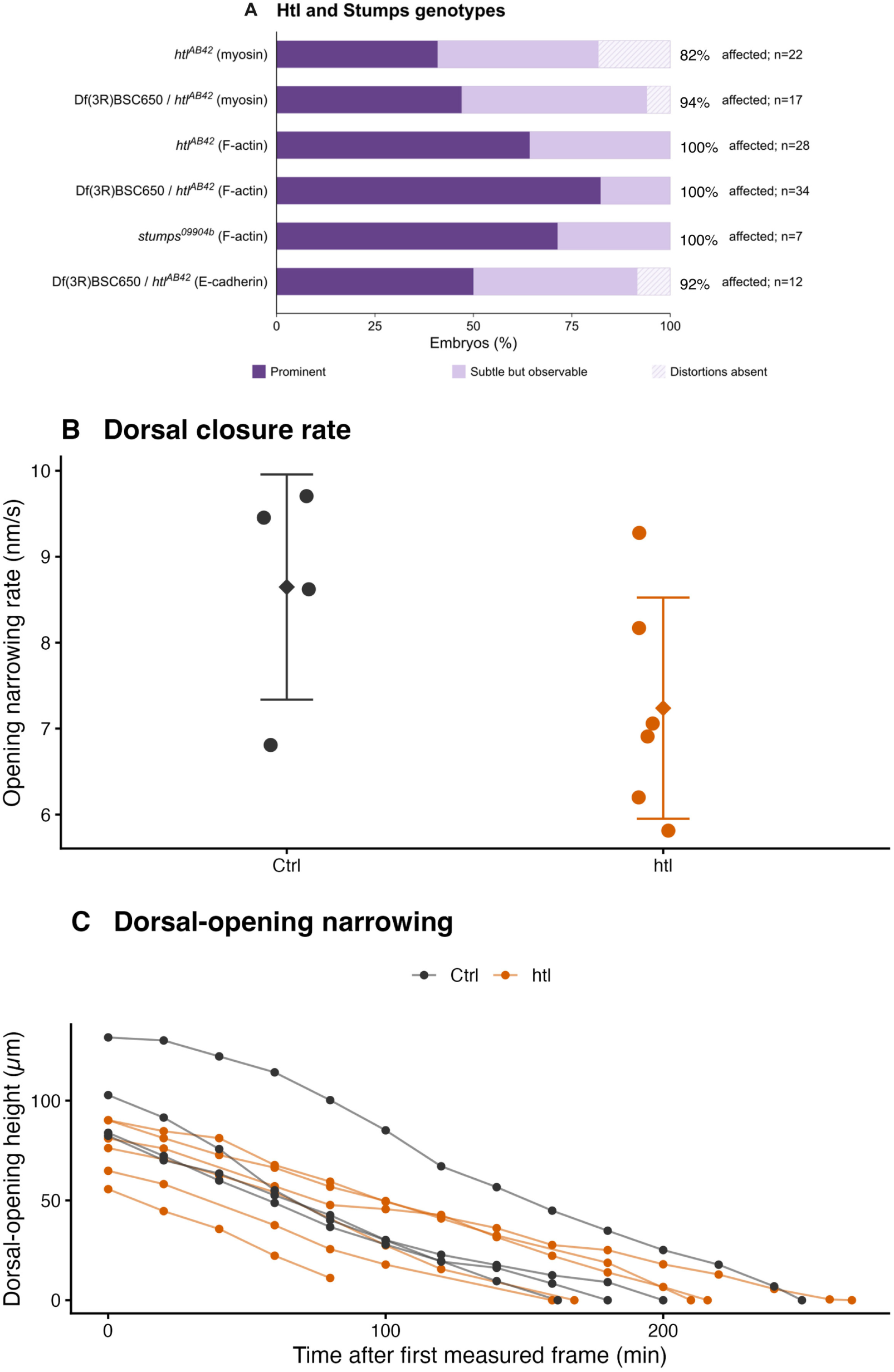
Penetrance, expressivity and closure rate in Htl and Stumps mutants. (A) Penetrance and expressivity for *htl* and *stumps*; dark purple, prominent distortions; light purple, subtle but observable distortions; pale patterned segments, distortions absent. See FIG. S4. (B) Independent opening-height measurements yielded one ordinary least-squares narrowing rate per embryo. Points represent embryos; diamonds and whiskers show means ± s.d. Control n=4; *htl* n=6. Mean rates were 9 ± 1 and 7 ± 1 nm/s, respectively (Holm-adjusted P=0.281 across four mutant–control comparisons). Htl embryos used an F-actin reporter; controls used Sqh. (C) Dorsal-opening height versus time, with lines connecting measurements within each embryo. Time begins at each embryo’s first measurement.

**Figure S8.**
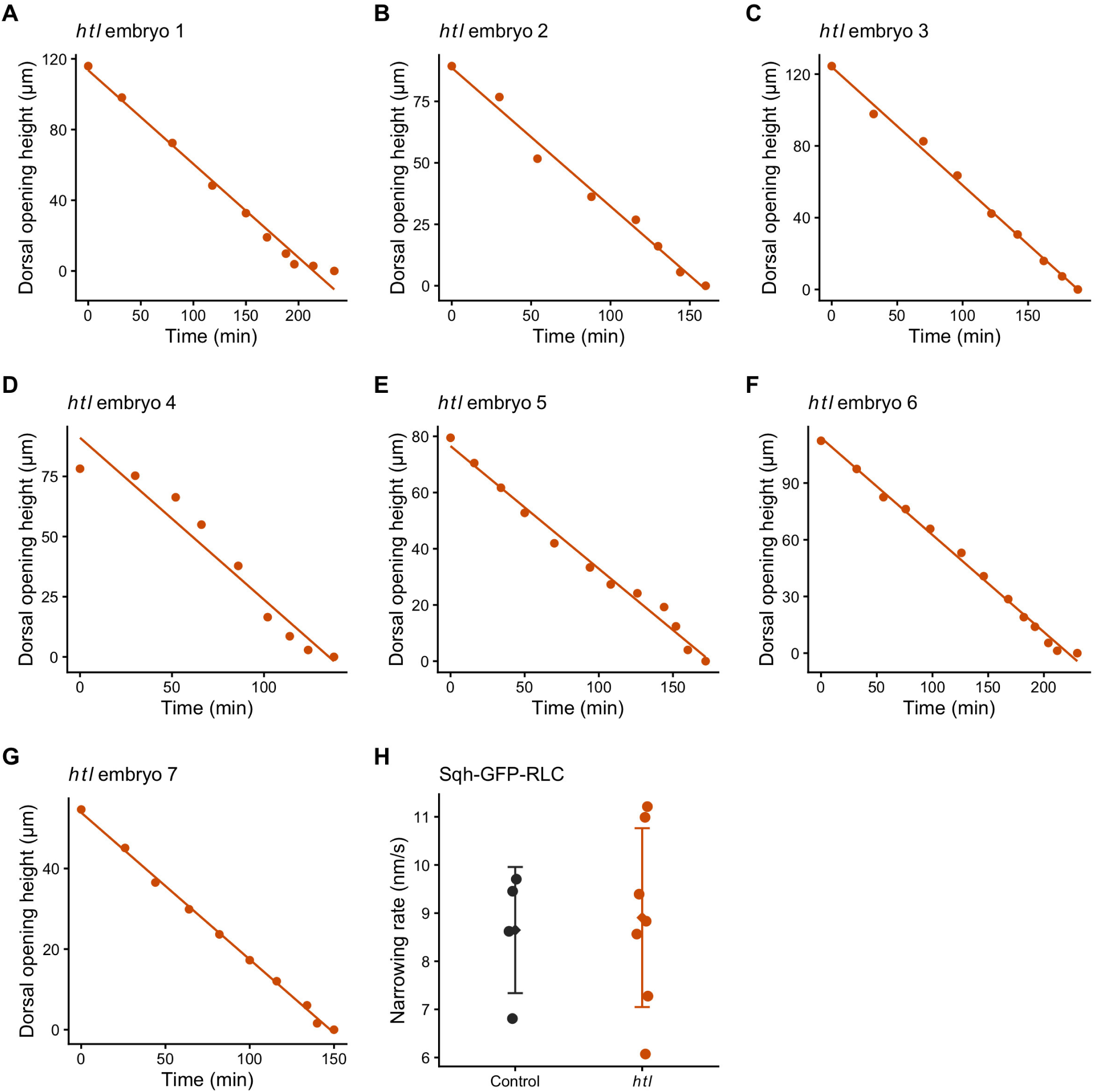
Dorsal opening narrowing in myosin-labelled *htl* embryos. (A–G) Dorsal opening height versus elapsed time for seven *htl* embryos expressing Sqh-GFP-RLC. Points show measurements; lines show ordinary least-squares fits for individual embryos. (H) Embryo-level narrowing rates for Sqh-GFP-RLC-labelled controls (n=4) and *htl* mutants (n=7). Circles represent individual embryos; diamonds and error bars indicate mean±s.d. Mean rates were 9±1 nm/s for controls and 9±2 nm/s for *htl* mutants (two-sided Welch t-test, t=−0.269, d.f.=8.35, P=0.795).

**Figure S9.**
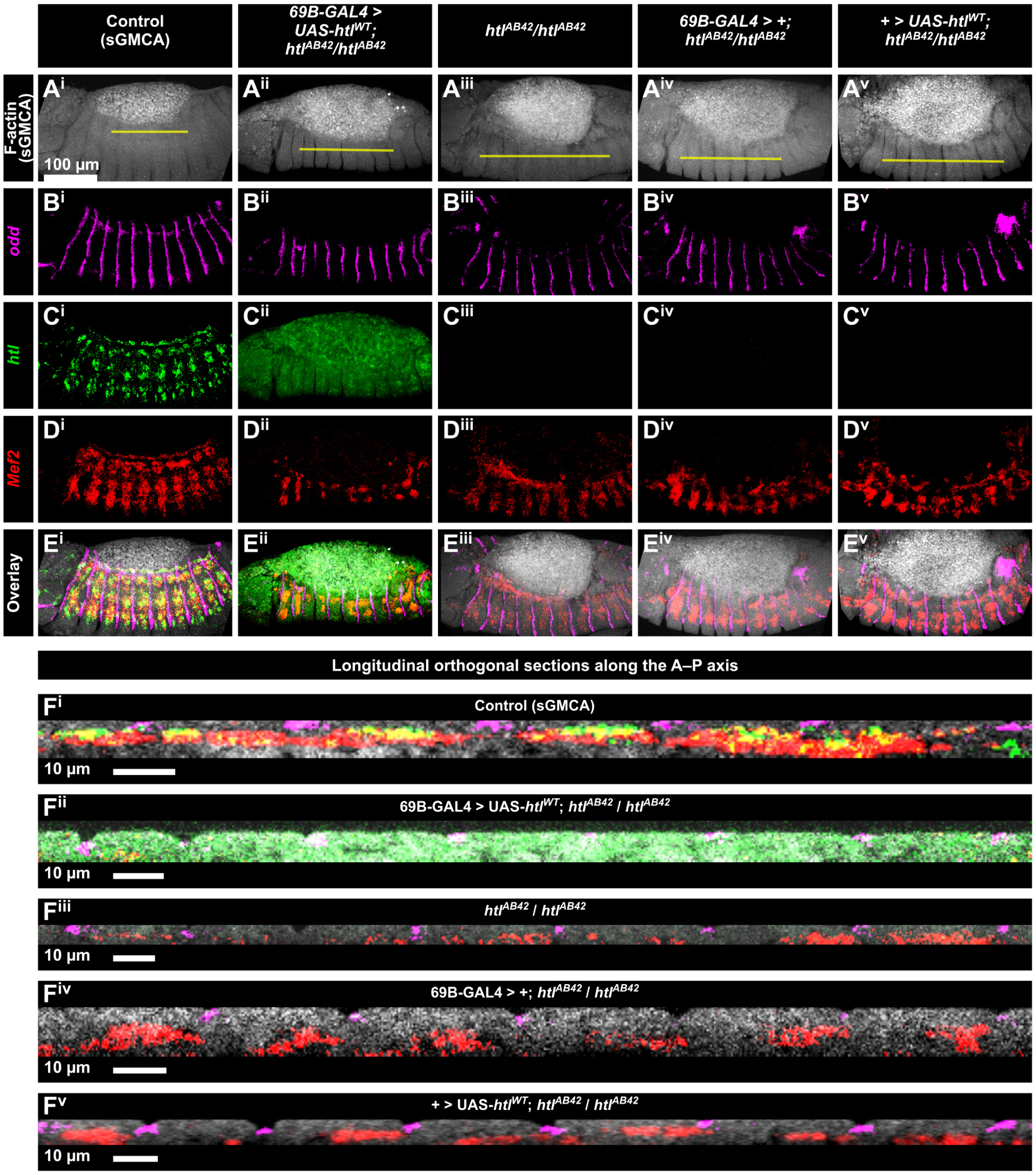
HCR-FISH validates tissue specific *htl* expression. Columns i–v show the genotypes indicated above the images: sGMCA control, *69B-GAL4*-driven *UAS-htl^WT^*expression in an *htl* mutant background, *htl* mutant, GAL4-only control and UAS-only control. Rows show the following. sGMCA fluorescence which is ubiquitously expressed (A^i^-A^v^). odd RNA which is an ectodermal marker, (B^i^-B^v^ and F^i^-F^v^, magenta). *htl* RNA, normally mesodermal (C^i^ and F^i^, green), also appears in the ectoderm when driven by *69B-GAL4* (C^ii^, E^ii^ and F^ii^, green). Mef2 RNA, mesodermal (D^i^-D^v^ and F^i^, F^iii^-F^v^, red) and overlays (E^i^-E^v^ and F^i^). Yellow lines in A mark the longitudinal positions used for the orthogonal sections in F^i^–F^v^; roman indices match the genotype columns. Ectodermal *htl* signal is visible in column ii, including C^ii^, E^ii^ and F^ii^. Scale bars, 100 *μ*m in overview images and 10 *μ*m in orthogonal sections.

**Figure S10.**
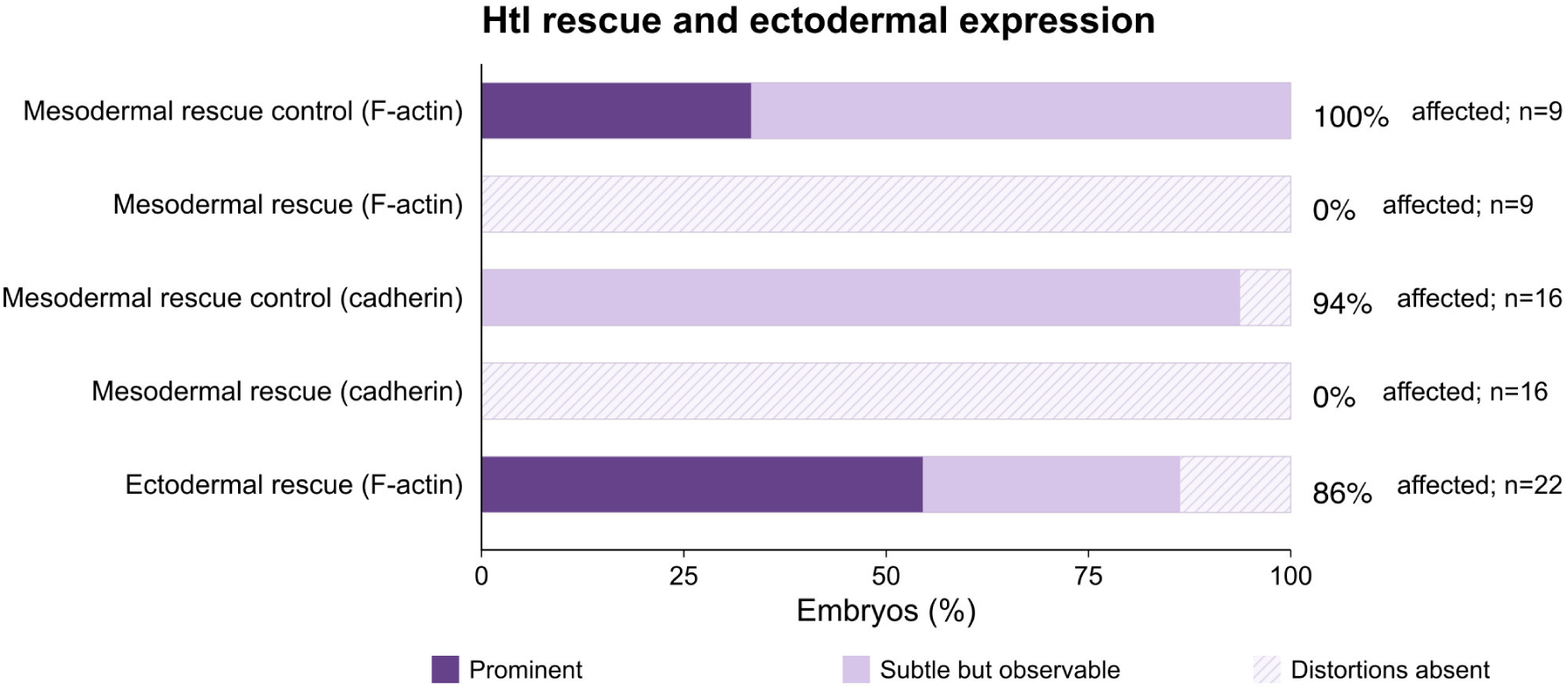
Penetrance and expressivity in *htl* mutant embryos following mesodermal or ectodermal Htl expression. See legend for FIG. S4. Bars show the penetrance and expressivity of prominent or subtle but observable phenotypes in *htl* mutant embryos. Mesodermal, but not ectodermal expression rescues these phenotypes.

**Figure S11.**
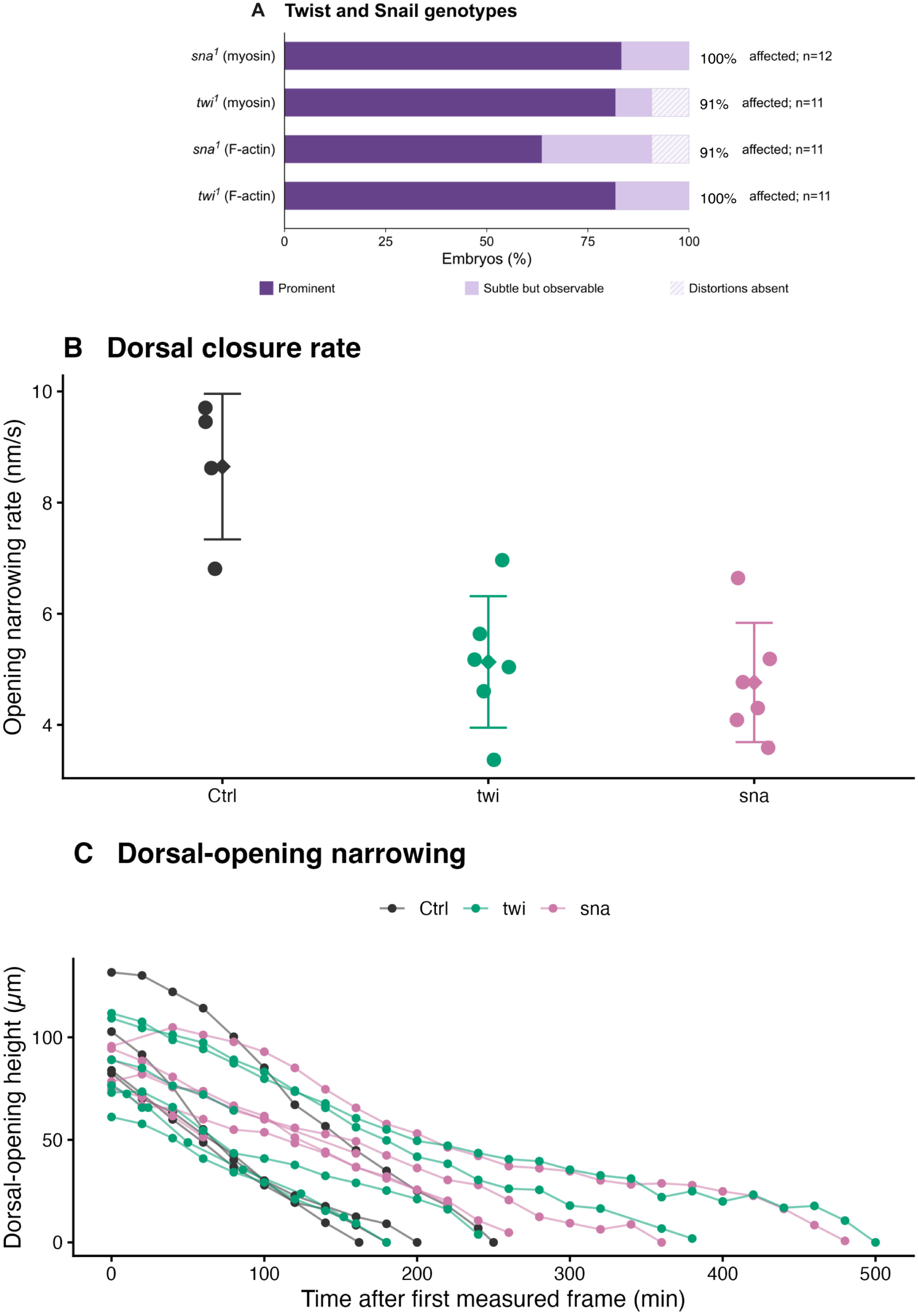
Penetrance, expressivity and closure rates in twist and snail mutants. (A) Penetrance and expressivity for *sna* and *twi*; dark purple, prominent distortions; light purple, subtle but observable distortions; pale patterned segments, distortions absent. See FIG. S4. (B) Independent opening-height measurements yielded one ordinary least-squares narrowing rate per embryo. Points represent embryos; diamonds and whiskers show means ± s.d. Control n=4; *twi* and *sna* n=6 each. Mean rates were 9 ± 1, 5.1 ± 1.2 and 5 ± 1 nm/s, respectively. Twi and *sna* rates were lower than control (Holm-adjusted P=0.0146 and 0.0127 across four mutant–control comparisons). (C) Dorsal-opening height versus time, with lines connecting measurements within each embryo. Time begins at each embryo’s first measurement.

**Figure S12.**
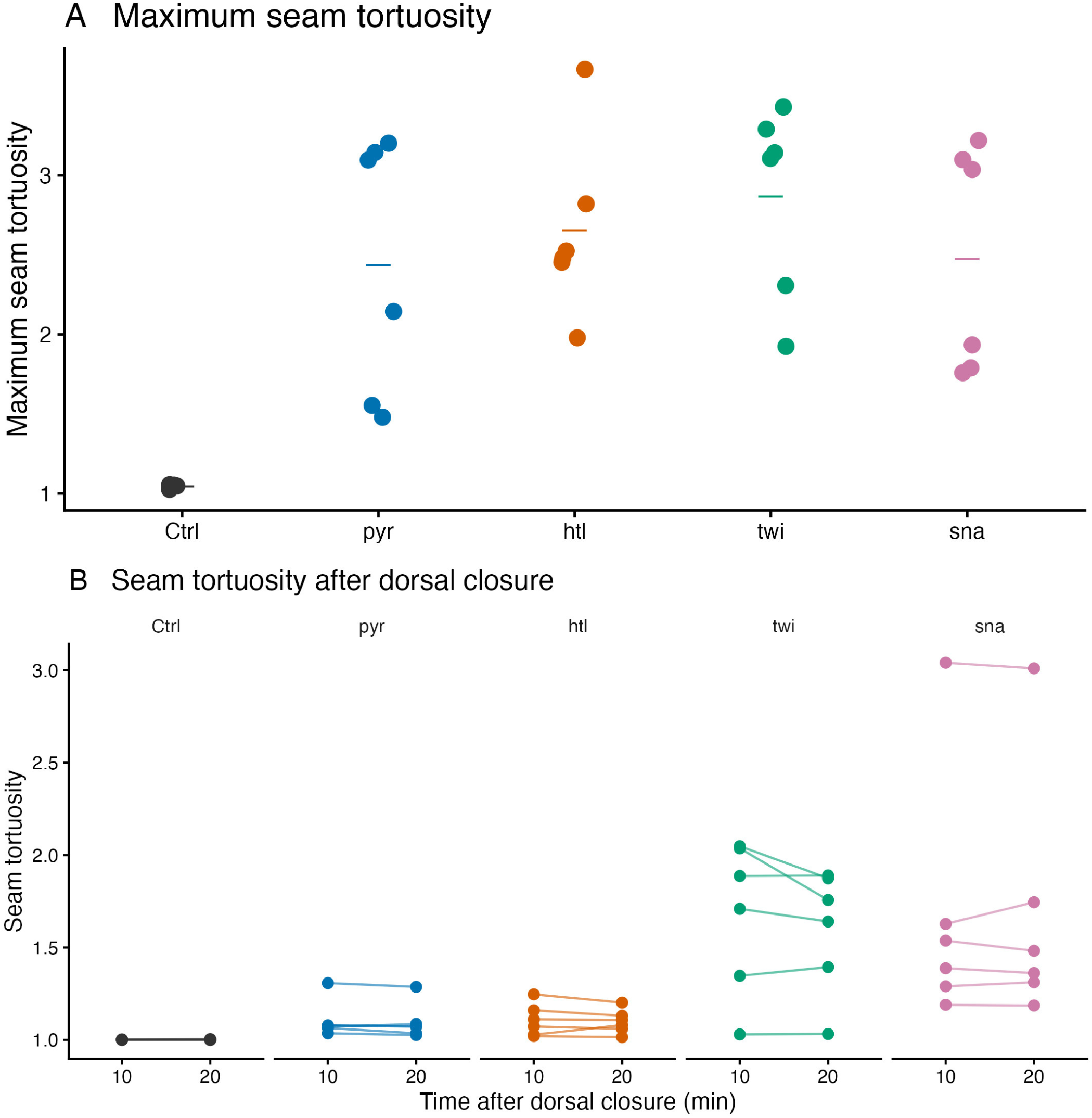
Maximum seam tortuosity and seam tortuosity after dorsal closure. (A) Maximum observed tortuosity per embryo. Points represent embryos; horizontal bars show group means. (B) Seam tortuosity at 10 and 20 min after dorsal closure; lines connect the two observations from each embryo. Control n=4; *pyr*, *htl*, *twi* and *sna* n=6 each. Tortuosity is seam contour length divided by straight-line endpoint distance. Figure 7L summarizes each embryo’s mean across the two post-closure times.

**Figure S13.**
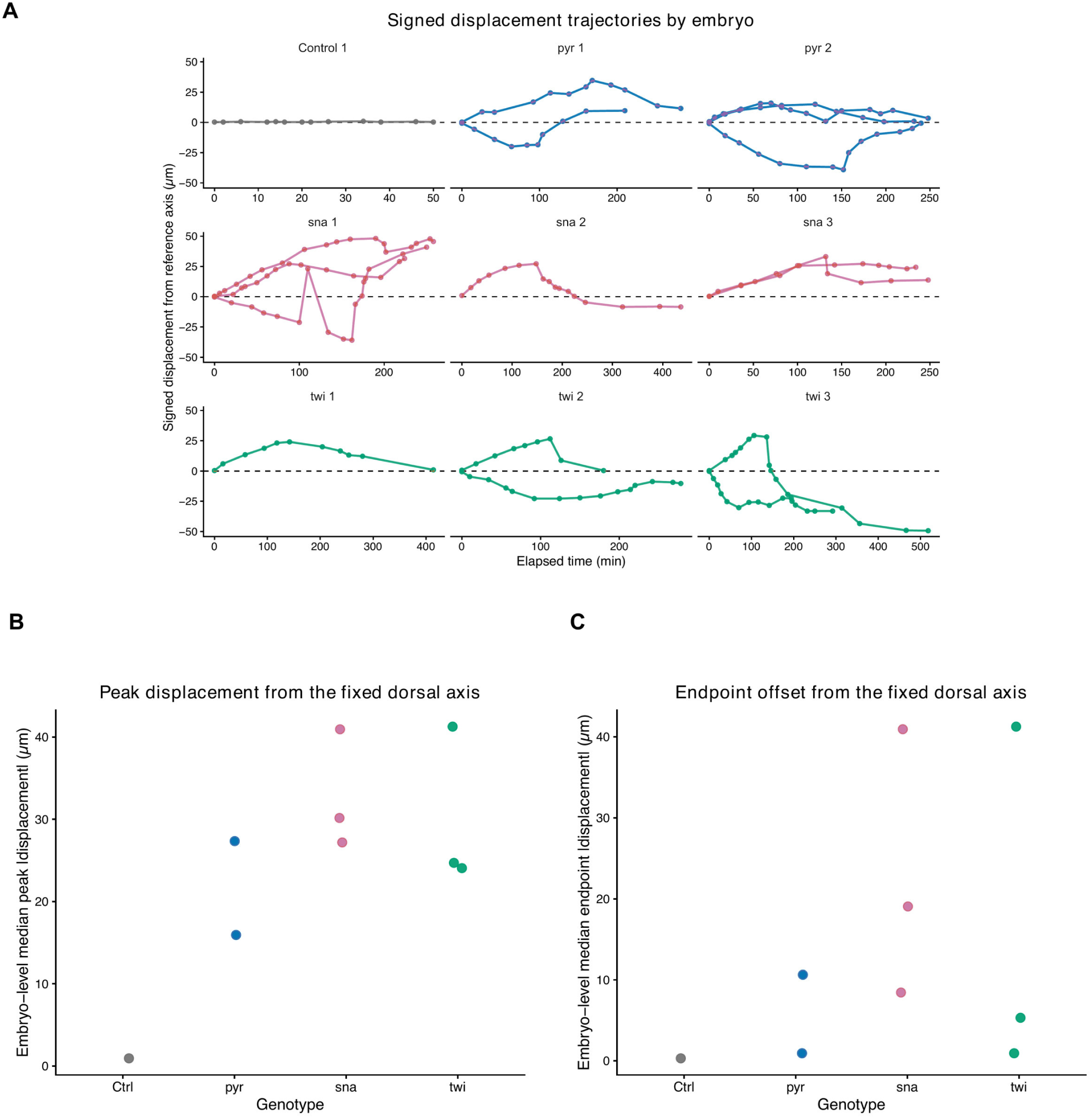
Individual seam displacement trajectories as a function of time. Time 0 marks the beginning of an experimental run. Perpendicular displacement of local distortions from a fixed reference axis, shown separately for each embryo as a function of time. Each plot is for a single embryo, multiple tracks on a plot indicate that more than one deviation was measured. Negative values indicate the displacement was below the dorsal midline of an embryo. (B,C) Each point is one embryo: the median across its tracked distortions of peak absolute displacement (B) or final absolute offset (C).

**Figure S14.**
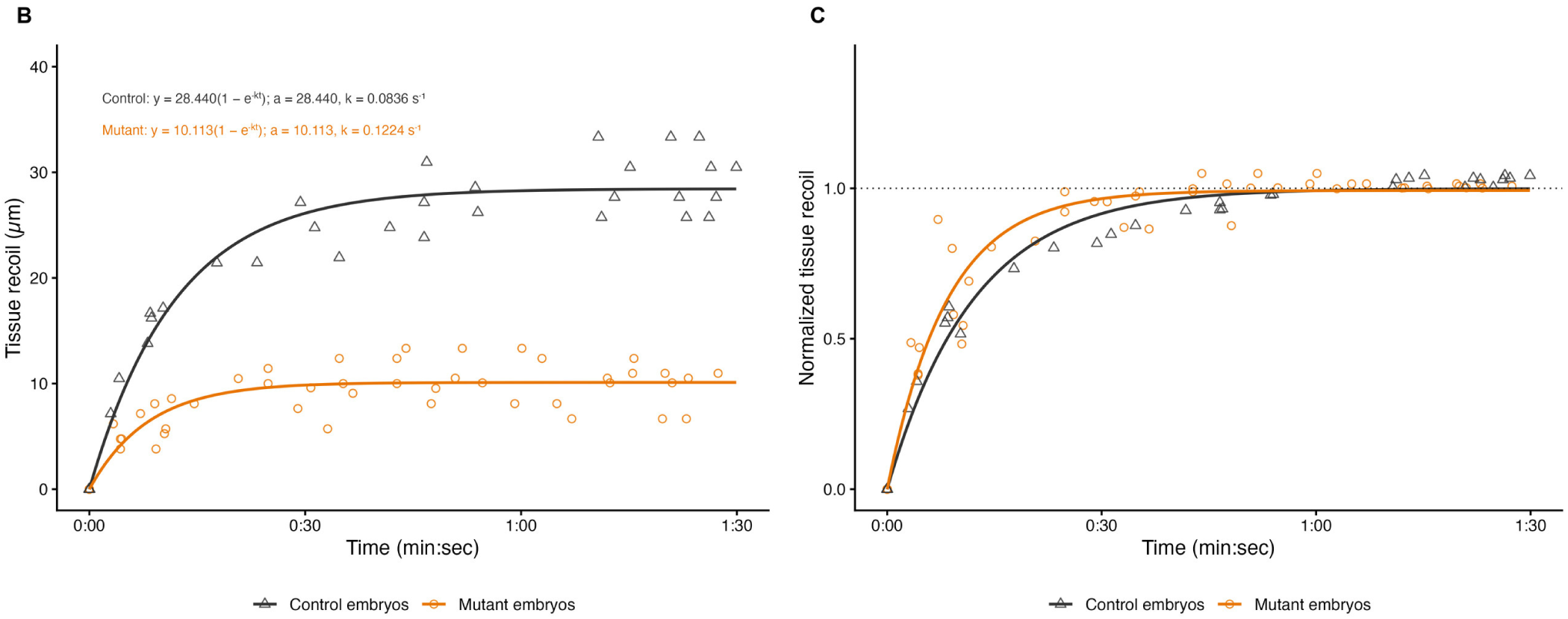
Recoil following ASPS surgical cuts perpendicular to the anterior–posterior axis, severing one or two rows of lateral epidermal cells and the amnioserosa. (Fig. 9D–I). (B) Raw displacement and exponential fits D(t)=a(1−exp(−kt)). (C) Normalized trajectories for control and mutant embryos obtained by dividing each trajectory by its individually fitted plateau, then fitting each normalized curve. Three control and six *htl* embryos contributed four and eight tracks, respectively, with one or two markers followed per cut. Statistics compare parameters fitted separately to raw tracks then averaged by embryo. Plateau P = 1.2 × 10^-5^ and rate constant P = 0.16 (two-sided Welch t-tests). Time is in min:sec. Normalization is for display and does not introduce a separate rate test; estimates and sample sizes are in Table S3.

**Figure S15.**
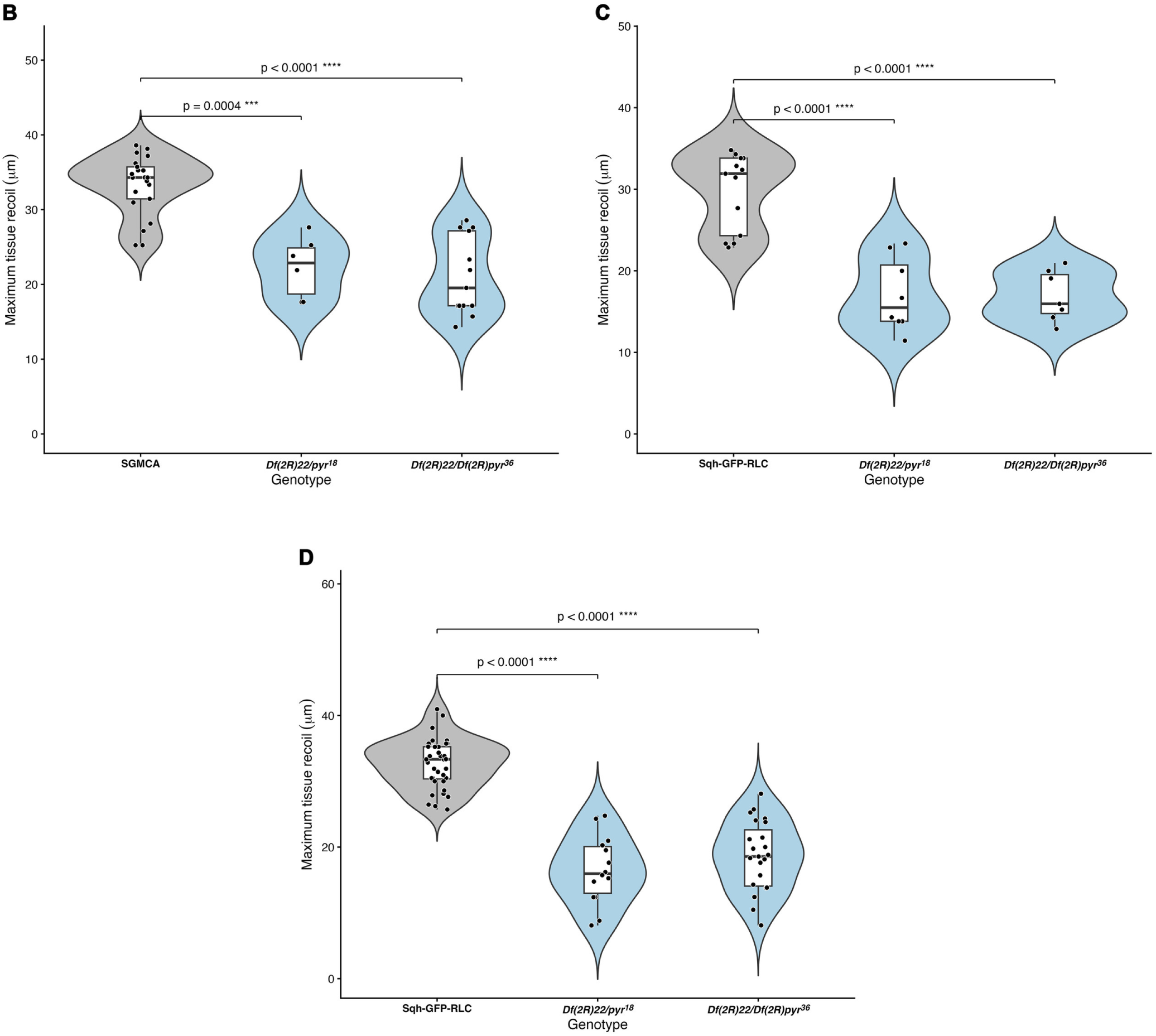
Maximum recoil distributions for control and *pyr* mutant allele combinations labeled with Sqh-GFP-RLC or sGMCA. (B–D) Maximum tissue recoil for control and *pyr* mutant combinations, measured with sGMCA after ASPS cuts (B), Sqh-GFP-RLC after amnioserosa cuts (C), and Sqh-GFP-RLC after ASPS cuts (D). Points represent measurements; boxes show the median and interquartile range, with whiskers extending to 1.5 times the interquartile range. Graphs show unadjusted two-sided Welch-test P values; adjusted P values and sample sizes are in Table S3.

**Figure S16.**
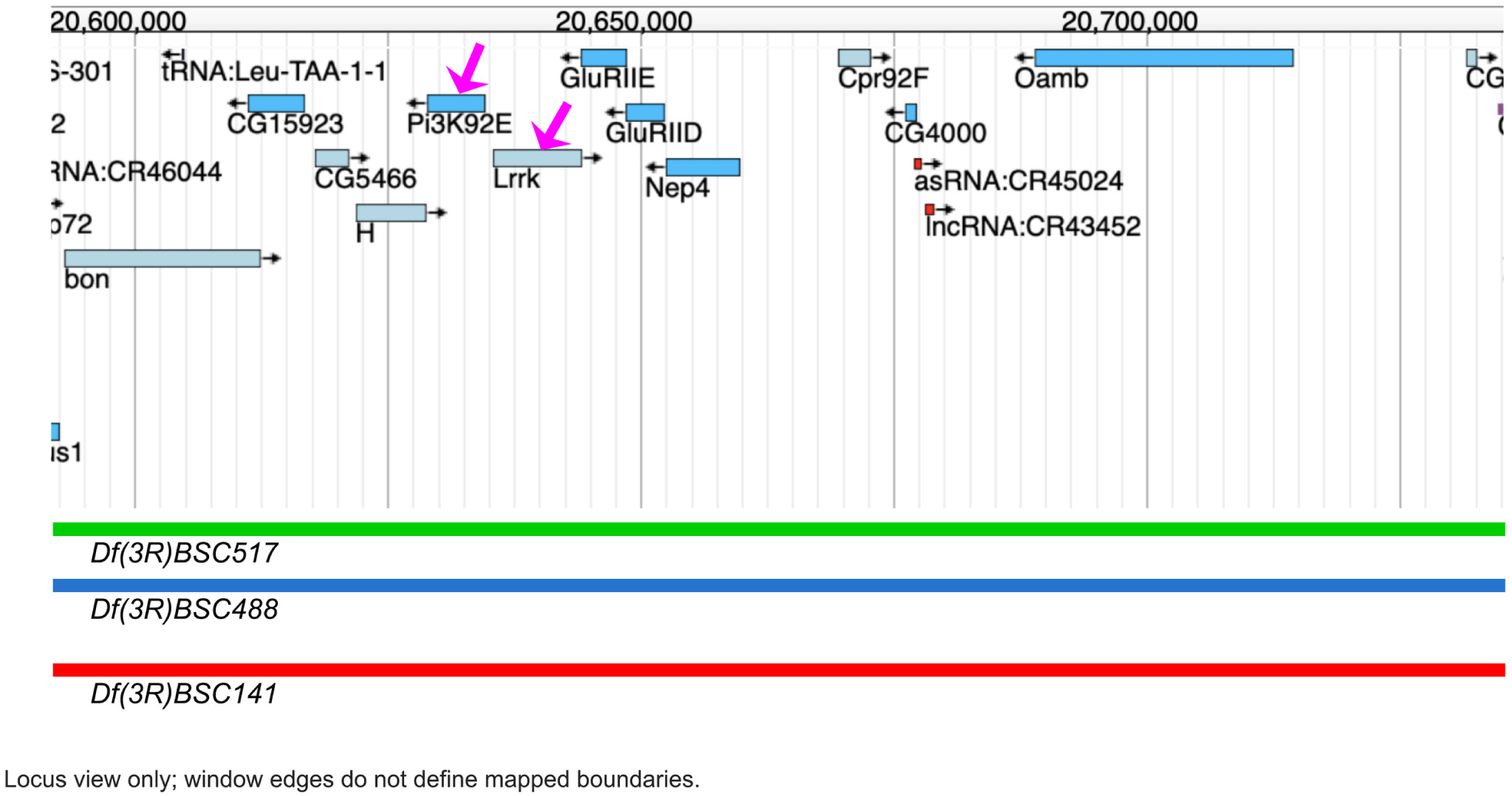
Genomic context of the *Pi3K92E* and *Lrrk* candidates on a map redrawn from FlyBase. Magenta arrows identify *Pi3K92E* and *Lrrk*. The displayed portions of three overlapping *Dfs* and their names in FlyBase are shown. Each *Df* extends beyond edge of the figure. *Df(3R)del6* and *Df(3R)GluRIID[attP]* are omitted. Representative micrographs are presented in main Fig. 10 where phenotypes for different genotypes are shown.

**Table S1.**
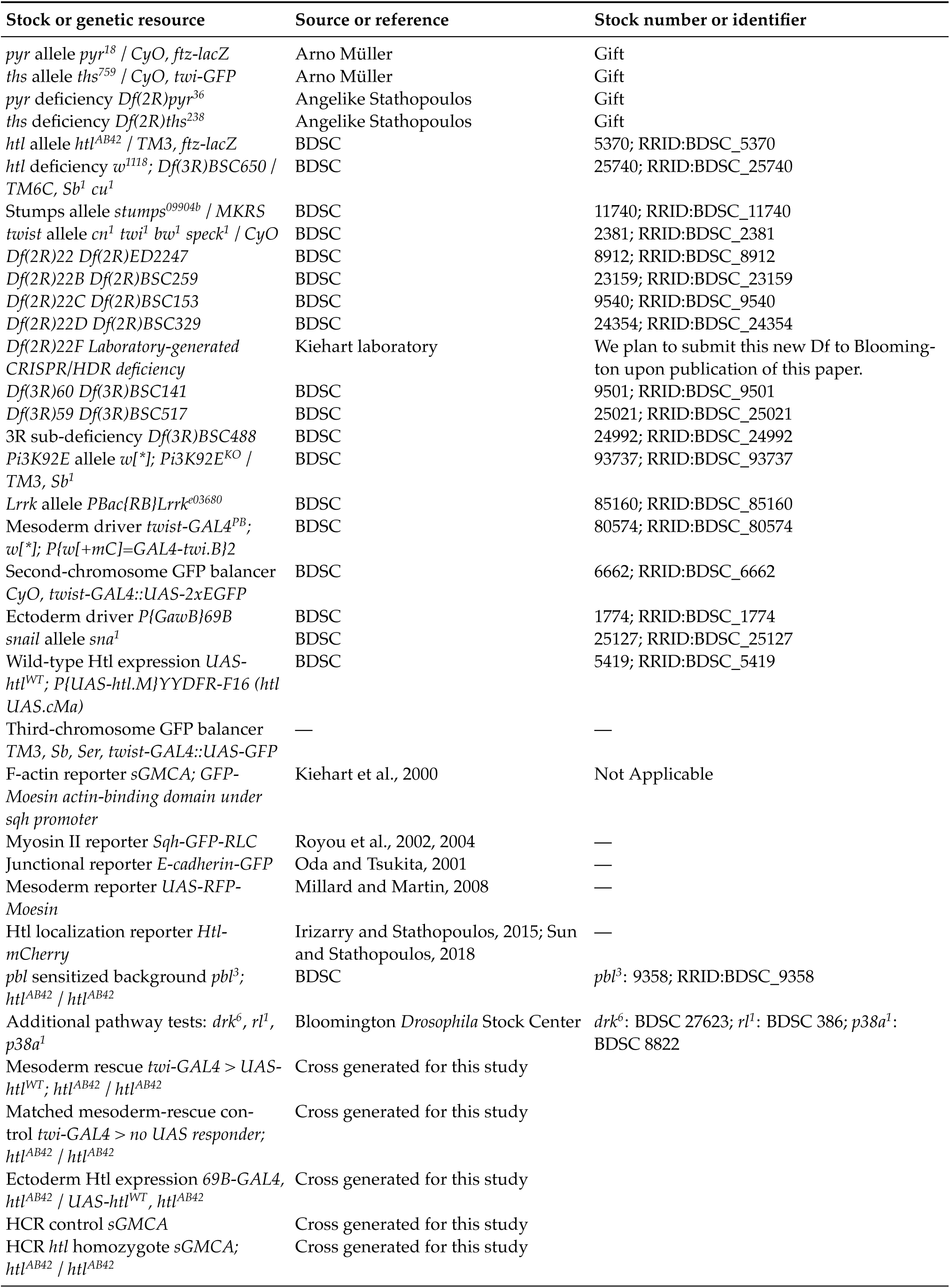
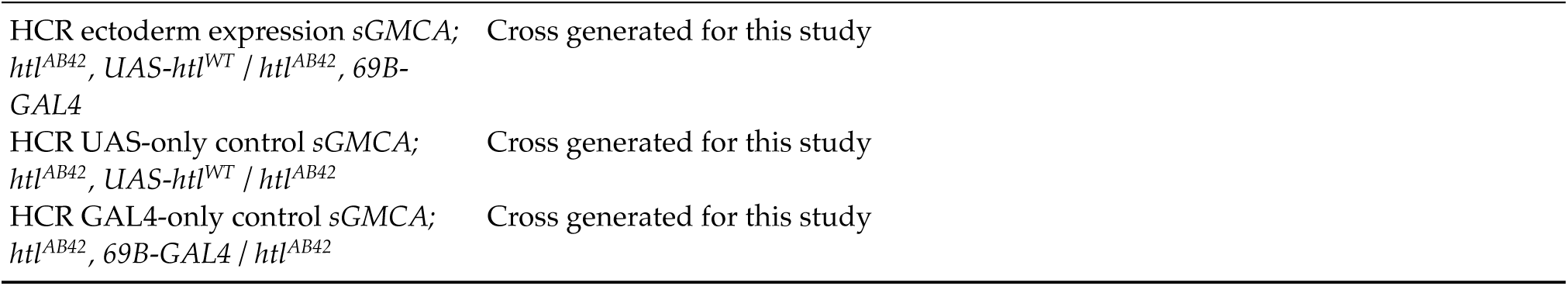
*Drosophila* stocks and genetic resources. BDSC, Bloomington *Drosophila* Stock Center; N/A, not applicable. Stock numbers identify catalogued alleles or transgenic insertions; laboratory crosses and recombined chromosomes are listed separately. An em dash indicates unavailable information.

| Stock or genetic resource | Source or reference | Stock number or identifier |
|---|---|---|
| *pyr* allele *pyr<sup>18</sup>* / *CyO, ftz-lacZ* | Arno Müller | Gift |
| *ths* allele *ths<sup>759</sup>* / *CyO, twi-GFP* | Arno Müller | Gift |
| *pyr* deficiency *Df(2R)pyr<sup>36</sup>* | Angelike Stathopoulos | Gift |
| *ths* deficiency *Df(2R)ths<sup>238</sup>* | Angelike Stathopoulos | Gift |
| *htl* allele *htl<sup>AB42</sup>* / *TM3, ftz-lacZ* | BDSC | 5370; RRID:BDSC_5370 |
| *htl* deficiency *w<sup>1118</sup>; Df(3R)BSC650* / *TM6C, Sb<sup>1</sup> cu<sup>1</sup>* | BDSC | 25740; RRID:BDSC_25740 |
| Stumps allele *stumps<sup>09904b</sup>* / *MKRS* | BDSC | 11740; RRID:BDSC_11740 |
| *twist* allele *cn<sup>1</sup> twi<sup>1</sup> bw<sup>1</sup> speck<sup>1</sup>* / *CyO* | BDSC | 2381; RRID:BDSC_2381 |
| *Df(2R)22 Df(2R)ED2247* | BDSC | 8912; RRID:BDSC_8912 |
| *Df(2R)22B Df(2R)BSC259* | BDSC | 23159; RRID:BDSC_23159 |
| *Df(2R)22C Df(2R)BSC153* | BDSC | 9540; RRID:BDSC_9540 |
| *Df(2R)22D Df(2R)BSC329* | BDSC | 24354; RRID:BDSC_24354 |
| *Df(2R)22F Laboratory-generated CRISPR/HDR deficiency* | Kiehart laboratory | We plan to submit this new Df to Blooming-ton upon publication of this paper. |
| *Df(3R)60 Df(3R)BSC141* | BDSC | 9501; RRID:BDSC_9501 |
| *Df(3R)59 Df(3R)BSC517* | BDSC | 25021; RRID:BDSC_25021 |
| 3R sub-deficiency *Df(3R)BSC488* | BDSC | 24992; RRID:BDSC_24992 |
| *Pi3K92E* allele *w[\*]; Pi3K92E<sup>KO</sup>* / *TM3, Sb<sup>1</sup>* | BDSC | 93737; RRID:BDSC_93737 |
| *Lrrk* allele *PBac{RB}Lrrk<sup>e03680</sup>* | BDSC | 85160; RRID:BDSC_85160 |
| Mesoderm driver *twist-GAL4<sup>PB</sup>*; *w[\*]; P{w[+mC]=GAL4-twi.B}2* | BDSC | 80574; RRID:BDSC_80574 |
| Second-chromosome GFP balancer *CyO, twist-GAL4::UAS-2xEGFP* | BDSC | 6662; RRID:BDSC_6662 |
| Ectoderm driver *P{GawB}69B* | BDSC | 1774; RRID:BDSC_1774 |
| *snail* allele *sna<sup>1</sup>* | BDSC | 25127; RRID:BDSC_25127 |
| Wild-type Htl expression *UAS-htl<sup>WT</sup>; P{UAS-htl.M}YYDFR-F16 (htl UAS.cMa)* | BDSC | 5419; RRID:BDSC_5419 |
| Third-chromosome GFP balancer *TM3, Sb, Ser, twist-GAL4::UAS-GFP* | — | — |
| F-actin reporter *sGMCA; GFP-Moesin actin-binding domain under sqh promoter* | Kiehart et al., 2000 | Not Applicable |
| Myosin II reporter *Sqh-GFP-RLC* | Royou et al., 2002, 2004 | — |
| Junctional reporter *E-cadherin-GFP* | Oda and Tsukita, 2001 | — |
| Mesoderm reporter *UAS-RFP-Moesin* | Millard and Martin, 2008 | — |
| Htl localization reporter *Htl-mCherry* | Irizarry and Stathopoulos, 2015; Sun and Stathopoulos, 2018 | — |
| *pbl* sensitized background *pbl<sup>3</sup>; htl<sup>AB42</sup>* / *htl<sup>AB42</sup>* | BDSC | *pbl<sup>3</sup>*: 9358; RRID:BDSC_9358 |
| Additional pathway tests: *drk<sup>6</sup>*, *rl<sup>1</sup>*, *p38a<sup>1</sup>* | Bloomington *Drosophila* Stock Center | *drk<sup>6</sup>*: BDSC 27623; *rl<sup>1</sup>*: BDSC 386; *p38a<sup>1</sup>*: BDSC 8822 |
| Mesoderm rescue *twi-GAL4* > *UAS-htl<sup>WT</sup>*; *htl<sup>AB42</sup>* / *htl<sup>AB42</sup>* | Cross generated for this study |  |
| Matched mesoderm-rescue con-trol *twi-GAL4* > *no UAS responder; htl<sup>AB42</sup>* / *htl<sup>AB42</sup>* | Cross generated for this study |  |
| Ectoderm Htl expression *69B-GAL4*, *htl<sup>AB42</sup>* / *UAS-htl<sup>WT</sup>*, *htl<sup>AB42</sup>* | Cross generated for this study |  |
| HCR control *sGMCA* | Cross generated for this study |  |
| HCR *htl* homozygote *sGMCA; htl<sup>AB42</sup>* / *htl<sup>AB42</sup>* | Cross generated for this study |  |
| HCR ectoderm expression *sGMCA; htl<sup>AB42</sup>*, *UAS-htl<sup>WT</sup>* / *htl<sup>AB42</sup>*, *69B-GAL4* | Cross generated for this study |  |
| HCR UAS-only control *sGMCA; htl<sup>AB42</sup>*, *UAS-htl<sup>WT</sup>* / *htl<sup>AB42</sup>* | Cross generated for this study |  |
| HCR GAL4-only control *sGMCA; htl<sup>AB42</sup>*, *69B-GAL4* / *htl<sup>AB42</sup>* | Cross generated for this study |  |

**Table S2.**
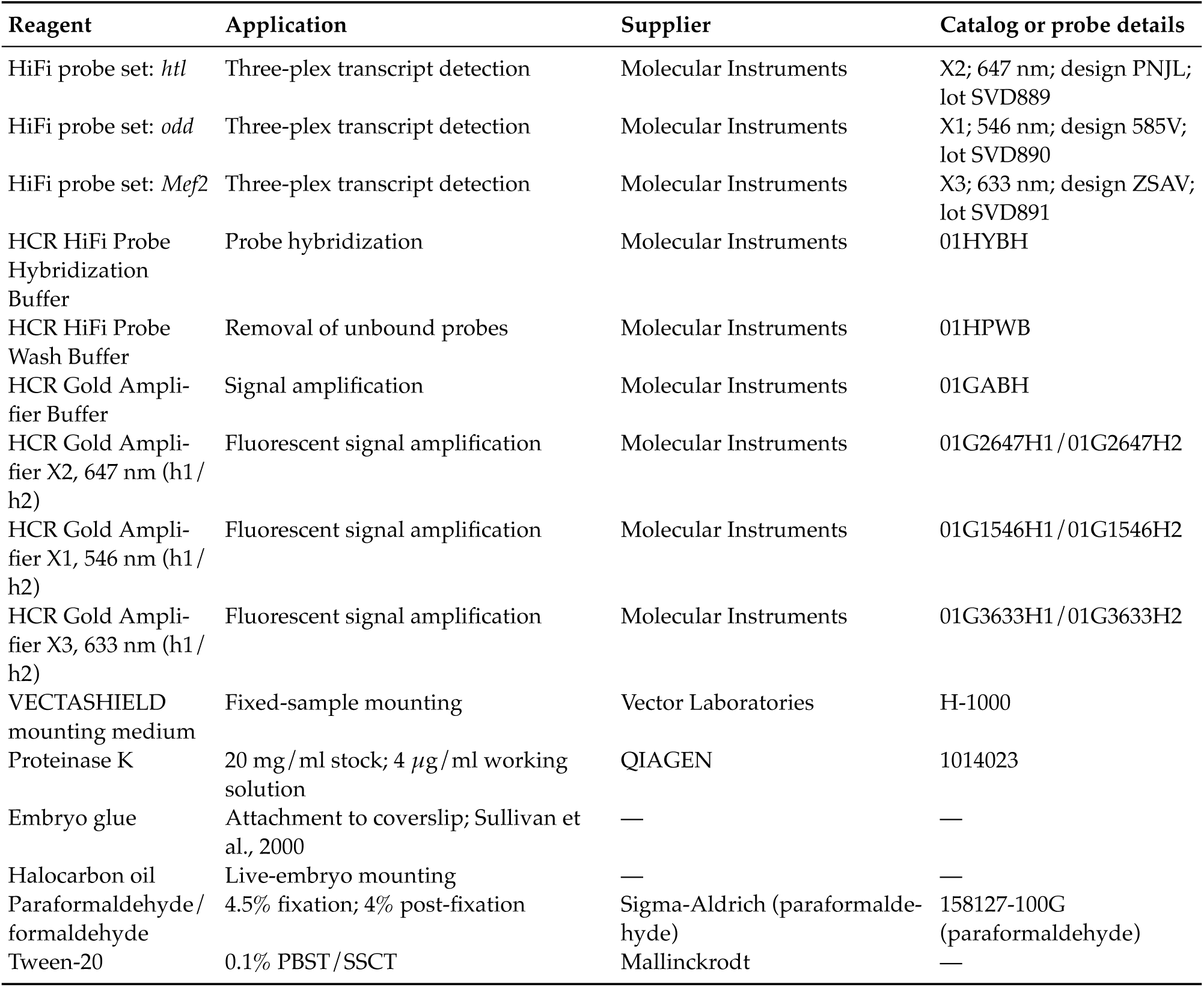
Reagents and RNA FISH probes.

| Reagent | Application | Supplier | Catalog or probe details |
|---|---|---|---|
| HiFi probe set: *htl* | Three-plex transcript detection | Molecular Instruments | X2; 647 nm; design PNJL; lot SVD889 |
| HiFi probe set: *odd* | Three-plex transcript detection | Molecular Instruments | X1; 546 nm; design 585V; lot SVD890 |
| HiFi probe set: *Mef2* | Three-plex transcript detection | Molecular Instruments | X3; 633 nm; design ZSAV; lot SVD891 |
| HCR HiFi Probe Hybridization Buffer | Probe hybridization | Molecular Instruments | 01HYBH |
| HCR HiFi Probe Wash Buffer | Removal of unbound probes | Molecular Instruments | 01HPWB |
| HCR Gold Ampli-fier Buffer | Signal amplification | Molecular Instruments | 01GABH |
| HCR Gold Ampli-fier X2, 647 nm (h1/h2) | Fluorescent signal amplification | Molecular Instruments | 01G2647H1/01G2647H2 |
| HCR Gold Ampli-fier X1, 546 nm (h1/h2) | Fluorescent signal amplification | Molecular Instruments | 01G1546H1/01G1546H2 |
| HCR Gold Ampli-fier X3, 633 nm (h1/h2) | Fluorescent signal amplification | Molecular Instruments | 01G3633H1/01G3633H2 |
| VECTASHIELD mounting medium | Fixed-sample mounting | Vector Laboratories | H-1000 |
| Proteinase K | 20 mg/ml stock; 4 µg/ml working solution | QIAGEN | 1014023 |
| Embryo glue | Attachment to coverslip; Sullivan et al., 2000 | — | — |
| Halocarbon oil | Live-embryo mounting | — | — |
| Paraformaldehyde/formaldehyde | 4.5% fixation; 4% post-fixation | Sigma-Aldrich (paraformalde-hyde) | 158127-100G (paraformaldehyde) |
| Tween-20 | 0.1% PBST/SSCT | Mallinckrodt | — |

**Table S3.** Statistical results. Tortuosity is dimensionless, RMS comparisons are on the raw micrometer scale, and closure-rate differences are in nm/s. Each section identifies the endpoint and statistical comparison. Mutant–control rate tests and direct *twi*/*sna*-versus-*pyr*/*htl* rate tests form separate Holm-adjusted families of four. Post-closure comparisons at 10 and 20 min form a family of two. Raw RMS mutant–control Welch tests are Holm adjusted across four comparisons; the equal-genotype class contrast is reported separately. Recoil rows report *htl*-minus-control plateau (*μ*m) and rate-constant (s^-1^) comparisons, with independent embryos as replicates. A1: seven embryos per genotype, 14 control/12 *htl* tracks; ASPS: three control/six *htl* embryos, four/eight tracks. The four recoil P values are unadjusted. Full numerical results and per-embryo estimates accompany the table.

| Measurement | Comparison | Estimate ± SE | Statistic | d.f.<br>see legend | P |
|---|---|---|---|---|---|
| **A. Peak seam tortuosity** |  |  |  |  |  |
| Seam tortuosity, peak | *pyr* - control | 1.4 ± 0.4 | t = 3.4 | 23 | 0.00897 |
| Seam tortuosity, peak | *htl* - control | 1.6 ± 0.4 | t = 3.93 | 23 | 0.00249 |
| Seam tortuosity, peak | *twi* - control | 1.8 ± 0.4 | t = 4.45 | 23 | 0.000696 |
| Seam tortuosity, peak | *sna* - control | 1.4 ± 0.4 | t = 3.49 | 23 | 0.00727 |
| Seam tortuosity, peak | mean(*twi*, *sna*) − mean(*pyr*, *htl*) | 0.1 ± 0.3 | t = 0.48 | 23 | 0.636 |
| Seam tortuosity, peak | Overall genotype effect | — | F = 5.63 | 4, 23 | 0.0026 |
| **B. Tortuosity after closure** |  |  |  |  |  |
| Mean T at +10/+20 min | mean(*twi*, *sna*) − mean(*pyr*, *htl*) | 0.6 ± 0.2 | t = 3.648 | 11.55 | 0.00355 |
| T at +20 min | mean(*twi*, *sna*) − mean(*pyr*, *htl*) | 0.5 ± 0.2 | t = 3.620 | 23.39 | 0.00174 |
| T at +10 min | mean(*twi*, *sna*) − mean(*pyr*, *htl*) | 0.6 ± 0.2 | t = 3.814 | 23.39 | 0.00174 |
| **C. Closure rates across genotypes** |  |  |  |  |  |
| Closing rate (nm/s) | *pyr* − control | -1 ± 1 | t = -1.136 | 7.899 | 0.2893 |
| Closing rate (nm/s) | *htl* − control | -1.4 ± 0.8 | t = -1.678 | 6.488 | 0.2811 |
| Closing rate (nm/s) | *twi* − control | -3.5 ± 0.8 | t = -4.318 | 6.072 | 0.01457 |
| Closing rate (nm/s) | *sna* − control | -3.9 ± 0.8 | t = -4.929 | 5.61 | 0.01272 |
| **D. Residual dorsal midline deviation** |  |  |  |  |  |
| Midline deviation, raw RMS (*µ*m) | *pyr* − control | 5 ± 1 | t = 4.796 | 4.04 | 0.016899 |
| Midline deviation, raw RMS (*µ*m) | *htl* − control | 2.5 ± 0.6 | t = 3.967 | 5.16 | 0.016899 |
| Midline deviation, raw RMS (*µ*m) | *twi* − control | 18 ± 2 | t = 8.128 | 5.01 | 0.0018086 |
| Midline deviation, raw RMS (*µ*m) | *sna* − control | 21 ± 4 | t = 5.827 | 5.00 | 0.0062884 |
| Midline deviation, raw RMS (*µ*m) | mean(*twi*, *sna*) − mean(*pyr*, *htl*) | 16 ± 2 | t = 7.128 | 9.86 | 3.434e-05 |
| Midline deviation, raw RMS (*µ*m) | Welch overall genotype effect | — | F = 28.44 | 4, 9.515 | 2.749e-05 |
| **E. Recoil after A1 cuts** |  |  |  |  |  |
| A1 recoil, plateau a | *htl* − control | -9 ± 2 | t = -3.82 | 10.8 | 0.00294 |
| A1 recoil, rate k | *htl* − control | 0.05 ± 0.02 | t = 3.05 | 7.16 | 0.01813 |
| **F. Recoil after ASPS cuts** |  |  |  |  |  |
| ASPS recoil, plateau a | *htl* − control | -18 ± 1 | t = -16.7 | 5.11 | 1.181e-05 |
| ASPS recoil, rate k | *htl* − control | 0.06 ± 0.04 | t = 1.66 | 5.14 | 0.1568 |

**Table S4.**
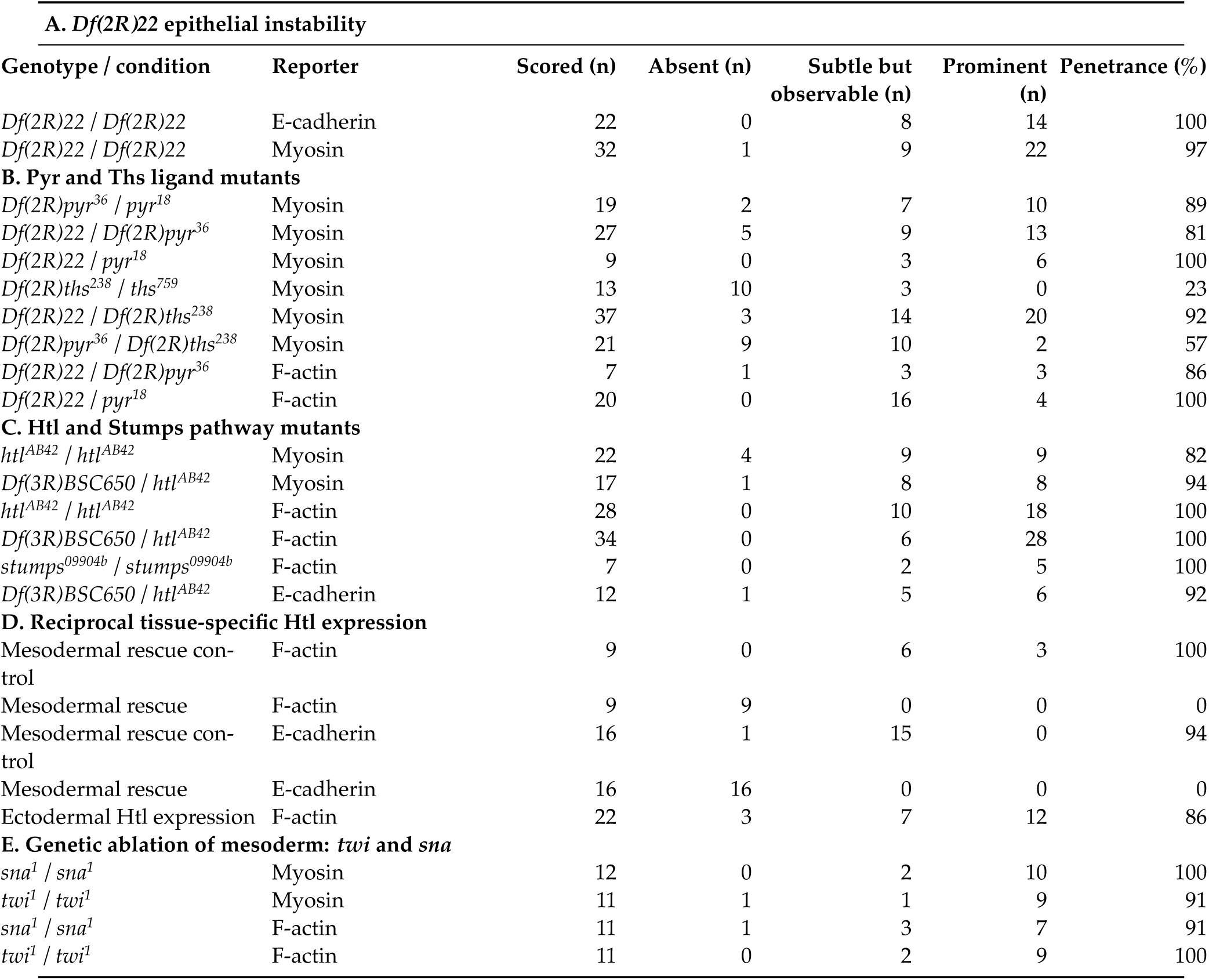
Penetrance and expressivity of epithelial distortions. Penetrance is the percentage of scored embryos with epithelial distortions (subtle but observable plus prominent). Expressivity was scored qualitatively as subtle but observable or prominent. Absent indicates no scored distortion. Categories refer to the scored seam-distortion phenotype; comparisons are made within reporter and genetic backgrounds. Matched mesodermal rescue controls and expression genotypes are defined in Figure 5 and Materials and Methods. Ectodermal Htl expression did not eliminate distortions. Denominators include scorable embryos: two F-actin and four cadherin control embryos were excluded for abnormal closure. All other listed embryos were scorable. Five representative embryos shown in micrographs do not replace the full scored cohorts reported here. Individual embryo scores are provided in the accompanying Table S4 workbook. Sections A–E follow the order of the corresponding Results sections.

| **A. *Df(2R)22* epithelial instability** |  |  |  |  |  |  |
| **Genotype / condition** | **Reporter** | **Scored (n)** | **Absent (n)** | **Subtle but observable (n)** | **Prominent (n)** | **Penetrance (%)** |
|---|---|---|---|---|---|---|
| *Df(2R)22* / *Df(2R)22* | E-cadherin | 22 | 0 | 8 | 14 | 100 |
| *Df(2R)22* / *Df(2R)22* | Myosin | 32 | 1 | 9 | 22 | 97 |
| **B. Pyr and Ths ligand mutants** |  |  |  |  |  |  |
| *Df(2R)pyr<sup>36</sup>* / *pyr<sup>18</sup>* | Myosin | 19 | 2 | 7 | 10 | 89 |
| *Df(2R)22* / *Df(2R)pyr<sup>36</sup>* | Myosin | 27 | 5 | 9 | 13 | 81 |
| *Df(2R)22* / *pyr<sup>18</sup>* | Myosin | 9 | 0 | 3 | 6 | 100 |
| *Df(2R)ths<sup>238</sup>* / *ths<sup>759</sup>* | Myosin | 13 | 10 | 3 | 0 | 23 |
| *Df(2R)22* / *Df(2R)ths<sup>238</sup>* | Myosin | 37 | 3 | 14 | 20 | 92 |
| *Df(2R)pyr<sup>36</sup>* / *Df(2R)ths<sup>238</sup>* | Myosin | 21 | 9 | 10 | 2 | 57 |
| *Df(2R)22* / *Df(2R)pyr<sup>36</sup>* | F-actin | 7 | 1 | 3 | 3 | 86 |
| *Df(2R)22* / *pyr<sup>18</sup>* | F-actin | 20 | 0 | 16 | 4 | 100 |
| **C. Htl and Stumps pathway mutants** |  |  |  |  |  |  |
| *htl<sup>AB42</sup>* / *htl<sup>AB42</sup>* | Myosin | 22 | 4 | 9 | 9 | 82 |
| *Df(3R)BSC650* / *htl<sup>AB42</sup>* | Myosin | 17 | 1 | 8 | 8 | 94 |
| *htl<sup>AB42</sup>* / *htl<sup>AB42</sup>* | F-actin | 28 | 0 | 10 | 18 | 100 |
| *Df(3R)BSC650* / *htl<sup>AB42</sup>* | F-actin | 34 | 0 | 6 | 28 | 100 |
| *stumps<sup>09904b</sup>* / *stumps<sup>09904b</sup>* | F-actin | 7 | 0 | 2 | 5 | 100 |
| *Df(3R)BSC650* / *htl<sup>AB42</sup>* | E-cadherin | 12 | 1 | 5 | 6 | 92 |
| **D. Reciprocal tissue-specific Htl expression** |  |  |  |  |  |  |
| Mesodermal rescue con-trol | F-actin | 9 | 0 | 6 | 3 | 100 |
| Mesodermal rescue | F-actin | 9 | 9 | 0 | 0 | 0 |
| Mesodermal rescue con-trol | E-cadherin | 16 | 1 | 15 | 0 | 94 |
| Mesodermal rescue | E-cadherin | 16 | 16 | 0 | 0 | 0 |
| Ectodermal Htl expression | F-actin | 22 | 3 | 7 | 12 | 86 |
| **E. Genetic ablation of mesoderm: *twi* and *sna*** |  |  |  |  |  |  |
| *sna<sup>1</sup>* / *sna<sup>1</sup>* | Myosin | 12 | 0 | 2 | 10 | 100 |
| *twi<sup>1</sup>* / *twi<sup>1</sup>* | Myosin | 11 | 1 | 1 | 9 | 91 |
| *sna<sup>1</sup>* / *sna<sup>1</sup>* | F-actin | 11 | 1 | 3 | 7 | 91 |
| *twi<sup>1</sup>* / *twi<sup>1</sup>* | F-actin | 11 | 0 | 2 | 9 | 100 |

## Supplementary statistical methods

### Tortuosity and closure progression

Dorsal-opening height and seam tortuosity were measured at the same acquisition frames. The descriptive plots comprise 284 paired measurements from 28 embryos; repeated measurements within an embryo are not independent biological replicates.

The class contrast was [mean(*twi*)+mean(*sna*)]/2 − [mean(*pyr*)+mean(*htl*)]/2, giving each mutant genotype equal weight. Comparisons at 10 and 20 min after closure were adjusted together by Holm’s method.

Closure rates were estimated by ordinary least-squares regression of dorsal-opening height against time, with one slope per embryo (Figs S4, S7, S11). Mean ± s.d. rates were 9 ± 1, 7 ± 2, 7 ± 1, 5.1 ± 1.2 and 5 ± 1 nm/s for control, *pyr*, *htl*, *twi* and *sna*, respectively (n=4 controls; n=6 per mutant). The four mutant–control Welch tests were Holm adjusted together: P=0.289, 0.281, 0.0146 and 0.0127 for *pyr*, *htl*, *twi* and *sna*, respectively. The four direct *twi*/*sna*-versus-*pyr*/*htl* comparisons formed a separate family; adjusted P values were 0.0885 for both comparisons with *pyr*, 0.0439 for *twi* versus *htl* and 0.0198 for *sna* versus *htl*.

### Numerical results for seam tortuosity and midline deviation

Peak seam tortuosity increased in both mesoderm-specification and FGF-pathway mutants, but peak severity did not distinguish the two classes. Tortuosity (T) measures contour length relative to the straight distance between the same endpoints; T = 1 denotes a straight seam. Peak T differed among genotypes (ANOVA, F(4, 23) = 5.63, P = 0.0026), with each mutant exceeding control in Dunnett-type comparisons (*pyr*, P = 0.0090; *htl*, P = 0.0025; *twi*, P = 0.0007; *sna*, P = 0.0073). However, the mean peak difference between the *twi / sna* and *pyr / htl* classes was small and not significant (0.1 ± 0.3 s.e.; t = 0.48, d.f. = 23, P = 0.636; Table S3). Embryo peaks and post-closure measurements are shown in FIG. S12.

Residual RMS deviation remained greater in *twi* and *sna* after local zigzags largely subsided (Fig. 7M). On the raw micrometer scale, *twi* exceeded control by 18.12 *μ*m (Holm-adjusted P = 0.00181) and *sna* by 20.68 *μ*m (Holm-adjusted P = 0.00629). The equally weighted *twi / sna* genotype mean exceeded *pyr / htl* by 15.57 ± 2.18 *μ*m s.e. (95% confidence interval 10.70–20.45 *μ*m; t = 7.13, d.f. = 9.86, P = 0.0000343; Table S3). Signed tracks are descriptive repeated observations, not independent replicates (FIG. S13).

## Supplementary movie legends

**Movie 1. Dorsal closure in an E-cadherin-labelled control embryo.** E-cadherin-GFP marks epithelial junctions during dorsal closure. Time is shown in minutes. Scale bar, 20 µm.

**Movie 2. Late amnioserosa defects in a *Df(2R)22* embryo.** A *Df(2R)22* homozygous embryo expressing E-cadherin-GFP develops PAS–DME separation, junctional discontinuity and amnioserosa disintegration during later dorsal closure. Time is shown in minutes. Scale bar, 20 µm.

**Movie 3. Early amnioserosa disintegration in a *Df(2R)22* embryo.** E-cadherin-GFP reveals extensive amnioserosa disruption and cellular debris in a *Df(2R)22* homozygous embryo. Time is shown in minutes. Scale bar, 20 µm.

**Movie 4. Dorsal closure in a myosin-labelled control embryo.** Sqh-GFP-RLC reveals the advancing purse strings and comparatively straight formed dorsal seam. Time is shown in minutes. Scale bar, 20 µm.

**Movie 5. Dynamic epithelial distortions in a *Df(2R)22* embryo.** Sqh-GFP-RLC reveals changing distortions of the advancing edge and formed dorsal epithelium in a *Df(2R)22* homozygous embryo. Time is shown in minutes. Scale bar, 20 µm.

**Movie 6. Epithelial distortions in a *pyr* mutant embryo.** A *Df(2R)pyr^36^* / *Df(2R)22* embryo expressing Sqh-GFP-RLC develops dynamic epithelial distortions during dorsal closure. Time is shown in minutes. Scale bar, 20 µm.

**Movie 7. Epithelial distortions in a *ths* mutant embryo.** A *Df(2R)ths^238^* / *Df(2R)22* embryo expressing Sqh-GFP-RLC develops epithelial distortions during dorsal closure. Time is shown in minutes. Scale bar, 20 µm.

**Movie 8. Dorsal closure in an F-actin-labelled control embryo.** The sGMCA reporter reveals epithelial organization during dorsal closure. Time is shown in minutes. Scale bar, 20 µm.

**Movie 9. Epithelial distortions in an *htl* mutant embryo.** An *htl^AB42^* / *Df(3R)BSC650* embryo expressing sGMCA develops changing epithelial distortions. Anterior and posterior image fieelds are stitched to display the closing epithelium. Time is shown in minutes. Scale bar, 20 µm.

**Movie 10. Epithelial distortions in a *stumps* mutant embryo.** An sGMCA-labelled *stumps^09^ ^904b^* homozygous embryo develops epithelial distortions during dorsal closure. Time is shown in minutes. Scale bar, 20 µm.

**Movie 11. Junctional abnormalities in an *htl* mutant embryo.** E-cadherin-GFP reveals altered epithelial associations and distortions in an *htl^AB42^* / *Df(3R)BSC650* embryo. Frames with inadequate focus are omitted; displayed elapsed times retain the acquisition timing. Time is shown in minutes. Scale bar, 20 µm.

**Movie 12. Htl localization beneath the closing epithelium.** Htl-mCherry is green and E-cadherin-GFP is magenta. Time is calculated from acquisition timestamps; the median interval between time points is approximately 4.3 min. Scale bar, 20 µm.

**Movie 13. Epithelial distortions in matched *htl* mutant controls.** Two *htl* mutant embryos carrying *twi-GAL4* without a *UAS-htl* responder show epithelial distortions; one also exhibits defective germband retraction. Time is shown in minutes. Scale bar, 20 µm.

**Movie 14. Mesodermal expression of *htl* suppresses epithelial distortions.** Two *twi-GAL4* > *UAS-htl^WT^*; *htl^AB42^* / *htl^AB42^* embryos illustrate the comparatively straight dorsal seam following mesodermal rescue. Time is shown in minutes. Scale bar, 20 µm.

**Movie 15. Ectodermal expression of *htl* does not suppress epithelial distortions.** A *69B-GAL4* > *UAS-htl^WT^*; *htl^AB42^* / *htl^AB42^* embryo retains epithelial distortions during dorsal closure. Time is shown in minutes. Scale bar, 20 µm.

**Movie 16. Mesodermal organization beneath wild-type ectoderm.** *twi-GAL4*-driven *UAS-RFP-Moesin* marks mesoderm in green; E-cadherin-GFP marks ectodermal junctions in magenta. The median acquisition interval is approximately 3.1 min.

**Movie 17. Mesodermal disorganization beneath *htl* mutant ectoderm.** Two *htl* mutant embryos are shown sequentially. *twi-GAL4*-driven *UAS-RFP-Moesin* marks mesoderm in green, and E-cadherin-GFP marks ectodermal junctions in magenta. Images were acquired approximately every 2 min. Time is shown separately for each example. Two time points acquired with an incorrect mesoderm-channel fielter are omitted from the second example. Scale bars, 20 µm.

**Movie 18. Persistent epithelial distortions in a *sna* mutant embryo.** Sqh-GFP-RLC reveals changing epithelial distortions and a tortuous seam in a *sna^1^* homozygous embryo. Time is shown in minutes. Scale bar, 20 µm.

**Movie 19. Persistent epithelial distortions in *twi* mutant embryos.** Two *twi^1^* homozygous embryos expressing Sqh-GFP-RLC illustrate epithelial distortions during dorsal closure and residual seam tortuosity. Time is shown in minutes. Scale bar, 20 µm. The two embryos are displayed side by side.

**Movie 20. Recoil following A1 laser incisions.** Control, *pyr* and *htl* embryos are shown from left to right. Canthus-to-canthus incisions bisect the amnioserosa along the dorsal midline. Elapsed acquisition time is displayed separately for each embryo. Images were acquired approximately every 0.2 s. Scale bars, 20 µm.

**Movie 21. Recoil following ASPS laser incisions.** Control, *pyr* and *htl* embryos are shown from left to right. Transverse incisions cross the amnioserosa, purse strings and adjacent lateral epidermis. Elapsed acquisition time is displayed separately for each embryo. Images were acquired approximately every 0.2 s. Scale bars, 20 µm.

**Movie 22. Local recoil following short laser incisions near epithelial distortions.** Three local incisions in *pyr* mutant embryos illustrate variation in recoil near individual distortion peaks. Images were acquired every 90 ms. Elapsed acquisition time is shown in minutes and seconds. Scale bars, 20 µm.

**Movie 23. Epithelial distortions in a chromosome 3R deficiency embryo.** A *Df(3R)BSC141* homozygous embryo expressing Sqh-GFP-RLC develops epithelial distortions during dorsal closure. Time is shown in minutes. Scale bar, 20 µm.

**Movie 24. Dorsal closure in a *Pi3K92E* trans-heterozygous embryo.** A *Pi3K92E* / *Df(3R)BSC141* embryo expressing Sqh-GFP-RLC illustrates the epithelial phenotype. Time is shown in minutes. Scale bar, 20 µm.

**Movie 25. Cortical configurations following reduced *pbl* dosage in an *htl* background.** Three *pbl^3^*, *htl^AB42^*/ *htl^AB42^* embryos expressing the F-actin reporter illustrate pull-like and protrusive epithelial confiegurations. Anterior and posterior fieelds are stitched for each embryo. Frame numbers are shown. Scale bars, 20 µm.

